# Translatome analysis reveals cellular network in DLK-dependent hippocampal glutamatergic neuron degeneration

**DOI:** 10.1101/2024.07.10.602846

**Authors:** Erin M Ritchie, Dilan Acar, Siming Zhong, Qianyi Pu, Yunbo Li, Binhai Zheng, Yishi Jin

**Affiliations:** Department of Neurobiology, School of Biological Sciences, University of California San Diego, La Jolla, CA 92093; Biomedical Sciences Graduate Program, School of Medicine, University of California San Diego, La Jolla, CA 92093; Department of Neurosciences, School of Medicine, University of California San Diego, La Jolla, CA 92093; Kavli Institute of Brain and Mind, University of California San Diego, La Jolla, CA 92093

**Keywords:** microtubules, Jun, stathmin, neurite differentiation, synapse formation, RiboTag-seq

## Abstract

The conserved MAP3K12/Dual Leucine Zipper Kinase (DLK) plays versatile roles in neuronal development, axon injury and stress responses, and neurodegeneration, depending on cell-type and cellular contexts. Emerging evidence implicates abnormal DLK signaling in several neurodegenerative diseases. However, our understanding of the DLK-dependent gene network in the central nervous system remains limited. Here, we investigated the roles of DLK in hippocampal glutamatergic neurons using conditional knockout and induced overexpression mice. We found that dorsal CA1 and dentate gyrus neurons are vulnerable to elevated expression of DLK, while CA3 neurons appear less vulnerable. We identified the DLK-dependent translatome that includes conserved molecular signatures and displays cell-type specificity. Increasing DLK signaling is associated with disruptions to microtubules, potentially involving STMN4. Additionally, primary cultured hippocampal neurons expressing different levels of DLK show altered neurite outgrowth, axon specification, and synapse formation. The identification of translational targets of DLK in hippocampal glutamatergic neurons has relevance to our understanding of selective neuron vulnerability under stress and pathological conditions.

## Introduction

The mammalian Mitogen Activated Protein Kinase Kinase Kinase (MAP3K12) Dual Leucine Zipper Kinase (DLK) is broadly expressed in the nervous system from early development to mature adults. DLK exerts its effects primarily through signal transduction cascades involving downstream MAP2Ks (MKK4, MKK7) and MAPKs (JNK, p38, ERK), which then phosphorylate many proteins, such as transcription factors including c-Jun, to regulate cellular responses (Asghari Adib et al., 2018; Hirai et al., 2006; Huang et al., 2017; Itoh et al., 2009; Jin & Zheng, 2019; Tedeschi & Bradke, 2013). In cultured neurons, DLK is localized to axons (Hirai et al., 2005; Lewcock et al., 2007), dendrites (Pozniak et al., 2013), and the Golgi apparatus (Hirai et al., 2002). DLK is also associated with transporting vesicles, which are considered platforms for DLK to serve as a sensor of neuronal stress or injury (Holland et al., 2016; Tortosa et al., 2022). Despite broad expression, functional investigations of DLK have been limited to a few cell types under specific conditions.

Constitutive DLK knockout (KO) mice, generated by removing the N-terminus of DLK, including the ATP binding motif of the kinase domain (Hirai et al., 2006), or by deleting the entire kinase domain (Ghosh et al., 2011), die perinatally. The development of the embryonic nervous system is largely normal, with mild defects in radial migration and axon track formation in the developing cortex (Hirai et al., 2006) and neuronal apoptosis during development of spinal motor neurons and dorsal root ganglion (DRG) neurons (Ghosh et al., 2011; Itoh et al., 2011). Selective removal of DLK in layer 2/3 cortical neurons starting at E16.5 results in increased dendritic spine volume (Pozniak et al., 2013). Induced deletion of DLK in adult mice causes no obvious brain structural defects; synapse size and density in hippocampus and cortex appear unaltered, although basal synaptic strength is mildly increased (Pozniak et al., 2013). In contrast, under injury or stress conditions, DLK exhibits critical context-specific roles. In DRG neurons, DLK is required for nerve growth factor withdrawal induced death, promotes neurite regrowth, and is also involved in retrograde injury signaling (Ghosh et al., 2011; Holland et al., 2016; Itoh et al., 2009; Shin et al., 2012). In a spinal cord injury model, DLK is required for *Pten* deletion*-*induced axon regeneration and sprouting as well as spontaneous sprouting of uninjured corticospinal tract neurons (Saikia et al., 2022). In optic nerve crush assay, DLK is necessary for *Pten* deletion*-*induced axon regeneration of retinal ganglion cells (RGC), but also contributes to injury-induced RGC death (Watkins et al., 2013). In a mouse model of stroke, increased DLK expression is associated with motor recovery following knockdown of the CCR5 chemokine receptor (Joy et al., 2019). These studies reveal critical roles of DLK in development, maintenance, and repair of neuronal circuits.

DLK is known to be expressed in hippocampal neurons (Blouin, 1996; Hirai et al., 2005; Mata et al., 1996). Loss of DLK, either constitutively or in adult animals, causes no discernable effect on hippocampal morphology (Hirai et al., 2006; Pozniak et al., 2013). Microarray-based gene expression analysis did not detect significant changes associated with loss of DLK in the hippocampus (Pozniak et al., 2013). However, following exposure to kainic acid, loss of DLK, or preventing phosphorylation of the downstream transcription factor c-Jun, significantly reduces neuron death in hippocampus (Behrens et al., 1999; Pozniak et al., 2013). Additionally, elevated levels of p-c-Jun are observed in hippocampus of patients with Alzheimer’s disease (Le Pichon et al., 2017). Induced human neurons treated with ApoE4, a prevalent ApoE variant associated with Alzheimer’s disease, also show upregulation of DLK, which leads to enhanced transcription of APP and thus Aβ levels (Huang et al., 2017). These data suggest that transcriptional changes downstream of DLK may be an important aspect of its signaling in hippocampal neuron degeneration under pathological conditions.

Here, we investigate the DLK-dependent molecular and cellular network in hippocampal glutamatergic neurons, which show selective vulnerability in Alzheimer’s disease, ischemic stroke, and excitotoxic injury. Using DLK conditional knockout and overexpression mice, we reveal hippocampal regional differences in neuronal death upon elevated DLK signaling. We describe translational changes in hippocampal glutamatergic neurons using RiboTag-seq analysis. We show that the key transcription factor c-Jun and a member of the stathmin family, STMN4, display DLK-dependent translation. Our analyses on hippocampal tissues and cultured neurons support the conclusion that the DLK-dependent signaling network has important roles in the regulation of microtubule homeostasis, neuritogenesis, and synapse formation.

## Results

### DLK conditional knockout in differentiating and mature glutamatergic neurons does not alter gross morphology of hippocampus

As a first step to define the roles of DLK in hippocampal glutamatergic neurons, we verified DLK expression in hippocampal tissue by RNAscope analysis. We observed strong signals in the glutamatergic pyramidal cells and granule cells in P15 mice (Fig.S1A), consistent with prior in situ data (Blouin, 1996; Lein et al., 2007; Mata et al., 1996). To selectively delete DLK in glutamatergic neurons, we generated *Vglut1^Cre/+^;DLK(cKO)^fl/fl^* mice. *DLK(cKO)^fl/fl^* have LoxP sites flanking the exon encoding the initiation ATG and the first 149 amino acids (Fig.S1C) (Chen et al., 2016; Li et al., 2021; Saikia et al., 2022). *Vglut1^Cre^*expression in hippocampus shows strong expression in CA3 and in a subset of pyramidal neurons close to stratum oriens in CA1 at P4, with broad expression in both CA1 and CA3 by P14 (Harris et al., 2014). In dentate gyrus, expression of *Vglut1^Cre^* begins in neurons nearer the molecular layer around P4, with expression spreading towards the polymorph layer gradually during the first two postnatal months. By western blot analysis of hippocampal protein extracts we found that full-length DLK protein was significantly reduced in *Vglut1^Cre/+^;DLK(cKO)^fl/fl^*(Fig.1A,B). The DLK antibody also detected a protein product of lower molecular weight at much reduced levels (Fig.S1B). As *Dlk* mRNA lacking the floxed exon was expressed (Fig.S1C), this lower-molecular weight protein could be produced by using a downstream alternative start codon (Fig.S1C1), but would lack the N-terminal palmitoylation motif and ATP-binding site that are essential for DLK activity (Holland et al., 2016; Huntwork-Rodriguez et al., 2013). These data provide validation for knockout of functional DLK protein in hippocampal glutamatergic neurons in *Vglut1^Cre/+^;DLK(cKO)^fl/fl^* mice.

**Figure 1.**
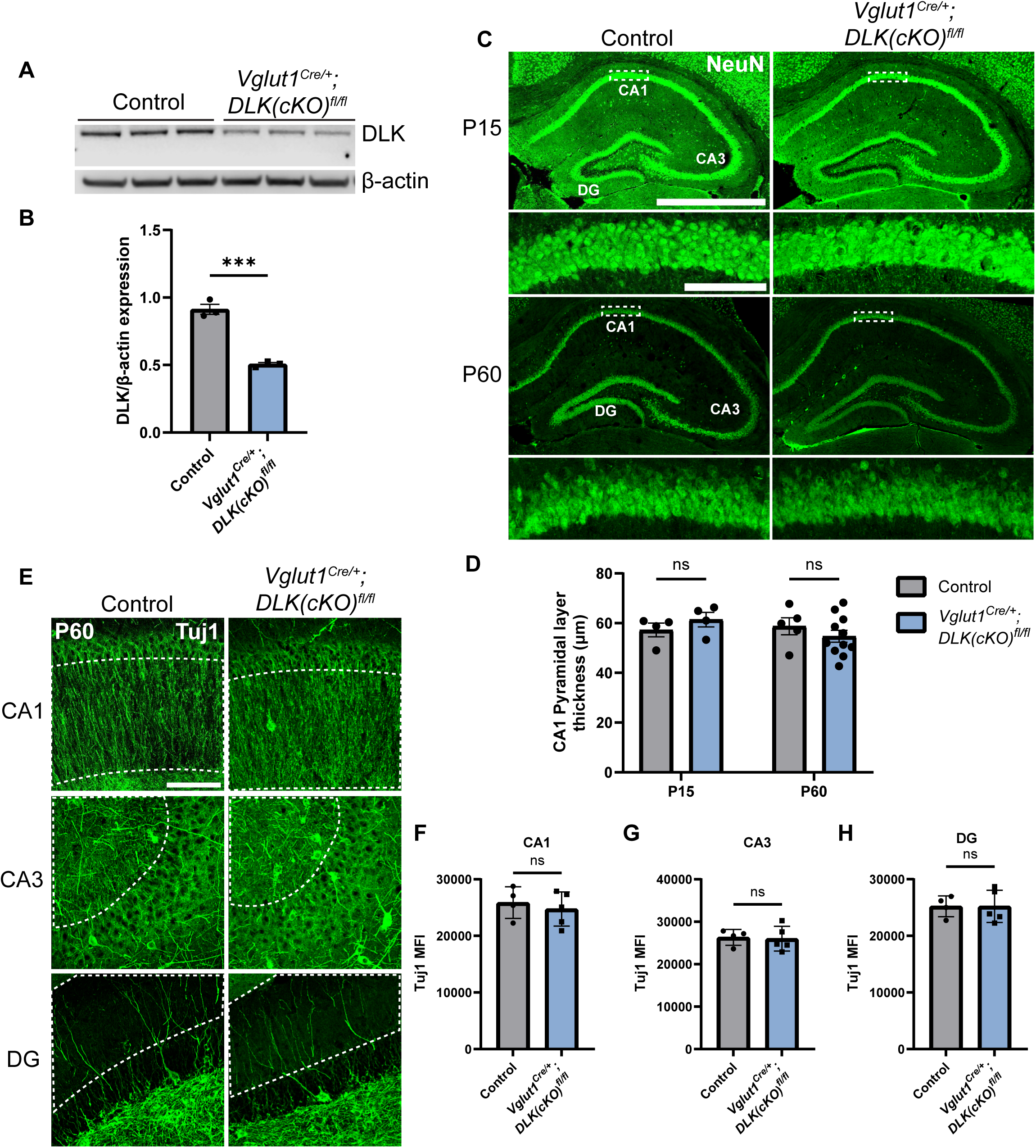
Deletion of DLK in postmitotic glutamatergic neurons does not alter gross morphology of hippocampus. (A) Western blot of DLK and β-actin in protein extracts of hippocampal tissue of *Vglut1^Cre/+^;DLK(cKO)^fl/fl^* and *DLK(cKO)^fl/fl^*littermate controls (age P60, each lane representing individual mice, N=3 mice/genotype). (B) Quantification of DLK protein level normalized to β-actin. Statistics: Unpaired t-test, *** P<0.001. (C) Confocal z-stack (max projection) images of NeuN immunostaining of coronal sections of the dorsal hippocampus in P15 and P60 mice of genotype indicated, respectively. Dashed boxes in CA1 pyramidal layers are enlarged below. Scale bar, 1,000 μm in hippocampi; 100 μm in CA1 layer. (D) Quantification of CA1 pyramidal layer thickness. Each dot represents averaged thickness from 3 sections per mouse; N≥4 mice/genotype per timepoint. Statistics: Two-way ANOVA with Holm-Sidak multiple comparison test; ns, not significant. (E) Confocal z-stack (max projection) images of Tuj1 immunostaining of hippocampus CA1, CA3, and DG regions in control and *Vglut1^Cre/+^;DLK(cKO)^fl/fl^* mice (age P60). Dashed outlines mark ROI (region of interest) for fluorescence intensity quantification. Scale bar, 100μm. (F,G,H) Tuj1 mean fluorescence intensity (MFI) after thresholding signals in dendritic regions in each hippocampal area. Each dot represents averaged intensity from 3 sections per mouse; N=4 control, 5 *Vglut1^Cre/+^;DLK(cKO)^fl/fl^*. Statistics: Unpaired t-test. ns, not significant.

The *Vglut1^Cre/+^;DLK(cKO)^fl/fl^* mice were indistinguishable from control littermate mice in behavior and appearance from birth to about one year of age. We examined tissue sections of hippocampus in P15 and P60 mice. Hippocampal sections stained with NeuN, a marker of neuronal nuclei (Mullen et al., 1992), showed no significant difference in overall position of neuronal soma or thickness of the CA1 pyramidal cell layer at either timepoint (Fig.1C,D). Neuronal morphology visualized by immunostaining with Tuj1, labeling neuron-specific β-III tubulin, also showed no detectable differences in the pattern and intensity of microtubules in *Vglut1^Cre/+^;DLK(cKO)^fl/fl^* mice, compared to control (Fig.1E-H). Gross morphology of hippocampus and surrounding tissues in 1 year old *Vglut1^Cre/+^;DLK(cKO)^fl/fl^*mice was indistinguishable from controls (Fig.S1E, S1G-I, S2A). These results show that DLK does not have essential roles in post-mitotic hippocampal glutamatergic neuron maintenance.

### Increasing expression levels of DLK leads to hippocampal neuron death, with dorsal CA1 neurons showing selective vulnerability

Several studies have reported that DLK protein levels increase under a variety of conditions, including optic nerve crush (Watkins et al., 2013), NGF withdrawal (∼2 fold) (Huntwork-Rodriguez et al., 2013; Larhammar et al., 2017), and sciatic nerve injury (Larhammar et al., 2017). Induced human neurons show increased DLK abundance about ∼4 fold in response to ApoE4 treatment (Huang et al., 2019). Increased expression of DLK can lead to its activation through dimerization and autophosphorylation (Nihalani et al., 2000). We thus asked how increased DLK signaling affects hippocampal glutamatergic neurons. We previously described a transgenic mouse, *H11-DLK^iOE/+^*, which allows Cre-dependent DLK overexpression (Li et al., 2021). The DLK transgene is coexpressed with tdTomato through a T2A peptide (Fig. S1D). By RNAscope analysis we observed increased *Dlk* mRNAs in glutamatergic neurons in CA1, CA3, and DG in *Vglut1 ^Cre/+^;H11-DLK^iOE/+^* mice at P15, compared to control mice (Fig.S3A,B). By immunostaining hippocampal sections with anti-DLK antibodies we observed increased protein levels particularly in regions with pyramidal neuron dendrites in *Vglut1 ^Cre/+^;H11-DLK^iOE/+^*, compared to control mice (Fig.S3C-E). Additional analysis at the mRNA level (supplemental excel, File S2. WT vs DLK(iOE) DEGs) and at the protein level (Fig.S8E) suggest that the increase in DLK abundance was around 3 times the control level. The localization patterns of DLK protein appeared to vary depending on region of hippocampus and age of animals in both control and *Vglut1 ^Cre/+^;H11-DLK^iOE/+^* mice (Fig.S3C).

*Vglut1 ^Cre/+^;H11-DLK^iOE/+^* mice were born normally, and developed noticeable progressive motor deficits around four months of age, which became worse by one year of age. We stained brain sections for NeuN at P10, P15, P60, and 1 year of age and observed a progressive reduction in brain size of these mice, compared to controls (Fig.S2B,C). At P10, the dorsal hippocampus in *Vglut1 ^Cre/+^;H11-DLK^iOE/+^*was indistinguishable from control (Fig.2A, 2B, S2B). By P15, the DLK(iOE) mice showed significant thinning of the CA1 pyramidal layer. We detected increased TUNEL staining signals in CA1 pyramidal layer, compared to control (Fig.S5F S5G). By P60, most CA1 pyramidal neurons were lost, while DG began to show thinning, which continued to worsen at 1 year of age (Fig.2A, 2B, S1F, S1I, S2B, S2C). In contrast, neurons in CA3 appeared less affected, even at 1 year of age (Fig.2A, S1F, S1H, S2C). Additionally, in P60, dorsal CA1 showed significantly fewer surviving neurons, while ventral CA1 pyramidal layer thickness appeared more similar to control than dorsal regions (Fig.S4A,B). Neuronal death generally induces reactive astrogliosis. We stained for GFAP, a marker of astrocyte reactivity. We found increased GFAP staining in *Vglut1 ^Cre/+^;H11-DLK^iOE/+^,* specifically in CA1 at P15, and at P60 in CA1 and DG, but not as strongly in CA3, compared to control mice (Fig.S5A-D). We also stained for IBA1, a marker of microglia, and found that *Vglut1 ^Cre/+^;H11-DLK^iOE/+^* mice showed increased IBA1 staining around the CA1 region, compared to control mice (Fig.S5E). Microglia appeared ramified in control mice and more reactive-looking in *Vglut1 ^Cre/+^;H11-DLK^iOE/+^* mice (Fig.S5E). Together, these data reveal that dorsal CA1 neurons show vulnerability to elevated DLK expression, while CA3 neurons appear less vulnerable to DLK overexpression.

**Figure 2.**
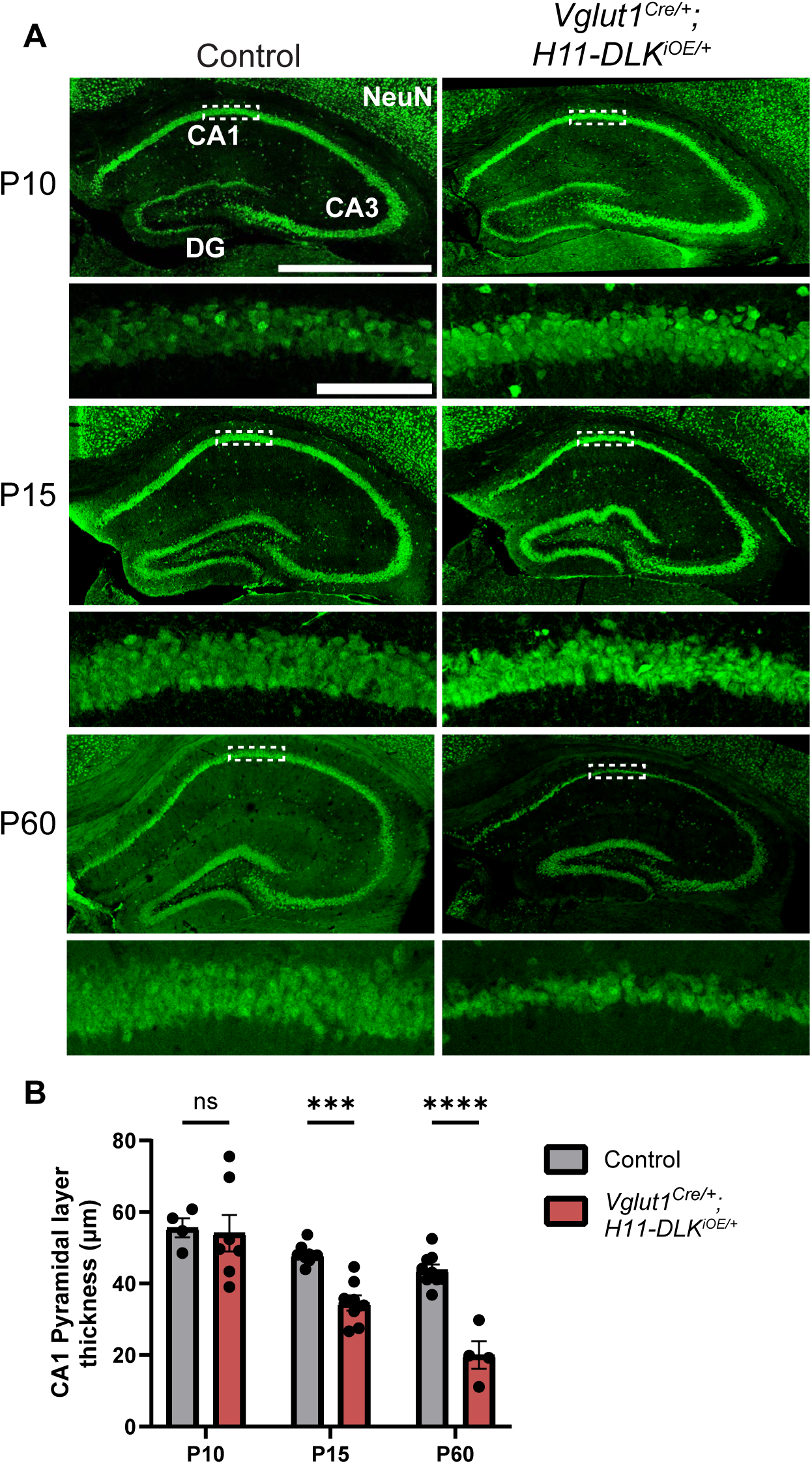
Induced DLK overexpression in hippocampal glutamatergic neurons causes degeneration of CA1 neurons. (A) Confocal z-stack (max projection) images of NeuN immunostaining of coronal sections from dorsal hippocampus in P10, P15, and P60 mice of genotype indicated. Dashed boxes mark CA1 pyramidal layers enlarged below. P60 images shown under different settings compared to P10 and P15 due to older staining. Scale bar, 1,000μm in hippocampi; 100μm in CA1 layer. (B) Quantification of CA1 pyramidal layer thickness. Data points represent averaged measurement from 3 sections per mouse, N≥4 mice/genotype at each timepoint. Statistics: Two-way ANOVA with Holm-Sidak multiple comparison test. ns, not significant; *** P<0.001; **** P<0.0001.

### DLK dependent translated genes are enriched in synapse formation and function

To gain understanding of molecular changes associated with DLK expression levels in glutamatergic neurons, we next conducted translating ribosome profiling and RNA sequencing (RiboTag profiling) using *Rpl22^HA^*mice, which enables Cre-dependent expression of an HA tagged RPL22, a component of the ribosome, from its endogenous locus (Sanz et al., 2009). We generated *Vglut1 ^Cre/+^;H11-DLK^iOE/+^;Rpl22^HA/+^*, *Vglut1^Cre/+^;DLK(cKO)^fl/fl^;Rpl22^HA/+^*, and their respective *Vglut1^Cre/+^;Rpl22^HA/+^* sibling controls. We made protein extracts from dissected hippocampi of P15 mice, a time point when some CA1 neuron degeneration induced by DLK overexpression was visible. We obtained affinity purified HA-immunoprecipitates with the associated actively translated RNAs (Fig.S6A) (n=3 DLK(iOE)/3 WT, n=4 DLK(cKO)/4 WT) and verified purity of the isolated RNA samples by qRT-PCR (Fig.S6B). We mapped >24 million deep sequencing reads per sample to approximately 14,000 genes. We found 260 genes that were differentially expressed and translated in DLK(iOE) neurons, including 114 up- and 146 down-regulated genes, compared to control (Fig.3A, using the cutoff of p_adj_<0.05, File S2 WT vs DLK(iOE) DEGs). 36 genes showed significant changes in DLK(cKO) neurons, including 12 up- and 24 down-regulated genes, compared to control (Fig.3B, p_adj_<0.05, File S1 WT vs DLK(cKO) DEGs). Among genes with statistically significant changes, 17 were detected in both DLK(cKO) and DLK(iOE) (Fig.S6C), of which 13 were upregulated in DLK(iOE) and downregulated in DLK(cKO), and 3 were downregulated in DLK(iOE) and upregulated in DLK(cKO) (Fig.S6D, S6E). The most significant differentially expressed genes included *Jun,* encoding the DLK downstream transcription factor c-Jun, *Stmn4,* encoding a member of the Stathmin tubulin-binding protein family, and *Sh2d3c*, encoding a SH2-domain cytoplasmic signaling protein (Dodelet et al., 1999; Vervoort et al., 2007). One gene, *Slc25a17*, a peroxisomal transporter for cofactors FAD, CoA, and others (Agrimi et al., 2012) and broadly implicated in oxidative stress, was upregulated in both DLK(cKO) and DLK(iOE), compared to control, though the relevance of this change may require further investigation. To systematically compare whether DLK regulates the translatome in a coordinated manner, we performed rank-rank hypergeometric overlap (RRHO) analysis (Plaisier et al., 2010) on the entire translated mRNAs detected in DLK(iOE) and DLK(cKO). We found that RRHO detected significant overlap in genes that were upregulated in DLK(iOE) and downregulated in DLK(cKO) as well as the reverse (Fig.3C), supporting a conclusion that expression of many of the same genes are dependent on DLK.

**Figure 3:**
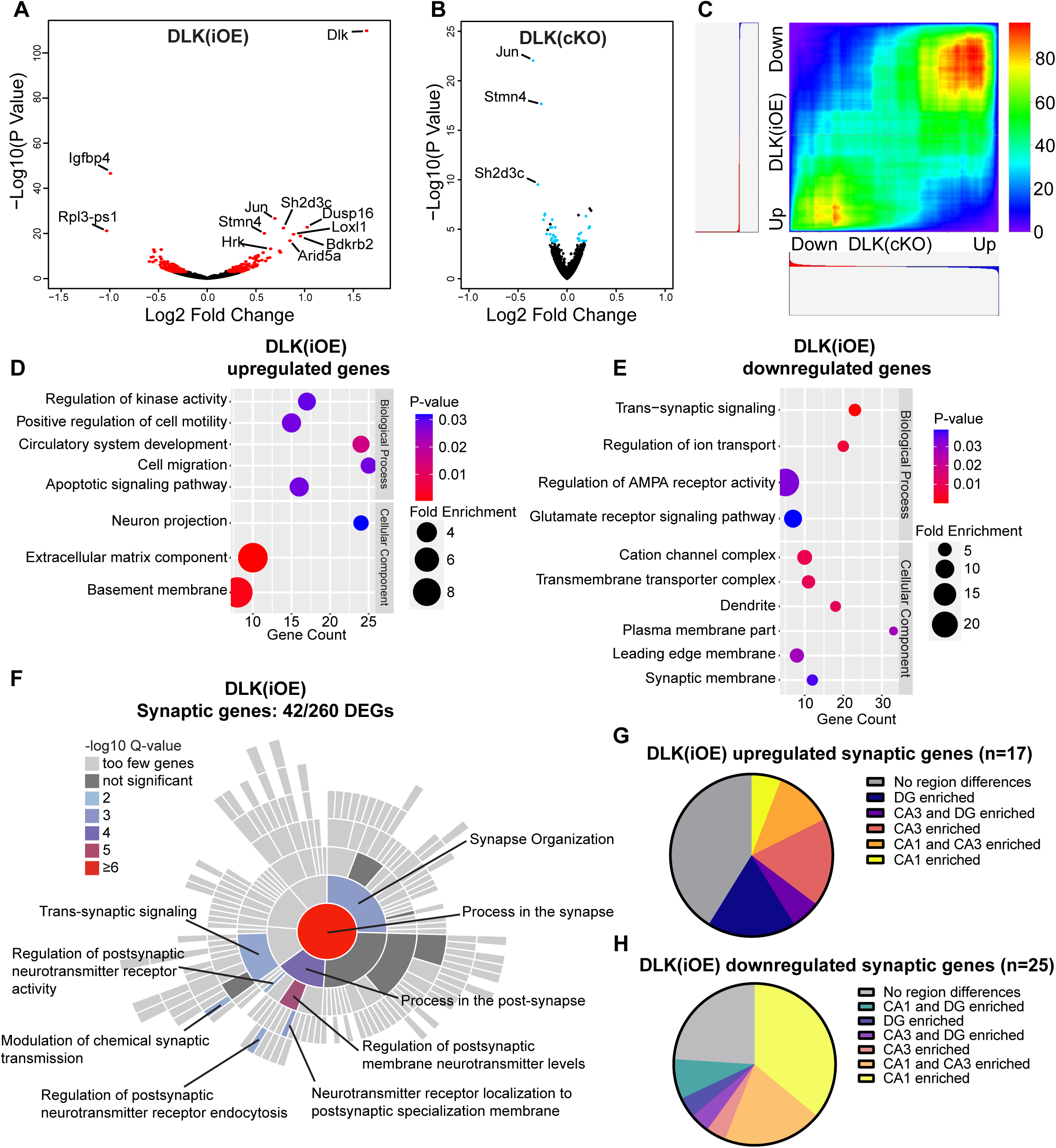
Differentially expressed genes revealed by RiboTag analysis of hippocampal glutamatergic neurons in DLK(cKO) and DLK(iOE) mice. (A) Volcano plot showing RiboTag analysis in *Vglut1^Cre/+;^H11-DLK^iOE/+^;Rpl22^HA/+^*vs *Vglut1^Cre/+^;Rpl22^HA/+^* (age P15). 260 genes (red) show differential expression with adjusted p-values < 0.05 in *Vglut1^Cre/+;^H11-DLK^iOE/+^*, compared to control; names of genes with p<1E-10 are labeled. (B) Volcano plot showing RiboTag analysis in *Vglut1^Cre/+^;DLK(cKO)^fl/fl^;Rpl22^HA/+^*vs *Vglut1^Cre/+^;Rpl22^HA/+^* (age P15). 36 genes (blue) show differential expression with adjusted p-values < 0.05; names of genes with p<1E-10 are labeled. (C) Rank-rank hypergeometric overlap (RRHO) comparison of gene expression in DLK(cKO) and DLK(iOE) RiboTag datasets shows enrichment of similar genes when DLK is low or high, respectively. Color represents the -log transformed hypergeometric p-values (blue for weaker p-value, red for stronger p-value). (D,E) Gene ontology (GO) analysis of significantly up- or down-regulated genes in hippocampal glutamatergic neurons of DLK(iOE) mice compared to the control. Colors correspond to P-values; circle size represents fold enrichment for the GO term; X position shows # of genes significantly enriched in the GO term. (F) SynGO sunburst plot shows enrichment of 42 differentially expressed genes from hippocampal glutamatergic neurons of DLK(iOE) mice, with color corresponding to significance. (G,H) Pie charts show distribution of the 42 synaptic genes up- or down-regulated in DLK(iOE), respectively, in CA1, CA3, and DG in dorsal hippocampus, based on *in situ* data (P56) in the Allen Mouse Brain Atlas.

To gain understanding of DLK-dependent signaling network, we performed gene ontology (GO) analysis on the 260 genes differentially translated in DLK(iOE) neurons, as this dataset gave greater ability to detect significant GO terms than using the 36 genes differentially expressed in DLK(cKO). The genes upregulated in DLK(iOE) (114) had enrichment in GO terms related to neuron apoptosis, cell migration, cell adhesion, and the extracellular matrix organization (Fig.3D), whereas the genes downregulated (146) had GO terms related to synaptic communication and ion transport (Fig.3E). Similar GO terms were also identified using the list of genes coordinately regulated by DLK, derived from our RRHO analysis. Among the genes upregulated in DLK(iOE), some were known to be involved in neurite outgrowth (*Plat*, *Tspan7, Hap1*), endocytosis or endosomal trafficking (*Snx16, Ston2, Hap1*), whereas the genes down-regulated in DLK(iOE) included ion channel subunits (*Cacng8, Cacng3, Grin2b, Scn1a*) and those in exocytosis and calcium related proteins (*Doc2b, Hpca, Cadps2, Rab3c, Rph3a*). A significant cluster of differentially expressed genes in DLK(iOE) included those that regulate AMPA receptors (*Nptx1, Nptxr, Cnih3, Gpc4, Arc, Tspan7)* and cell adhesion molecules (*Nectin1, Flrt3, Pcdh8, Plxnd1*). A further survey using SynGO, a curated resource for genes related to synapse formation and function (Koopmans et al., 2019), revealed 42 of 260 differentially expressed genes in DLK(iOE) showed significant enrichment in synaptic organization and postsynaptic receptor signaling processes (Fig.3F). Conversely, 10 of the 36 differentially expressed genes in DLK(cKO) were annotated to function in similar synaptic processes as in DLK(iOE) (Fig.S6J). The bioinformatic analysis suggests that increased DLK expression can promote translation of genes related to neurite outgrowth and branching and reduce those related to the maturation and function of synapses.

The hippocampus is comprised of multiple glutamatergic neuron types with distinct spatial patterns of gene expression (Lein et al., 2004). As we observed regional vulnerability to DLK overexpression, we next asked if the differentially expressed genes associated with DLK(iOE) might show correlation to the neuronal vulnerability. We first surveyed the endogenous expression pattern of the 260 significantly changed genes in DLK(iOE) in hippocampus using in situ data from 8-week-old mice from the Allen Brain Atlas (Lein et al., 2007). We found that about a third of the genes downregulated in DLK(iOE) showed enriched expression in CA1 (Fig.S6G), and some of these genes, including *Tenm3, Lamp5,* and *Mpped1,* were up-regulated in DLK(cKO) (Fig.S6H, S6I). In comparison, about 50% of the genes upregulated in DLK(iOE) showed comparable expression among hippocampal cell types (Fig.S6F). Additionally, among the 42 synaptic genes that were differentially expressed in DLK(iOE), a notable portion of the downregulated genes showed enriched expression in CA1 (Fig.3H), while the upregulated genes were expressed in all regions (Fig.3G).

Additionally, we compared our Vglut1-RiboTag datasets with CamK2-RiboTag and Grik4-RiboTag datasets from 6-week-old wild type mice reported by (Traunmüller et al., 2023; GSE209870). We defined a list of genes enriched in CamK2-expressing CA1 neurons relative to Grik4-expressing CA3 neurons (CA1 genes), and those enriched in Grik4-expressing CA3 neurons (CA3 genes) (File S3). When compared with the entire list of Vglut1-RiboTag profiling in our control and DLK(cKO), we found CA1 genes tended to be expressed more in DLK(cKO) mice, compared to control (Fig.S6K), while CA3 genes showed a slight enrichment in control though the trend was less significant, and were less clustered towards one genotype (Fig.S6L). Moreover, many CA1 genes related to cell-type specification, such as *FoxP1*, *Satb2*, *Wfs1*, *Gpr161*, *Adcy8*, *Ndst3*, *Chrna5*, *Ldb2*, *Ptpru*, and *Ntm,* did not show significant downregulation when DLK was overexpressed. These observations imply that DLK likely specifically down-regulates CA1 genes both under normal conditions and when overexpressed, with a stronger effect on CA1 genes, compared to CA3 genes. Overall, the informatic analysis suggests that decreased expression of CA1 enriched genes may contribute to CA1 neuron vulnerability to elevated DLK, although it is also possible that the observed down-regulation of these genes is a secondary effect associated with CA1 neuron degeneration.

### DLK regulates translation of JUN and STMN4

The transcription factor c-Jun is a key downstream factor in DLK and JNK signaling (Hirai et al., 2006; Itoh et al., 2009; Welsbie et al., 2017). Our RiboTag analysis suggests that expression and translation of *Jun* mRNA to be significantly dependent on DLK expression levels (Fig.3A,B). To further test this observation, we performed immunostaining of hippocampal tissues using an antibody recognizing total c-Jun. In control mice, glutamatergic neurons in CA1 had low but detectable c-Jun immunostaining at P10 and P15, but reduced intensity at P60; those in CA3 showed an overall low level of c-Jun immunostaining at P10, P15 and P60; and those in DG showed a low level of c-Jun immunostaining at P10 and P15, and an increased intensity at P60 (Fig.S7A,C,E). In *Vglut1 ^Cre/+^;H11-DLK^iOE/+^* mice at P10 when no discernable neuron degeneration was seen in any regions of hippocampus, only CA3 neurons showed a significant increase of immunostaining intensity of c-Jun, compared to control (Fig.S7A). In P15 mice, we observed further increased immunostaining intensity of c-Jun in CA1, CA3, and DG, with the strongest increase (∼4-fold) in CA1, compared to age-matched control mice (Fig.S7C). The overall increased c-Jun staining is consistent with RiboTag analysis. We also analyzed *Vglut1^Cre/+^;DLK(cKO)^fl/fl^* mice at P60, and observed a trend for decreased c-Jun in CA3 (Fig.S7E); the modest effects of DLK(cKO) on c-Jun proteins could be due to detection limitations for low levels of c-Jun. As phosphorylation of c-Jun (p-c-Jun) is known to reflect activation of DLK and JNK signaling (Hirai et al., 2006), we further investigated p-c-Jun levels in these mice. In control mice at P10, P15, only a few neuronal nuclei showed strong staining with p-c-Jun in CA1, CA3 and DG. In *Vglut1 ^Cre/+^;H11-DLK^iOE/+^*mice, we observed increased p-c-Jun positive nuclei in CA1 at P10, and strong increase in CA1 (∼10-fold), CA3 (∼6-fold), and DG (∼8-fold) at P15 (Fig.S7B,D). The levels of p-c-Jun remained elevated in the surviving neurons in all three regions of *Vglut1 ^Cre/+^;H11-DLK^iOE/+^* mice at P60 (Fig.S4C). In *Vglut1^Cre/+^;DLK(cKO)^fl/fl^*mice, p-c-Jun levels in CA3 showed a significant reduction, with the trend of reduced levels in CA1 and DG, compared to control mice (Fig.S7F, P60). These results are consistent with a conclusion that translation of *Jun* mRNAs and phosphorylation of c-Jun show dependency on levels of DLK, with CA1 neurons showing higher dependence upon DLK overexpression.

The Stathmin family of proteins is thought to regulate microtubules through sequestering tubulin dimers (Charbaut et al., 2001; Chauvin and Sobel, 2015). This family of proteins includes four genes, all of which were identified in our hippocampal glutamatergic neuron translatome (Fig.S8A), and only *Stmn4* showed significant up-regulation in DLK(iOE) and down-regulation in DLK(cKO), respectively (Fig.3A,B,). We verified STMN4 protein expression by western blot analysis of hippocampal protein extracts. In control mice, we detected the levels of STMN4 to peak around P8 (Fig.S8B,C,F,G). The abundance of STMN4 in DLK(cKO) and DLK(iOE) was subtly altered, which could be due to broad expression of STMN4 in hippocampus masking specific changes in glutamatergic neurons. We thus examined *Stmn4* mRNAs in hippocampus by RNAscope. *Stmn4* mRNAs were present in glutamatergic neurons across all regions of the hippocampus, with strongest expression in CA1 pyramidal neurons. While *Stmn4* mRNA puncta number in these neurons was comparable between *Vglut1^Cre/+^;DLK(cKO)^fl/fl^*mice and control, in *Vglut1 ^Cre/+^;H11-DLK^iOE/+^*mice, glutamatergic neurons in CA1, CA3, and DG all showed upregulation of *Stmn4,* compared to control (Fig.4A-C, S9A,C). These data support a role of DLK in modulating expression and translation of *Stmn4*.

**Figure 4.**
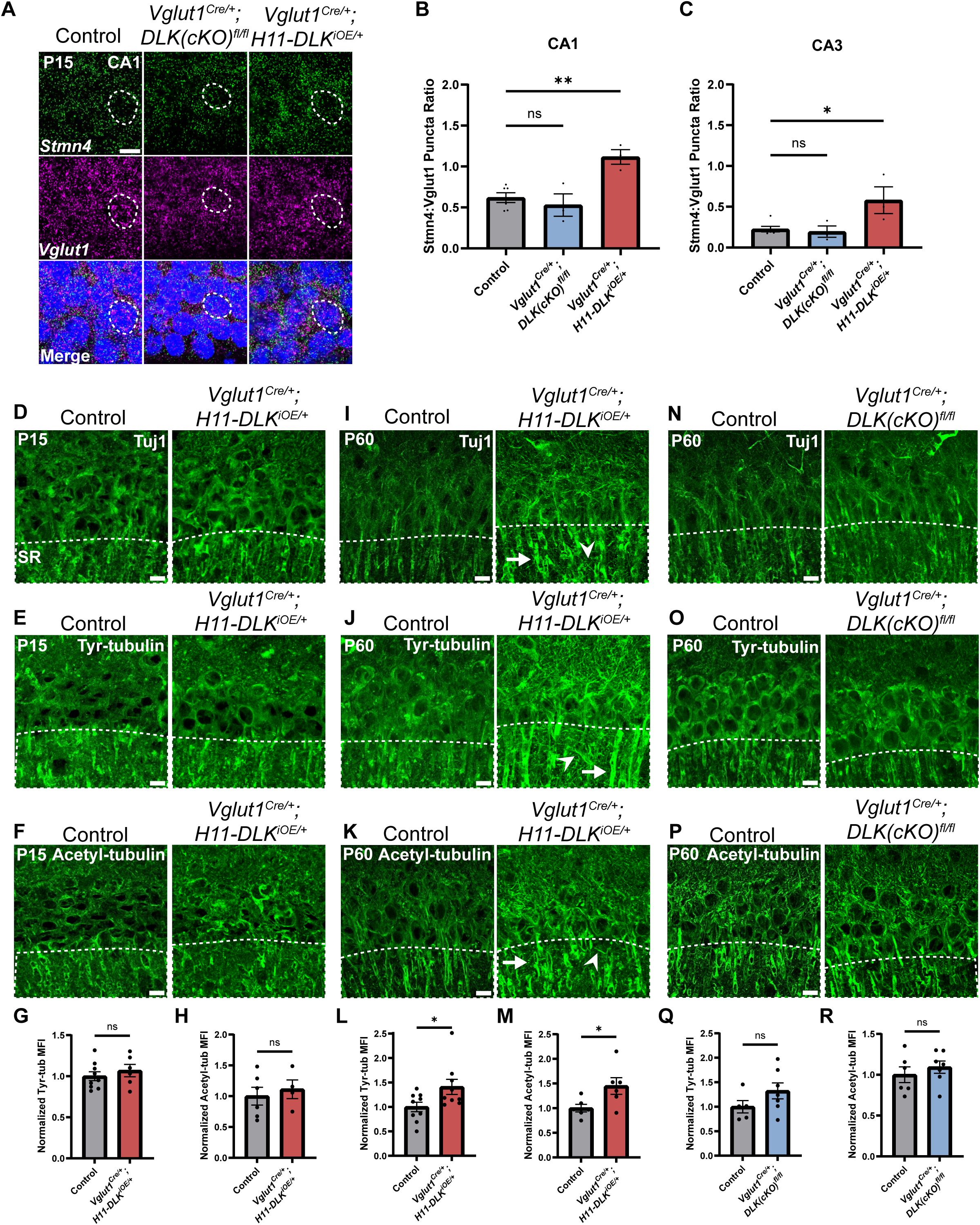
*Stmn4* and microtubule homeostasis show dependency on the expression levels of DLK. (A) Confocal single-slice image of RNAscope analysis of *Stmn4* and *Vglut1* mRNAs in hippocampal neurons. Dashed circle outlines single nuclei. Scale bar, 10μm. (B,C) Quantification of the ratio of *Stmn4* to *Vglut1* RNAscope puncta in same nuclei of CA1 and CA3 neurons, respectively. N= 6,3,3 mice of respective genotypes, quantified from >50 cells per genotype from 4 sections per mouse. Statistics: One way ANOVA with Dunnett’s multiple comparison test, ns, not significant; ** P<0.01; **** P<0.0001. (D-F) Confocal z-stack (max projection) images of CA1 immunostained for Tuj1, tyrosinated tubulin, and acetylated tubulin, respectively, in control and *Vglut1^Cre/+;^H11-DLK^iOE/+^* mice of P15. SR: stratum radiatum. (G-H) Normalized mean fluorescence intensity (MFI) of tyrosinated and acetylated tubulin, respectively, after thresholding signals in SR in CA1 (dashed outlines on images in E-F). N=9, 6 mice, 3 sections averaged per mouse in (G); N=6, 4 mice, 3 sections averaged per mouse in (H). (I-K) Confocal z-stack (max projection) images of immunostained CA1 sections for Tuj1, tyrosinated tubulin, and acetylated tubulin, respectively, in control and *Vglut1^Cre/+;^H11-DLK^iOE/+^* mice of P60. (L-M) Normalized MFI of tyrosinated and acetylated tubulin, respectively, after thresholding signal in SR in CA1 (dashed outlines on images in J-K). N=9, 9 mice, 3 sections averaged per mouse in L; N=6, 6 mice, 3 sections averaged per mouse in M. (N-P) Confocal z-stack (max projection) images of immunostained CA1 sections for Tuj1, tyrosinated tubulin, and acetylated tubulin, respectively, in control and *Vglut1^Cre/+^;DLK(cKO)^fl/fl^*mice of P60. (Q-R) Normalized MFI for tyrosinated and acetylated tubulin, respectively, after thresholding signal in SR in CA1 (dashed outlines on images in O-P). N=5, 7 mice, 3 sections averaged per mouse in Q; N=6, 7 mice, 3 sections averaged per mouse in R. All tubulin images shown as maximum projection of z-stack. Scale bar, 10μm. In I-J, arrows point to apical dendrites with elevated immunostaining signal; arrowheads point to thin neurites with elevated signal. Statistics in (G,H,L,M,Q,R): Unpaired t-test. ns, not significant; * P<0.05.

### Elevated DLK signaling may disrupt microtubule homeostasis in hippocampal CA1 neurons

Substantial studies from other types of neurons in mice and invertebrate animals have linked DLK signaling with the regulation of microtubule cytoskeleton (Asghari Adib et al., 2018; Jin & Zheng, 2019; Tedeschi & Bradke, 2013). To assess whether DLK affects microtubules in hippocampal glutamatergic neurons, we performed Tuj1 immunostaining. We did not detect obvious changes in *Vglut1^Cre/+^;DLK(cKO)^fl/fl^* when compared to controls at P60 (Fig.1E). In *Vglut1 ^Cre/+^;H11-DLK^iOE/+^* mice at P15 expression levels and patterns of neuronal microtubules in each region of hippocampus appeared similar to control (Fig.4D, S9E), although we found the overall Tuj1 staining pattern at P15 to be less defined and consistent. By P60, many CA1 neurons died and the hippocampus exhibited thinning of all strata within CA1, and the Tuj1 staining pattern became less organized in parallel dendrites in the stratum radiatum (SR) region of CA1 (Fig.4I). Increased Tuj1 staining in thin branches extended in varied directions, with bright staining seen in the apical dendrite near the pyramidal neuron cell body.

Several post-translational modifications of microtubules are thought to correlate with stable or dynamic state of microtubules. To explore whether DLK expression levels affected microtubule post-translational modifications, we performed immunostaining for acetylated tubulin, a modification generally associated with stable, longer-lived microtubules, and tyrosinated tubulin, a terminal amino acid that can be removed and is typically found on dynamic microtubules (Janke and Magiera, 2020). We detected no significant difference in the staining pattern and intensity of either tyrosinated tubulin or acetylated tubulin in *Vglut1^Cre/+^;DLK(cKO)^fl/fl^* mice, compared with age-matched control mice (Fig.4N-R). In *Vglut1 ^Cre/+^;H11-DLK^iOE/+^* mice at P15, both tubulin modifications showed no significant differences in pattern or intensity in CA1 SR, compared to age-matched control mice (Fig.4E-H), despite neuron death beginning in CA1. By P60, we observed increased staining intensity of acetylated tubulin and tyrosinated tubulin in the apical dendrites of surviving neurons in *Vglut1 ^Cre/+^;H11-DLK^iOE/+^*mice, particularly with tyrosinated tubulin staining revealing bright signals on small, thin branches (Fig.4J-M). To discern whether such microtubule modification changes were from neurons, we immunostained tissue sections with antibodies for MAP2, a neuron specific microtubule associated protein. We observed bright MAP2 signal in thin branches extending in varied directions in *Vglut1 ^Cre/+^;H11-DLK^iOE/+^* mice, compared to age-matched control mice (Fig.S9F,G). Together this analysis suggests increased DLK expression may likely alter neuronal microtubule homeostasis and/or integrity.

### Increasing DLK expression alters synapses in dorsal CA1

A theme revealed in our hippocampal glutamatergic neuron RiboTag profiling suggests that translation of synaptic proteins may depend on the expression levels of DLK. To evaluate this observation, we examined synapses in the hippocampus by immunostaining for Bassoon, a core protein in the presynaptic active zone, Vesicular Glutamate Transporter 1 (VGLUT1) for synaptic vesicles, and Homer1, a post-synaptic scaffolding protein. In control mice and *Vglut1^Cre/+^;DLK(cKO)^fl/fl^*at P60, Bassoon staining in stratum radiatum (SR) of dorsal CA1, where CA3 neurons synapse onto CA1 dendrites, showed discrete puncta that were mostly apposed to the postsynaptic marker Homer1, representing properly formed synapses (Fig.5A). We measured size and density by counting Bassoon and Homer1 puncta and the sites where Bassoon and Homer1 overlap, a proxy for synapses. We detected no significant difference in *Vglut1^Cre/+^;DLK(cKO)^fl/fl^*, compared to control (Fig.5A-F). To assess effects of DLK overexpression on synapses, we immunostained hippocampal sections from both P10 and P15, with age-matched littermate controls. Quantification of Bassoon and Homer1 immunostaining revealed no significant differences in CA1 SR and CA3 SR and SL in P10 mice of *Vglut1 ^Cre/+^;H11-DLK^iOE/+^* and control (Fig.S11A-F, S12A-J). In P15, Bassoon density and size in CA1 SR were comparable in both mice (Fig 5G, H, K), while Homer1 density and size were reduced in DLK(iOE) (Fig.5G,I, L). Overall synapse number in CA1 SR was similar in DLK(iOE) and control mice (Fig.5J). Similar analysis on CA3 SR and SL detected no significant difference from control (Fig.S12M-V). Staining of VGLUT1 protein showed less discrete puncta than those of Bassoon or Homer1, with small puncta and larger clusters of puncta close together (Fig.S10A,D). In DLK(cKO) we observed a trend towards an increased number of VGLUT1 puncta (p=0.0653) with no change to puncta size (Fig.S10A-C). In DLK(iOE) we observed fewer VGLUT1 puncta in SR, consistent with the analysis on Homer1 at P15, with no significant change to puncta size (Fig.S10D-F). These data reveal that while conditional knockout of DLK may not have a strong effect on glutamatergic synapses, increased expression of DLK leads to mild alteration in the CA1 region at P15, correlating with the onset of CA1 neurodegeneration.

**Figure 5.**
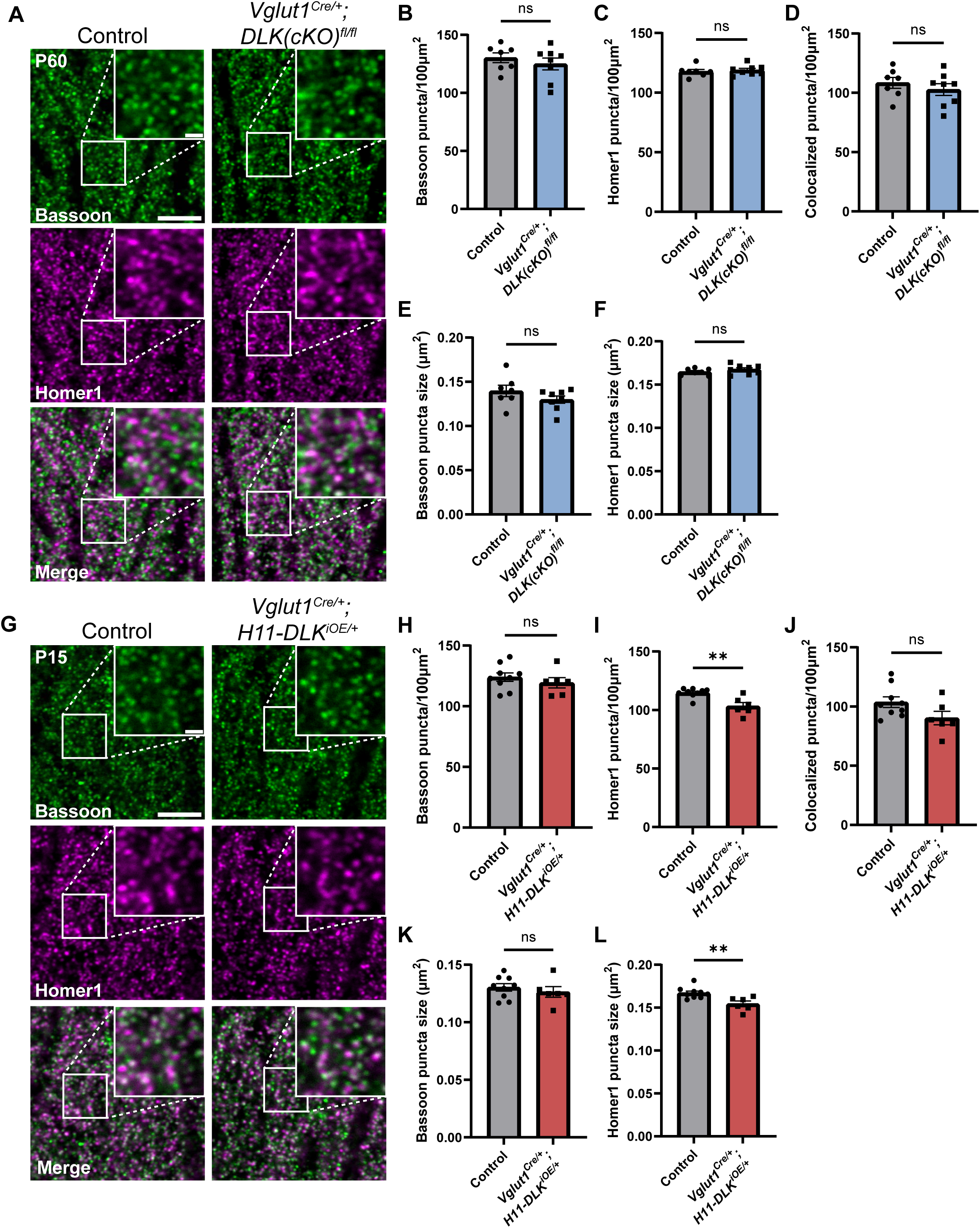
Hippocampal dorsal CA1 glutamatergic neurons show altered synapses following increased DLK expression. (A) Confocal single-slice images of Bassoon and Homer1 immunostaining in CA1 stratum radiatum (SR) of control and *Vglut1^Cre/+^;DLK(cKO)^fl/fl^*mice of P60. (B-C) Quantification of Bassoon and Homer1 puncta density, respectively. (D) Quantification of co-localization of Bassoon and Homer1. (E-F) Quantification of Bassoon and Homer1 puncta size. Data points represent average values per mouse from 3 sections. N=7 control, and 8 *Vglut1^Cre/+^;DLK(cKO)^fl/fl^*mice. Statistics: unpaired t-test or Mann-Whitney U test if not passing normality. ns, not significant. (G) Confocal single-slice images of Bassoon and Homer1 immunostaining in CA1 SR of control and *Vglut1^Cre/+^;H11-DLK^iOE/+^* mice of P15. (H-I) Quantification of Bassoon and Homer1 puncta density, respectively. (J) Quantification of co-localization of Bassoon and Homer1. (K-L) Quantification of Bassoon and Homer1 puncta size. Data points represent average values per mouse from 3 sections, N=9 control, and 6 *Vglut1^Cre/+^;H11-DLK^iOE/+^* mice. Statistics: unpaired t-test or Mann-Whitney U test if not passing normality. ns, not significant; ** P<0.01. Scale bars, 5μm in panel images, and 1μm in enlarged images.

### High levels of DLK cause short neurite formation in primary hippocampal neurons

To gain better resolution on how DLK expression levels affect glutamatergic neuron morphology and synapses, we next turned to primary hippocampal cultures. To enable visualization of Vglut1 positive neurons, we introduced a floxed Rosa26-tdTomato reporter (Madisen et al., 2010) into *Vglut1^Cre/+^, Vglut1^Cre/+^;DLK(cKO)^fl/fl^,* and *Vglut1 ^Cre/+^;H11-DLK^iOE/+^* mice. We prepared primary hippocampal neurons from P1 pups of respective crosses (Methods), so around 1⁄4 of glutamatergic (VGlut1-Cre) neurons in the cultures had both tdTomato and the genotype of interest (DLK(cKO/cKO), WT, or DLK(iOE/+)). We did not notice an obvious effect of DLK(iOE) or DLK(cKO) on neuron density in cultures at DIV2. To assess neuronal type distribution in our cultures, we immunostained DIV14 neurons with antibodies for Satb2, as a CA1 marker (Nielsen et al., 2010), and Prox1, as a marker of DG neurons (Iwano et al., 2012). We did not observe significant differences in the proportion of cells labeled with each marker in DLK(cKO) or DLK(iOE) cultures (Fig.S13E). These data are consistent with the idea that DLK signaling does not have a strong role in neuron-type specification both *in vivo* and *in vitro*.

We verified DLK protein pattern and levels by immunostaining with DLK antibodies (Fig.6A). In DIV2 neurons from control mice, DLK was present in cell soma, likely reflecting Golgi apparatus localization as reported (Hirai et al., 2002), and showed a punctate pattern in neurites, particularly the axon growth cone regions. We also immunostained for STMN4 and observed a similar punctate localization in the cell soma, neurites, and growth cones (Fig.6A), in line with published data (Chauvin et al., 2008; Gavet et al., 2002). STMN4 puncta appeared to be non-overlapping with DLK (Fig.S13B). In DIV2 neurons from DLK(cKO), STMN4 exhibited a similar punctate pattern, with intensity comparable to that in control neurons. In neurons from DLK(iOE), DLK levels were increased, and STMN4 levels were also increased (Spearman correlation r=0.7454) (Fig.6A,B), supporting our RiboTag analysis. Expression of another member of Stathmin, STMN2, is associated with DLK-dependent responses in DRG neurons (Summers et al., 2020; Thornburg-Suresh et al., 2023). Although our RiboTag analysis did not identify significant changes of *Stmn2* (Fig.S8A), we tested whether STMN2 protein levels could be altered in our cultured hippocampal neurons. In DIV2 control neurons STMN2 staining showed punctate localization in the perinuclear region (Gavet et al., 2002; Lutjens et al., 2000), along with punctate signals in neurites and growth cones, similar to STMN4. By co-immunostaining analysis of DLK and STMN2, we detected a positive correlation between DLK and STMN2 (Fig.S13C,D, Spearman correlation r=0.4693), albeit to a moderate level in comparison to that of DLK and STMN4. These data suggest that in hippocampal glutamatergic neurons DLK has a stronger effect on STMN4 levels but may also regulate protein levels of STMN2.

**Figure 6.**
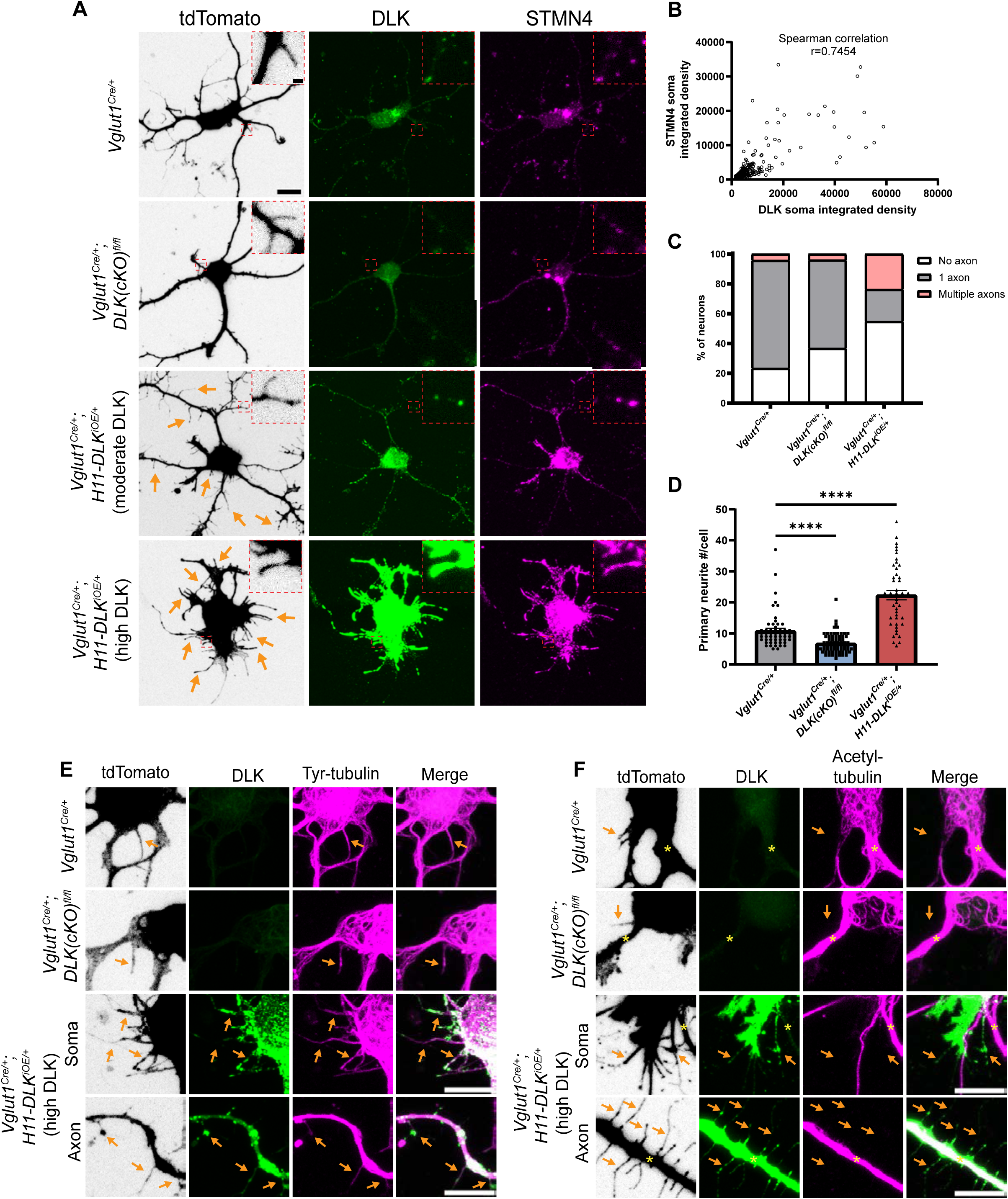
DLK promotes short neurite formation in primary cultured hippocampal neurons. (A) Confocal images of DIV2 primary hippocampal glutamatergic neurons immunostained with DLK and STMN4. Neurons with indicated genotypes are labeled by tdTomato from Cre-dependent Rosa26-tdTomato, generated from hippocampi in P1 pups from the following crosses: for control: *Vglut1^Cre/+^* X *Rosa26^tdT/+^*; for DLKcKO: *Vglut1^Cre/+^;DLK(cKO)^fl/fl^* X *Vglut1^Cre/+^;DLK(cKO)^fl/fl^;Rosa26^tdT/+^;* for DLKiOE: *H11-DLK^iOE/iOE^* X *Vglut1 ^Cre/+^;Rosa26^tdT/+^.* Orange arrows point to some of the thin neurites from neurons overexpressing DLK. Red dashes outline enlarged view of neurites. Scale bar, 10μm neuron, 1μm enlarged view. (B) Graph shows positive correlation between STMN4 immunostaining, measured as integrated density (Area X MFI) in neuronal soma, to integrated density of DLK immunostaining. N≥3 cultures/genotype, ≥60 cells/genotype. Spearman correlation r=0.7454. (C) Quantification of percentage of neurons with no, one, or more than one axon (defined by neurites longer than 90μm) in each genotype. Number of neurons: 47 from 3 Vglut1^Cre^ (control) cultures, 49 from 3 DLK(cKO) cultures, 42 from 4 DLK(iOE) cultures. Statistics: Fisher’s exact test shows significance (P<0.0001) between the three genotypes and number of axons. Axon formation in control vs DLK(cKO): P=0.1857. Formation of multiple axons in control vs DLK(cKO): P>0.9999. Axon formation in control vs DLK(iOE): P=0.0042. Formation of multiple axons in control vs DLK(iOE): P=0.0001. (D) Quantification of number of primary neurites, which include both branches and filopodia, per neuron. Number of neurons: 55 from 4 Vglut1^Cre^ (control) cultures, 70 from 4 DLK(cKO) cultures, 45 from 5 DLK(iOE) cultures. Statistics, Kruskal-Wallis test with Dunn’s multiple comparison test. **** P<0.0001. (E) Confocal z-stack images of tyrosinated tubulin immunostaining from DIV2 cultures of genotypes indicated, showing that filopodia structures (arrows) around the soma and axons of neurons with high expression of DLK have tyrosinated tubulin. (F) Confocal z-stack images of acetylated tubulin immunostaining from DIV2 cultures of genotypes indicated, showing that filopodia structures (arrows) around the soma and axons of neurons with high expression of DLK do not have acetylated tubulin. Asterisks indicate stable branches containing acetylated tubulin. Scale bar in E-F,10μm. Tyrosinated tubulin and acetylated tubulin staining shows saturated appearance to visualize staining in thin neurites.

In hippocampal cultures at DIV2 neurites are actively growing and establish thicker branches, which form dendrites and axons (Dotti et al., 1988). In our control cultures at DIV2, the majority of tdTomato labeled neurons developed multiple neurites from the cell soma, with one neurite developing into an axon, defined here as a neurite longer than 90 µm (Fig.S13A). Additionally, thin, often short, neurites were observed branching off from the cell soma, axons, dendrites, and growth cones. In DLK(cKO) cultures at DIV2, we observed a trend of more neurons without an axon (Fig.6C), though the differentiated axons appeared morphologically indistinguishable from control. The total number of neurites around the cell soma in DLK(cKO) neurons was significantly reduced, compared to control (Fig.6A,D). In DLK(iOE) cultures at DIV2, we observed a significant increase in the percentage of neurons without an axon and also neurons with multiple axons, compared to control cultures (Fig.6C, S13A). Moreover, neurons expressing high levels of DLK protein displayed an increased number of neurites either around the cell soma as primary neurites or as secondary neurites, compared to control (Fig.6A,D). Such neurites were typically thin, and frequently appeared short, like filopodia. These thin neurites sometimes developed a rounded tip and showed beading appearance, resembling degeneration (Fig.6A,E). These data suggest a role for DLK in neurite formation and axon specification in cultured hippocampal glutamatergic neurons.

We further analyzed microtubules in individual neurites of the DIV2 neurons. Control neurons exhibited staining for tyrosinated tubulin in differentiated axons and dendrites as well as filopodia and towards peripheral regions of growth cones (Fig.6E). Acetylated tubulin was present in differentiated axons and dendrites and in the central region of growth cones where stable microtubules were present (Fig.6F), and was absent from filopodia and microtubules in the peripheral regions of growth cones. The thin neurites in neurons expressing very high levels of DLK appeared to have thin bundles of microtubules, and these neurites generally were not associated with a growth cone or microtubules splaying apart at the end as was common in WT growth cones (Fig.6E). Neurites from neurons with high DLK expression also had tyrosinated tubulin, while acetylated tubulin was frequently absent (Fig.6E,F). Additionally, STMN4 was present in the thin neurites, especially in those with high levels of DLK (Fig.6A), suggesting the thin neurites may likely be dynamic in nature. These results suggest that in cultured hippocampal neurons high levels of DLK promotes formation of short, thin, dynamic branches.

### Increased DLK expression alters synapses in primary hippocampal neurons

Our RiboTag data showed enrichment of synaptic genes in both DLK(cKO) and DLK(iOE) (Fig. 3F, S6J, File S1 WT vs DLK(cKO) DEGs, File S2 WT vs DLK(iOE) DEGs). Some of these genes function in cell adhesion, calcium signaling, and AMPA receptor expression, which may affect dendritic spine morphology and synaptic connections. Increased DLK levels led to reduced Homer1 density in hippocampal tissue (Fig.5J). To further investigate the effects of DLK on synapses, we immunostained the cultured hippocampal neurons at DIV14 with Bassoon. The control neurons showed discrete Bassoon puncta in axons (Fig.7A-C). DLK(cKO) neurons showed no significant change in Bassoon puncta size or density. In contrast, DLK(iOE) neurons showed abnormal Bassoon staining that was larger and irregular in shape (Fig.7A-C), suggesting that high levels of DLK disrupted presynaptic active zones.

**Figure 7.**
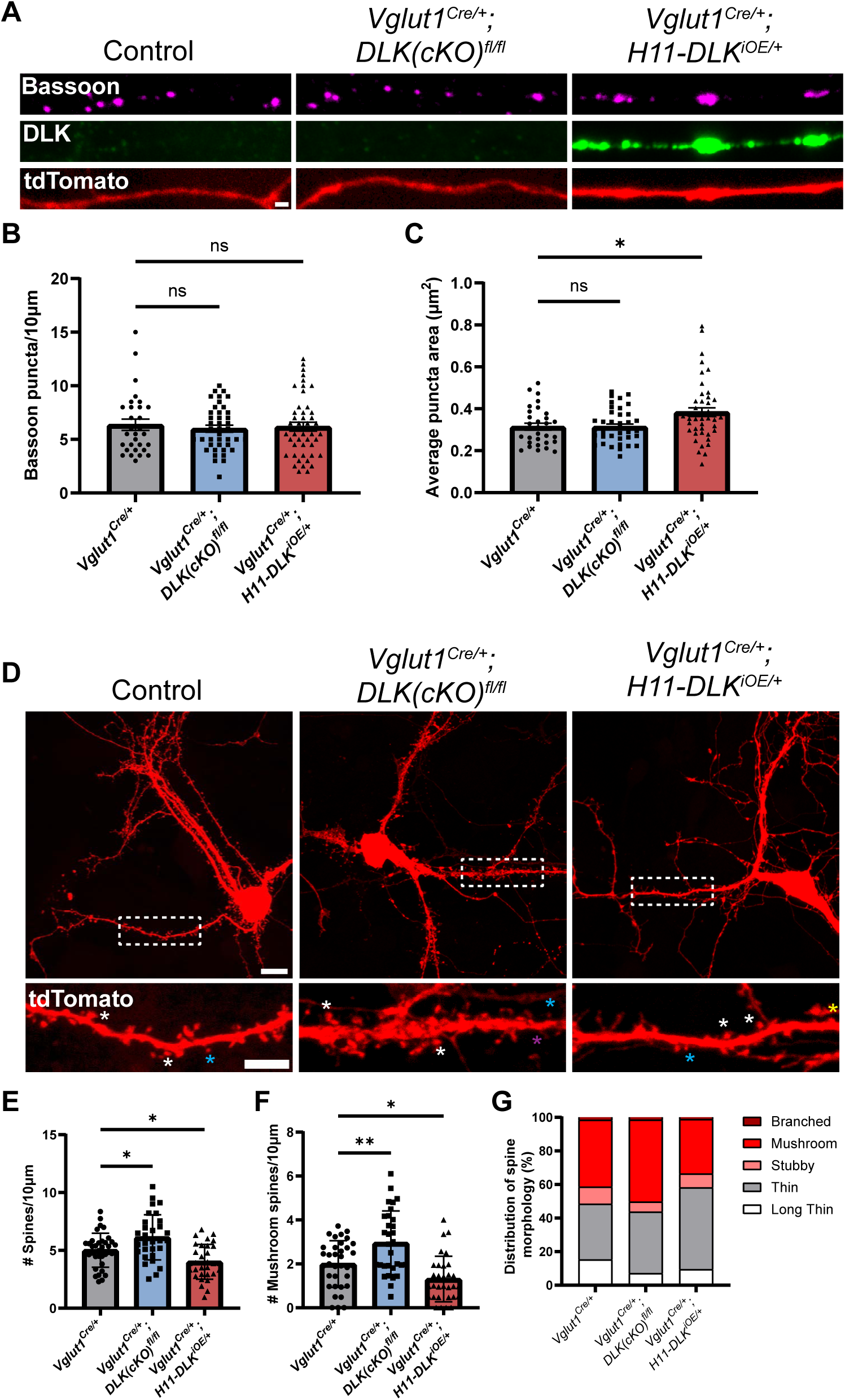
Increasing DLK expression alters synapse formation in primary cultured hippocampal neurons. (A) Confocal images of axons of DIV14 neurons of indicated genotype, co-stained with Bassoon and DLK. Neurons with indicated genotypes are labeled by tdTomato from Cre-dependent Rosa26-tdTomato generated from the following crosses: for control: *Vglut1^Cre/+^* X *Rosa26^tdT/+^*; for DLKcKO: *Vglut1^Cre/+^;DLK(cKO)^fl/fl^* X *Vglut1^Cre/+^;DLK(cKO)^fl/fl^;Rosa26^tdT/+^;* for DLKiOE: *H11-DLK^iOE/iOE^* X *Vglut1 ^Cre/+^;Rosa26^tdT/+^.* (B) Quantification of bassoon puncta density. (C) Quantification of average bassoon puncta size from individual neurons. Number of neurons: 30 from 3 Vglut1-cre (control) cultures, 41 from 3 DLK(cKO) cultures, 46 from 4 DLK(iOE) cultures. Statistics: One way ANOVA with Dunnett’s multiple comparison test. ns, not significant; * P<0.05. (D) Confocal z-stack images of DIV14 neurons of indicated genotype, labeled by Rosa26-tdTomato. Dashed boxes outline dendrites enlarged below for dendritic spines. Asterisks provide some examples of spine types; long thin with purple; thin with blue; mushroom with white; stubby with yellow. Scale bar, 10μm top, 5μm bottom. (E) Quantification of dendritic spine density. (F) Quantification of mushroom spine density. (G) Distribution of spine types. (E-G) Number of neurons: 35 from 3 Vglut1-cre (control) cultures, 31 from 3 DLK(cKO) cultures, 31 from 3 DLK(iOE) cultures. Statistics: One way ANOVA with Dunnett’s multiple comparison test. * P<0.05; ** P<0.01.

Morphology of dendritic spines is associated with differences in maturity, with mushroom spines representing a more mature morphology than thin spines (Yoshihara et al., 2009). To assess how DLK expression levels affect dendritic spine morphology and frequency in cultured neurons, we evaluated both the density and type of dendritic spines formed at DIV14 on neurons with spines, visualized by tdTomato. We categorized spine morphology in different types, following previous studies (Risher et al., 2014). Neurons from DLK(cKO) showed spines at a higher density than control neurons, with significantly more mushroom spines (Fig.7D-G). In contrast, neurons from DLK(iOE) cultures had a reduced density of dendritic spines compared to control neurons (Fig.7D, F). DLK(iOE) neurons also formed spines that tended to be more immature, with significantly fewer mushroom spines and a higher percentage of thin spines (Fig.7D, G). These results reveal that expression levels of DLK appear to be inversely correlated with spine density and maturity.

## Discussion

### Selective vulnerability of hippocampal glutamatergic neurons to increased DLK expression

Under normal conditions, the abundance of endogenous DLK in many parts of the brain is generally kept at low level. Elevated DLK signaling has been associated with traumatic injury and implicated in Alzheimer’s disease and other neurodegenerative conditions (Asghari Adib et al., 2018; Huang et al., 2017; Jin & Zheng, 2019; Le Pichon et al., 2017; Tedeschi & Bradke, 2013). Despite its broad expression, we know little about DLK’s role in the central nervous system. In this study, we combined conditional knockout and overexpression of DLK to uncover its roles in the hippocampal glutamatergic neurons. Our finding that conditional deletion of DLK in the glutamatergic neurons using *Vglut1^Cre^* in late embryonic development does not cause discernable morphological defects is consistent with the previous reports that hippocampal neurons are largely normal in constitutive knockout of DLK (Hirai et al., 2006; Hirai et al., 2011). In contrast, induced overexpression of DLK, which leads to activation of JNK signaling evidenced by increased p-c-Jun, causes the glutamatergic neurons in dorsal CA1 and dentate gyrus to undergo pronounced death, while CA3 neurons appear less vulnerable even under chronic elevated DLK expression. The levels of DLK in our DLK(iOE) mice model appear comparable to those reported under traumatic injury and chronic stress. The pattern of DLK-induced neuronal death shares similarity to the differential vulnerability of CA1 and CA3 neurons reported in patients with Alzheimer’s disease (West et al., 1994), and animal models of oxidative stress (Wilde et al., 1997), ischemia (Smith et al., 1984), and glutamate excitotoxicity from NMDA (Vornov et al., 1991). The dorsal-ventral hippocampal neuron death pattern associated with increased expression of DLK is also similar to that observed in animal models of ischemia (Smith et al., 1984). Such regional differences of hippocampal neurons in response to insults or genetic manipulation may be attributed to multiple factors, such as the nature of the neural network (Viana da Silva et al., 2024), intrinsic differences between CA1 and CA3 neurons in their abilities to buffer calcium changes, mitochondrial stress, protein homeostasis, glutamate receptor distribution (Schmidt-Kastner, 2015), and as discussed further, the degree to which transcription factors, such as p-c-Jun or other AP1 factors, are activated under different conditions.

### DLK-dependent cellular network exhibits commonality and cell-type specificity

DLK to JNK signaling is known to lead to transcriptional regulation. Several studies have used transcriptomic profiling to reveal DLK-dependent gene expression in different regions of the brain, such as cerebellum and forebrain, and in specific neuron types, such as DRG neurons and RGC neurons following axon injury or nerve growth factor withdrawal (Goodwani et al., 2020; Hu et al., 2019; Larhammar et al., 2017; Le Pichon et al., 2017; Shin et al., 2019; Watkins et al., 2013). One recent study reported RiboTag profiling of the DLK-dependent gene network in axotomized spinal cord motor neurons (Asghari Adib et al., 2024). In agreement with the overall findings from these studies, we find that loss of DLK in hippocampal glutamatergic neurons results in modest expression changes in a small number of genes, while overexpression of DLK leads to expression changes in a larger set of genes. Gene ontology analysis of our hippocampal glutamatergic neuron translatome reveals a similar set of terms as found in the other expression studies, including neuron differentiation, apoptosis, ion transport, and synaptic regulation.

Comparison of the translational targets of DLK in our study with these prior analyses also shows notable differences that are likely specific to neuron-type and contexts of experimental manipulations. For example, we find a strong induction of *Jun* translation associated with increased expression of DLK, but no significant changes in *Atf3* or *Atf4* translation, which were reported to show DLK-dependent increases in axotomized spinal cord motoneurons, and injured RGCs and DRGs (Asghari Adib et al., 2024; Larhammar et al., 2017; Shin et al., 2019; Watkins et al., 2013). Most ATF4 target genes (Somasundaram et al., 2023) also show no significant changes in our hippocampal glutamatergic neuron translatome.

Moreover, we find a cohort of synaptic genes showing expression dependency on DLK (such as *Tenm3, Nptx1*, and *Nptxr),* but not any of the complement genes (*C1qa, C1qb, C1qc*), which are up-regulated in the regenerating spinal cord motor neurons where neuron-immune cell interaction has a critical role (Asghari Adib et al., 2024). Genomic structure and regulation intrinsic to the cell type may be a major factor underlying the gene expression differences in ours and other studies. Elevated DLK signaling in axotomized neurons may promote a strong regenerative response through activation of transcription factors, such as ATF3 and ATF4, whereas JUN and others that are actively expressed in hippocampal neurons may lead to a strong effect on refining synapses in response to DLK signaling. Overall, ours and the previous studies underscore the importance of systematic dissection of molecular pathways to understand neuron-type specific functionality to DLK signaling.

Our analysis of the spatial expression patterns of genes that showed association with DLK expression levels provides molecular insight to the differential vulnerability of hippocampal glutamatergic neurons under neurodegenerative conditions. We find that a select set of genes enriched in CA1 are up-regulated in DLK knockout and down-regulated upon DLK overexpression. The c-Jun transcription factor has a key role in hippocampal cell death responses as mutations preventing c-Jun phosphorylation led to decreased neuronal apoptosis in the hippocampus following treatment with kainic acid (Behrens et al., 1999). Basal levels of c-Jun and phosphorylated c-Jun in hippocampus are generally low (Goodwani et al., 2020; Pozniak et al., 2013). We find modest reductions in p-c-Jun in DLK(cKO) glutamatergic neurons, consistent with previous studies of the constitutive knockout of DLK (Hirai et al., 2006). In contrast, in DLK(iOE) neurons, translation of c-Jun and phosphorylation of c-Jun are increased, with CA1 neurons exhibiting higher increase than CA3 neurons. The c-Jun promoter has consensus AP1 sites, and c-Jun can regulate its own expression levels in cancer cell lines (Angel et al., 1988), NGF-deprived sympathetic neurons (Eilers et al., 1998) and kainic acid treated hippocampus (Mielke et al., 1999). While our data does not pinpoint the molecular changes explaining why CA3 would show less vulnerability to increased DLK, we may speculate that DLK(iOE) induced signal transduction amplification may differ in CA1 vs CA3. CA1 genes appear to be more strongly regulated than CA3 genes, consistent with our observation that increased c-Jun expression in CA1 is greater than that in CA3. Other parallel molecular factors may also contribute to resilience of CA3 neurons to DLK(iOE), such as HSP70 chaperones, different JNK isoforms, and phosphatases, some of which showed differential expression in our RiboTag analysis of DLK(iOE) vs WT (shown in File S2. WT vs DLK(iOE) DEGs). Together with other genes that show dependency on DLK, the DLK and Jun regulatory network contributes to the regional differences in hippocampal neuronal vulnerability under pathological conditions.

### Conserved functions of DLK in regulating Stathmins

Stathmins are tubulin binding proteins broadly expressed in many types of neurons. Several studies have reported that DLK can regulate the expression of different Stathmin isoforms in multiple neuron types under injury conditions (Asghari Adib et al., 2024; DeVault et al., 2024; Hu et al., 2019; Larhammar et al., 2017; Le Pichon et al., 2017; Shin et al., 2019). In the hippocampus *Stmn2* is expressed at a higher level than *Stmn4*, with the relative ratios of *Stmn4*:*Stmn2* in hippocampus much higher than in DRGs (Zeisel et al., 2018). We find that DLK can modulate expression and translation of *Stmn4* in hippocampal neurons. ChIP-seq data for Jun from ENCODE (ENCSR000ERO) suggest a possible binding of Jun in the promoter region of Stmn4 (ENCODE Project Consortium, 2012; Luo et al., 2020). STMN4 expression in hippocampus peaks around P8, correlating to neurite outgrowth and synapse formation and pruning (Paolicelli et al., 2011). At the level of hippocampal tissue, loss of DLK causes no detectable changes to microtubules, while increased levels of DLK appear to alter microtubule homeostasis in dendrites, with generally increased levels of both stable and dynamic microtubule markers. The CA1 neurons in DLK(iOE) also show fewer parallel microtubule arrays of apical dendrites, with short branches extending in varied directions. Our results from primary hippocampal neurons support roles for DLK in both short neurite and axon formation, similar to observations in cortical neurons, where DLK contributes to stage specific regulation of microtubules (Hirai et al., 2011). In primary cortical neurons overexpression of STMN4 can increase neurite length and branching when an epigenetic cofactor regulating MT dynamics was knocked down (Tapias et al., 2021). We speculate that DLK-dependent regulation of STMN4 and other STMNs may have a critical role in the long-term cytoskeletal rearrangements for neuronal morphology and synapse formation or stability. Nonetheless, as Stmns have considerable redundancy in expression and function, changes in STMN4 alone are unlikely to be a major factor for the observed hippocampal regional neuron death.

### Conserved roles of DLK in synapse formation and maintenance

The *in vivo* functions of the DLK family of proteins were first revealed in studies of synapse formation in *C. elegans* and Drosophila (Collins et al., 2006; Nakata et al., 2005). Our hippocampal glutamatergic neuron translatomic data extends this function by revealing a strong theme of DLK-dependent network in synapse organization, such as adhesion molecules and regulation of trans-synaptic signaling, especially related to AMPA receptor expression and calcium signaling, such as changes in Neuronal Pentraxin 1 and Neuronal Pentraxin Receptor *Nptx1* and *Nptxr* (Gómez de San José et al., 2022). From our synapse analysis in culture, we find increased DLK alters pattern of presynaptic protein Bassoon, consistent with the findings on *C. elegans* and Drosophila synapses (Nakata et al., 2005; Collins et al., 2006). We also find DLK regulates dendritic spine morphology, with loss of DLK associated with a greater number of spines with more mature spine morphology, while increased DLK was associated with fewer and less mature spines. These results are similar to that observed in layer 2/3 cortical neurons where loss of DLK is associated with larger dendritic spines in developing neurons and higher density of spines when exposed to Aβ plaques, which lead to loss of nearby spines (Le Pichon et al., 2017; Pozniak et al., 2013). In CA1 dendritic regions, DLK overexpression reduced Homer1 density, suggesting synaptic defects may correlate with the onset of degeneration. In axotomized spinal cord motor neurons DLK induces activation of complement, leading to microglial pruning of synapses in injured motoneurons (Asghari Adib et al., 2024). These data support a conserved role of DLK in synapse formation and maintenance, through regulating the translation of genes involved in neuron outgrowth, synaptic adhesion, and synapse activity.

### Limitation of our study

We have investigated roles of DLK in hippocampal glutamatergic neuron development, synapse regulation, and neuron death processes. We infer that DLK-dependent expression and translation of CA1 enriched genes may likely play roles in regional vulnerability to increased DLK signaling. However, our RiboTag profiling was performed with whole hippocampus at time when CA1 death was noticeable. Our analysis of spatial expression patterns of DLK-dependent genes relies on available data from adult animals, which may not reflect the patterns at P15, or in response to altered DLK. We cannot rule out that some of the decreased expression of CA1 enriched genes in DLK(iOE) could be secondary due to neuronal death that could result in fewer CA1 neurons present in our mRNA samples. Our analysis also does not directly address why CA3 neurons are less vulnerable to increased DLK expression. Future studies using cell-type specific RiboTag profiling and other methods at a refined time window will be required to address how DLK dependent signaling interacts with other networks underlying hippocampal regional neuron vulnerability to pathological insults. While we find evidence for apoptosis, other forms of cell death may also occur. Additional experiments will be needed to elucidate *in vivo* roles of STMN4 and its interaction with other STMNs. It is worth noting that a systematic analysis of gene networks in neuron types selectively vulnerable to Alzheimer’s disease has suggested processes related to axon plasticity and synaptic vesicle transmission, particularly with relation to microtubule dynamics, may be involved in the neuronal vulnerability (Roussarie et al., 2020). Combining gene profiling of specific cell types in hippocampus with advanced technology in function dissection will continue to provide clarification to roles of DLK in the central nervous system under normal and pathological conditions.

## Acknowledgements

We thank members of our labs for their support and valuable discussion throughout this work. We are grateful to Emily Griffin in Susan Ackerman’s lab and Caitlin Rodriguez in Aaron Gitler’s lab for advice on troubleshooting immunoprecipitation of ribosomes for RiboTag, to Brenda Bloodgood for advice with RNAscope experiments, Gentry Patrick, Lara Dozier, and Frank Bradke for their guidance in primary hippocampal cultures, Megan Williams and Stacey Glasgow for advice and CA1 and CA3 neuron antibodies, and Gareth Thomas for discussion and comments. This publication includes data generated at the UC San Diego IGM Genomics Center utilizing an Illumina NovaSeq 6000 that was purchased with funding from a National Institutes of Health SIG grant (#S10 OD026929). D.A was supported by the TÜBİTAK 2214-A International Research Fellowship Programme. E.M.R received an Innovative Research Grant from the Kavli Institute for Brain and Mind. This work was supported by a grant from NIH (NS R35 127314 to Y.J.).

## Author Contributions

Conceptualization, EMR and YJ; Methodology, EMR, YL, BZ, and YJ; Investigation, EMR, DA, SZ, and QP; Resources, BZ and YJ; Writing-Original Draft, EMR and YJ; Writing-Review and Editing, EMR, DA, SZ, YL, BZ, and YJ; Supervision, YJ; Funding Acquisition, EMR and YJ.

## Declaration of Interests

The authors declare no competing interests.

## Data and code availability

Sequencing datasets have been deposited in the Gene Expression Omnibus (Accession GSE266662).

## Supplemental Materials: 14 Supplemental Figures, 1 Supplemental Table, 3 Supplemental files

Document S1. Supplemental methods, Figures S1-S14 and Table S1

File S1. Excel file containing DLK(cKO) differentially expressed genes

File S2. Excel file containing DLK(iOE) differentially expressed genes

File S3. Excel file containing CamK2 and Grik4 RiboTag enriched genes

## Methods

### EXPERIMENTAL MICE

All animal protocols were approved by the Animal Care and Use Committee of the University of California San Diego. Map3k12fl (*DLK(cKO)^fl/fl^*) allele was made by Dr. Lawrence B. Holzman (Univ. Penn) and reported in (Chen et al., 2016; Li et al., 2021; Saikia et al., 2022). Map3k12 (*H11-DLK^iOE/+^*) transgene was described in Li et al., 2021. Vglut1^Cre^ allele (JAX stock #023527) was described in Harris et al., 2014. RiboTag allele (JAX stock #029977) was described in Sanz et al., 2009. ROSA26-loxP-STOP-loxP-tdTomato fl/fl reporter line (JAX stock #007914) was constructed in Madisen et al., 2010. Standard mating procedure was followed to generate *Vglut1^Cre/+^;DLK(cKO)^fl/fl^* and *Vglut1 ^Cre/+^;H11-DLK^iOE/+^* experimental mice. Genotyping primers are in Supplemental table 1. Sibling control mice had either *Vglut1^Cre/+^* or *DLK(cKO)^fl/fl^* or *H11-DLK^iOE/+^*allele alone. All experiments used both male and female mice. *Vglut1 ^Cre^* dependent tdTomato expression from *H11-DLK^iOE/+^* transgene was observed in most or all CA3, many CA1 neurons, with limited number of DG neurons at P15, similar to the described *Vglut1 ^Cre^*reporter line (Harris et al., 2014), and was throughout all regions by P60. *Vglut1 ^Cre/+^;H11-DLK^iOE/+^*mice around 4 months of age developed noticeable progressive motor deficits, which were likely unrelated to hippocampal glutamatergic neuron death, and were not studied further.

### WESTERN BLOTTING

Samples of brain tissue were lysed in ice-cold RIPA buffer (50mM Tris/HCl pH 7.4, 150mM NaCl, 0.5% DOC, 0.1% SDS, 1% NP-40 freshly supplemented with protease inhibitor cocktail and 1mM PMSF). Tissues were homogenized by Dounce homogenization using 30 passes pestle A and 30 passes pestle B. Samples were spun down at 13,000 x g for 10 min at 4C. Supernatants were collected, and protein concentration was determined using the BCA assay (Thermo Scientific, 23227). Equal concentration of proteins (∼10-20ng) were run on NuPAGE™ 4-12% Bis-Tris Gel, 1.0 mm (Invitrogen, NP0322BOX) with 20X NuPAGE™ MES SDS Running Buffer (Invitrogen, NP0002). Protein samples were transferred to a PVDF membrane (0.2 μm, Bio-RAD, 1620177) by Mini Trans-Blot Cell at 100 mA for 1 hour at 4°C. Membranes were blocked in 5% skim milk in TBST for 1 hour at room temperature, and then incubated with primary antibody in 3% BSA or 5% skim milk in TBST at 4°C overnight. Membranes were washed 3 x 10 min in TBST and incubated with 1:5000 of the appropriate HRP-conjugated secondary antibody in 3% BSA in TBST at room temperature for 1hr, then washed 3 x 10 min in TBST. Bands were detected using enhanced chemiluminescence (ECL) reagents (GE Healthcare, RPN2106) or Pico PLUS Chemiluminescent Substrate (Thermo Scientific, 34580) using a Licor Odyssey XF Imager. Molecular weight markers were PageRuler Plus Prestained Protein Ladder (Thermo Scientific, 26619) or Precision Plus Protein Ladder (Bio-Rad, 1610374).

Quantification of western blot images was performed by measuring identical size regions from each band, subtracting the background signal, and normalizing to internal actin controls for each sample. Time course analysis was further normalized to P1 WT protein levels. All images shown had N=3 biological replicates.

### IMMUNOFLUORESCENCE OF HIPPOCAMPAL TISSUES

Mice were transcardially perfused with saline solution followed by 4% PFA in PBS. Brains were dissected and post-fixed overnight in 4% PFA at 4°C, then washed with PBS and transferred to 30% sucrose in PBS for at least three days. Brains were mounted coronally for cryosectioning in OCT Compound (Fisher HealthCare, 4585) on dry ice. Sections were cut to 25µm thickness, divided evenly among six wells, and stored in PBS with 0.01% sodium azide at 4°C until staining. For immunostaining, free floating sections were washed 3 times in 0.2% Triton X-100 in PBS, blocked for 1 hour at room temperature in 5% donkey serum in 0.4% Triton X-100 in PBS, then incubated with primary antibodies in 2% donkey serum in 0.4% Triton X-100 in PBS overnight at 4°C rocking. Following three washes with 0.2% Triton in PBS, sections were incubated with secondary antibodies in 2% donkey serum in 0.4% Triton X-100 in PBS for 1 hour at room temperature. Sections were again washed three times with 0.2% Triton X-100 in PBS, stained with DAPI for 10 min (14.3mM in PBS) and washed three times in PBS before mounting on glass slides using Prolong Diamond Antifade Mountant. TUNEL staining was performed using the DeadEnd Fluorometric TUNEL System (Promega, G3250) with a modified protocol as described previously (Li et al., 2021).

### IMMUNOPRECIPITATION AND ISOLATION OF RIBOSOME ASSOCIATED mRNA

Immunoprecipitation of HA-tagged ribosomes was conducted following the protocol described in Sanz et al., 2019 (Sanz et al., 2019). Briefly, hippocampi from both hemispheres were dissected in ice cold PBS from mice of desired genotypes at postnatal day 15, and were stored at -80°C before further processing. Frozen tissues were homogenized by Dounce homogenization using 30 passes pestle A and 30 passes pestle B in 1.5mL homogenization buffer (50mM Tris, pH 7.5, 1% NP-40, 100mM KCl, 12mM MgCl_2_, 100ug/mL cycloheximide, cOmplete EDTA-free protease inhibitor cocktail (Roche), 1mg/mL heparin, 200U/mL RNasin, 1mM DTT). Following centrifugation at 10,000 x g for 10 minutes at 4°C, 5µg anti-HA high affinity (Roche) were added to the supernatant and incubated 4 hours rotating end-over-end at 4°C. The entire antibody-lysate solution was added to 400µl Protein G Dynabeads per sample overnight rotating end-over-end at 4°C. High salt buffer was prepared (50mM Tris, pH 7.5, 1% NP-40, 300mM KCl, 12mM MgCl_2_, 100ug/mL cycloheximide, 0.5mM DTT), and beads were washed 3 x 10 minutes using a magnetic tube rack. During the final wash, samples were transferred to a new tube, and beads were eluted in 350µl of RLT buffer (from the Qiagen RNAeasy Minikit) supplemented with 1% β-mercaptoethanol. RNA was extracted following manufacturer’s instructions in the RNAeasy Minikit (Qiagen). RNA integrity was measured using an Agilent TapeStation conducted at the IGM Genomics Center, University of California, San Diego, La Jolla, CA. All RNA for sequencing had RIN≥8.0, 28S/18S≥1.0.

To confirm immunoprecitipation in RiboTag IP samples, 10% of IP sample was isolated after final wash in high salt buffer. After removal of high salt buffer, protein was eluted in 2X RIPA buffer and 4x Laemmli Sample Buffer (Bio-Rad, 161–0747) by heating 10 min at 50°C. Beads were separated using a magnetic tube rack, the supernatant was isolated and beta-mercaptoethanol was added. Samples were boiled at 95°C for 10 min and centrifuged 5 min at 13,000 x g. Immunoprecipitated samples were separated by SDS-PAGE using Any kD Mini-PROTEAN TGX Precast Protein Gels (Bio-Rad, 4569034).

To ensure appropriate depletion of transcripts from non-Vglut1 expressing cells, we performed qRT-PCR analysis on representative marker genes for cell types in immunoprecipitated glutamatergic neuron RNA relative to whole hippocampal RNA. Briefly, RNAs isolated from whole hippocampi and immuoprecipitated from glutamatergic neurons were reverse transcribed to cDNA using Superscript III First Strand Synthesis System (Invitrogen, cat#18080051) following the manufacturer’s protocol. 100ng RNA/sample was reverse transcribed with random hexamers. iQ Sybr Green Supermix (Bio-Rad, #1708880) was used for qPCR, and mRNA levels of marker genes (*Vglut1* (glutamatergic neurons), *Wfs1* (CA1 neurons), *Gfap* (Astrocytes), and *Vgat* (inhibitory neurons)) were normalized to *Gapdh* expression. Expression levels of qRT-PCR samples was analyzed using the CFX Real-Time PCR Detection System and CFX Manager Software (Bio-Rad). Relative enrichment of marker genes was evaluated using the comparative *C_T_* method. All samples were run in triplicate. Primers for *Gapdh*, *Vglut1*, *Wfs1*, *Gfap*, and *Vgat* are from Furlanis et al., 2019 (see Supplemental table 1).

### SEQUENCING

Library preparation and sequencing for ribosome associated mRNAs were performed by the UCSD IGM Genomics Center using Illumina Stranded mRNA Prep. Sequencing was performed on NovaSeq S4 with PE100 reads.

### READ MAPPING

Following paired end RNA sequencing of isolated RNA, >24 million reads per sample were obtained (n=3DLK(iOE)/3WT, n=4 DLK(cKO)/4WT). The Galaxy platform was used for read mapping and differential expression analysis (Afgan et al., 2018). Read quality was checked using FastQC (version 0.11.8). Reads were mapped to the mouse reference genome (mm10) using STAR galaxy version 2.6.0b-1 with default settings (Dobin et al., 2013). Four DLK(cKO) and controls included 2 male and 2 female. For DLK(iOE), one female sample was removed from each genotype (control and DLK(iOE) due to read mapping variability/read quality, resulting in N=3 per genotype (2 male/1 female). Mapped reads were assigned to genes using featureCounts version 1.6.3 (Liao et al., 2014). High Pearson correlation (r > 0.99) was observed between all Vglut1^Cre/+^;DLK(cKO)^fl/fl^;Rpl22^HA/+^ or Vglut1^Cre/+;^Hipp11-DLK(iOE)/+;Rpl22^HA/+^ samples and their respective littermate controls. Differential gene expression analysis was conducted using DESeq2 galaxy version 2.11.40.2 (Love et al., 2014) with genotype, sex, and batch included as factors in the analysis. Generation of volcano plots was performed in RStudio version 1.2.1335 using the ggplot2 package version 3.3.5 (Wickham, 2016). Heatmaps were generated using the heatmap.2 function on Galaxy (Galaxy version 3.0.1) using normalized gene counts with a log2 transformation and scaling by row.

### GENE ONTOLOGY

Gene ontology analysis was performed using DAVID 2021 version (Huang et al., 2009) on genes found to be differentially expressed with p<0.05. For gene ontology and pathway analysis, background gene lists were generated by removing any gene with a base mean from DEseq2 normalization less than 1. *Gfap* was removed from GO and pathway analysis as a differentially expressed gene as it likely reflects a small amount of contamination from non-Vglut1 positive cells. DAVID analysis was performed using default thresholds, and Benjamini corrected p-values are reported. GO terms displayed in figures were chosen from top terms reaching significance related to biological processes or cellular components (BP5, CC4 or CC5) categories after filtering terms for semantic similarity. For SynGO analysis, mouse genes detected as differentially expressed were converted to human IDs using the ID conversion tool, and analysis was performed using the brain expressed background gene list provided by SynGO (Koopmans et al., 2019) (Version/release 20210225).

### RANK RANK HYPERGEOMETRIC OVERLAP (RRHO) ANALYSIS FOR CORRELATION OF GENE EXPRESSION PATTERNS

We used Rank Rank Hypergeometric overlap (https://systems.crump.ucla.edu/rankrank/rankranksimple.php) to compare DLK(iOE) and DLK(cKO) translatome datasets (Plaisier et al., 2010). Input gene lists included 12740 genes which were expressed across all samples. For each gene, the -Log10P_adj_ was multiplied by the sign of the fold change to obtain the metric used for ranking. Both DLK(iOE) and DLK(cKO) datasets were ranked in order to have increasing DLK along the x and y axis. RRHO was run using a step size of 100 genes. The Benjamini-Yekutieli corrected graph is shown.

### HIPPOCAMPAL SPATIAL EXPRESSION ANALYSIS

Comparison with gene expression databases: Gene expression patterns of differentially translated genes were evaluated using Allen Mouse Brain Atlas in situ data from P56 mice, and supplemented with data from Habib et al., 2016 through the Single cell portal from the Broad Institute or data from Zeisel et al., 2018, adolescent data through mousebrain.org when no in situ data was available or when expression was weak.

Comparison with CamK2-RiboTag and Grik4-RiboTag data: Gene set enrichment analysis (GSEA) (4.2.2) (Subramanian et al., 2005) was performed on Vglut1-RiboTag expression data after filtering lowly expressed genes using normalized counts. Analysis was conducted using the parameters: 1000 permutations, no collapse gene set, and permutation type gene set, with all other settings as default. To define gene sets for CA1 or CA3 enriched genes, we analyzed RiboTag datasets (Traunmüller et al., 2023; GSE209870) in wild type 6-week-old CA1 and CA3 neurons, from CamK2-cre and Grik4-cre mice, respectively. We compared the CamK2-RiboTag dataset and Grik4-RiboTag expression dataset to identify genes which were enriched in CA1 compared to CA3 or vice versa, applying an expression filter (average of at least 50 reads/animal) to ensure genes enriched in a particular region were expressed. The top 100 genes enriched in CamK2-RiboTag relative to Grik4-RiboTag were considered “CA1 genes”. The top 100 genes enriched in Grik4-RiboTag relative to CamK2-RiboTag were considered “CA3 genes”. 82 out of 100 GRIK4 (CA3) and 83 out of 100 CAMK2 (CA1) enriched genes were expressed in both our WT and DLKcKO samples (Supplementary excel File S3 CamK2 Grik4 enriched genes).

### RNASCOPE ANALYSIS

The RNAscope Fluorescent Multiplex Reagent kit (Cat. #320850) Amp 4 Alt A-FL(Wang et al., 2012) with probes from Advanced Cell Diagnostics were used. The protocol was carried out under RNase-free conditions and following the manufacturer’s instructions. Mice were anesthetized with isoflurane prior to decapitation. Brains were dissected immediately and flash frozen in OCT at -80°C. Fresh-frozen tissue was cryosectioned coronally to 20µm, collected on glass slides (Superfrost Plus), and stored at -80°C. Slides were fixed with 4% paraformaldehyde, dehydrated with 50% ethanol, 70% ethanol, and 2x washes in 100% ethanol for 5 min each at RT, followed by incubation in Protease IV reagent for 30 min at 40°C. Hybridization with target probes was performed at 40°C for 2 hours in a humidified slide box in an incubator followed by wash and amplification steps according to the manufacturer’s protocol. Finally, tissue was counterstained with DAPI, and mounted with Prolong diamond antifade mountant. All target probes were multiplexed with probes for *Vglut1* to label glutamatergic neurons.

### PRIMARY HIPPOCAMPAL NEURON CULTURES AND IMMUNOSTAINING

Prior to preparing cultures, Poly-D-Lysine (Corning, Cat#354210) was coated on 12mm glass coverslips (0.2 mg/mL) or 6-well plates (0.05mg/ml) for two days at 37°C. Neurons with indicated genotypes are labeled by tdTomato from Cre-dependent Rosa26-tdTomato generated from the following crosses: for control: *Vglut1^Cre/+^* X *Rosa26^tdT/+^*; for DLKcKO: *Vglut1^Cre/+^;DLK(cKO)^fl/fl^* X *Vglut1^Cre/+^;DLK(cKO)^fl/fl^;Rosa26^tdT/+^;* for DLKiOE: *H11-DLK^iOE/iOE^* X *Vglut1 ^Cre/+^;Rosa26^tdT/+^.* Primary neurons were generated from hippocampi of P1 pups. Mice were rapidly decapitated, then brains were removed, placed into ice cold HBSS (calcium- and magnesium-free) supplemented with 10mM HEPES for removal of meninges and dissection of hippocampi (Kaech and Banker, 2006). Dissected hippocampi were dissociated in HBSS with HEPES in 0.25% trypsin for 15 minutes at 37°C, and were then washed 3 times with 5ml of 20% Fetal bovine serum in HBSS. Dissociated cells from a litter were pooled into the same culture. Cells were triturated in Opti-MEM supplemented with 20mM glucose by five passes with an unpolished glass pipette and five to ten passes using a fire polished glass pipette. Cells were counted using a hemocytometer, and 60,000 cells were plated per coverslip into a 24-well plate or 300,000 per well of a 6-well dish. Cultures were kept in an incubator at 37°C with 5% CO2. After four hours, plating media was replaced with prewarmed Neurobasal Medium supplemented with glutamine, penicillin/streptomycin, and B27. Cells were fixed after 48 hours (DIV2) or on DIV14 with prewarmed 4% PFA/4% sucrose in PBS for 20 min at room temperature followed by 3 washes with PBS. Media were changed carefully to minimize impacts to growth cone morphology.

Staining of fixed neurons was performed in 24-well plates. Coverslips were incubated in 50mM ammonium chloride for 10 min, followed by 3 washes PBS, 5 min 0.1% Triton X-100 in PBS, and blocking in 30mg/ml Bovine serum albumin (BSA) in 0.1% Triton in PBS for 30 min. Coverslips were incubated in primary antibody diluted in 30mg/ml BSA in 0.1% Triton in PBS according to antibody table for 90 minutes at room temperature followed by four washes in 0.1% Triton in PBS. Secondary antibodies were diluted in 30mg/ml BSA in 0.1% Triton in PBS with 1% donkey serum according to antibody table, and incubated for 60 minutes at room temperature. Finally, coverslips were washed three times in 0.2% Triton in PBS, stained with DAPI, washed three times with PBS, and mounted using Prolong Diamond Antifade Mountant. For an unknown reason, co-immunostaining of DLK and tyrosinated tubulin led to a pattern of DLK staining different from the punctate appearance of DLK observed in other conditions. The typical appearance of DLK could still be observed in cells with high levels of DLK. This altered appearance was not observed during co-immunostaining of DLK and acetylated tubulin.

### CONFOCAL IMAGING AND QUANTIFICATION

Fluorescent images were acquired using a Zeiss LSM800 confocal microscope using a 10x, 20x, or 63x objective. All tissue sections and neurons within the same experiment were imaged under identical conditions. For brain tissue, three sections per mouse were imaged with a minimum of three mice per genotype for data analysis. Dorsal hippocampal images were taken from approximately bregma -1.5mm to -2.3mm. For image analysis, the quantification was performed blind to genotype or in an automated manner when possible. All image processing and analysis was performed using Fiji distribution of ImageJ unless otherwise specified (Schindelin et al., 2012).

For quantification of mRNA puncta, ROI were drawn to count puncta overlapping with nuclei of *Vglut1* positive cells. Individual puncta were counted from >50 cells per genotype in a blinded manner. Puncta counts were normalized to *Vglut1* puncta counts to control for variability in staining or preservation of RNA. Three to four sections per mouse were quantified and three mice per genotype were stained with each probe.

Pyramidal cell layer thickness was measured across CA1 by averaging the lengths of three perpendicular lines extending across the maximum projected z-stack of the pyramidal cell layer for each section. Three sections were averaged per mouse from dorsal hippocampus. For sections including ventral hippocampus, cell layer thickness of CA1 was measured using three lines either above the ventral edge of the suprapyramidal blade of dentate gyrus (Dong et al., 2009) for dorsal hippocampus (posterior) quantifications or below the ventral edge of the DG for ventral CA1 quantifications. Hippocampal cross-sectional area was measured by tracing outlines of CA1, CA3, or DG (including dendritic layers) in dorsal hippocampus sections.

DLK signal intensity in immunofluorescence images was quantified by drawing an outline around CA1, CA3, or DG (all cell layers), and measuring the mean fluorescence intensity.

Tuj1, tyrosinated tubulin, acetylated tubulin, and MAP2 intensities were measured using the mean gray value from auto thresholding (default) over stratum radiatum of CA1, the molecular layer of DG, or stratum lacunosum-moleculare, stratum radiatum, and stratum lucidum of CA3.

Staining of p-c-Jun in conditional knockout animals and c-Jun intensities were quantified from 20x images using mean gray values of ROIs for each brain slice cropped around the pyramidal cell or granule cell layers with background subtraction of non-nuclear signal from dendritic regions. Analysis of p-c-Jun positive nuclei in DLK(iOE) mice was counted from 10x images with using an intensity threshold of 20000 (P10) or 110 or 140 (P15) depending on imaging conditions. Nuclei were separated using a watershed, and all nuclei larger than 10µm^2^ were counted.

TUNEL positive signals were counted as fluorescent signals overlapping with the pyramidal cell or dentate granule cell layer in each region from 10x tile scan images of dorsal hippocampus. Z-stacks covering the entire section were max projected for quantification.

VGLUT1 (protein), Bassoon, and Homer1 puncta were quantified from stratum radiatum of dorsal CA1. Images were quantified using a single slice image, and a 25×25µm ROI was chosen to minimize absence of puncta due to cell bodies. A gaussian filter of 1 pixel was applied to the image. Background subtraction was performed using a rolling ball radius of 10 pixels, and an automated threshold was applied to the image using the Otsu method followed by a watershed to separate clustered puncta. Puncta larger than 2 pixels were counted for individual proteins. Overlap of Bassoon and Homer1 puncta of any size were counted. The number and average size of puncta were recorded from two images per brain section and three sections per mouse.

GFAP mean fluorescence intensity was quantified in a 312µm x 312µm box around the pyramidal cell or granule cell layers of CA1, CA3, and DG with background intensity subtracted after measuring from an area without GFAP signal.

Neurons were selected for neurite outgrowth and axon analysis after confirming DLK protein level by antibody staining and measurement of DLK fluorescence intensity in cell soma at DIV2. While we used tdTomato as a reporter for Vglut1 positive neurons, not all tdTomato positive neurons showed detectable differences in DLK levels at this early (DIV2) timepoint. Cell somas were outlined using tdTomato, and DLK integrated density was measured. Integrated density reflects the mean gray value multiplied by the area. Vglut1^Cre/+^ control cells were selected for further analysis if DLK integrated density was 4,000-8,500. Cells from DLK(cKO) cultures with integrated density values of DLK less than 4,000 were selected for further analysis as “DLK(cKO)” and cells from DLK(iOE) with integrated density values of greater than 8,500 were selected for further analysis as “DLK(iOE)”. While we observed variably increased DLK signals in DLK(iOE) neurons from moderate to strong increases, all DLK(iOE) neurons with increased levels above the set threshold were grouped together in quantifications due to limited numbers of neurons. Primary neurites were counted in a blinded manner from tdTomato channel, counting both branches and filopodia originating from cell soma region. Neurites were considered as axons in axon specification analysis if longer than 90µm.

Bassoon puncta in cell culture were quantified from 20µm stretches of neurite. Regions for analysis were selected based on tdTomato positive signal on thin processes without dendritic spines exhibiting Bassoon signal, that was not in a region densely populated by Bassoon signal from other neurites. DLK levels were also used to select ROIs. Signal from tdTomato was used to create a 10 pixel ROI along the neurite. Bassoon puncta were identified in a blinded manner by smoothing the image, applying a triangle threshold, and manually dividing merged puncta based on bassoon intensity. All puncta 5 pixels or larger and overlapping with tdTomato signal were analyzed for puncta size and density.

Dendritic spines were quantified from a 20µm countable and representative stretch of dendrite within 75µm of the neuron soma from one of the three largest dendrites. Spines density was counted using tdTomato signal, and calculated by counting total dendritic spines divided by the traced length of dendrite. Spines were manually categorized following measurements in Risher et al., 2014. Filopodia (>2µm) are not included in spine density counts. Spines were quantified from 3 independent cultures per genotype with 8-16 neurons per culture.

### STASTICAL ANALYSIS

All statistical analysis shown in graphs was performed using GraphPad Prism 9.4.0. Points represent individual values, with bars reflecting mean values, and error bars plotting standard error of the mean (SEM).

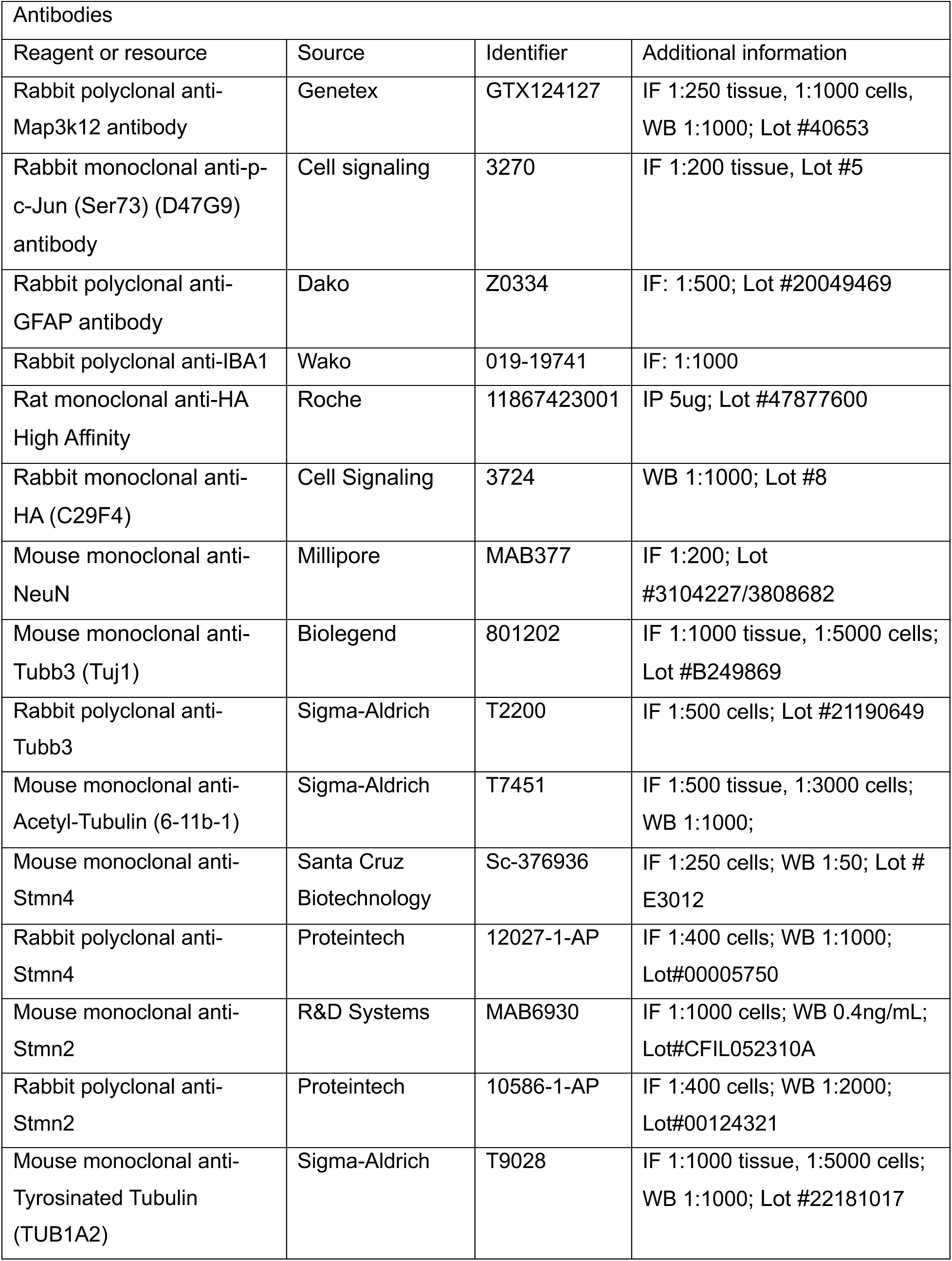

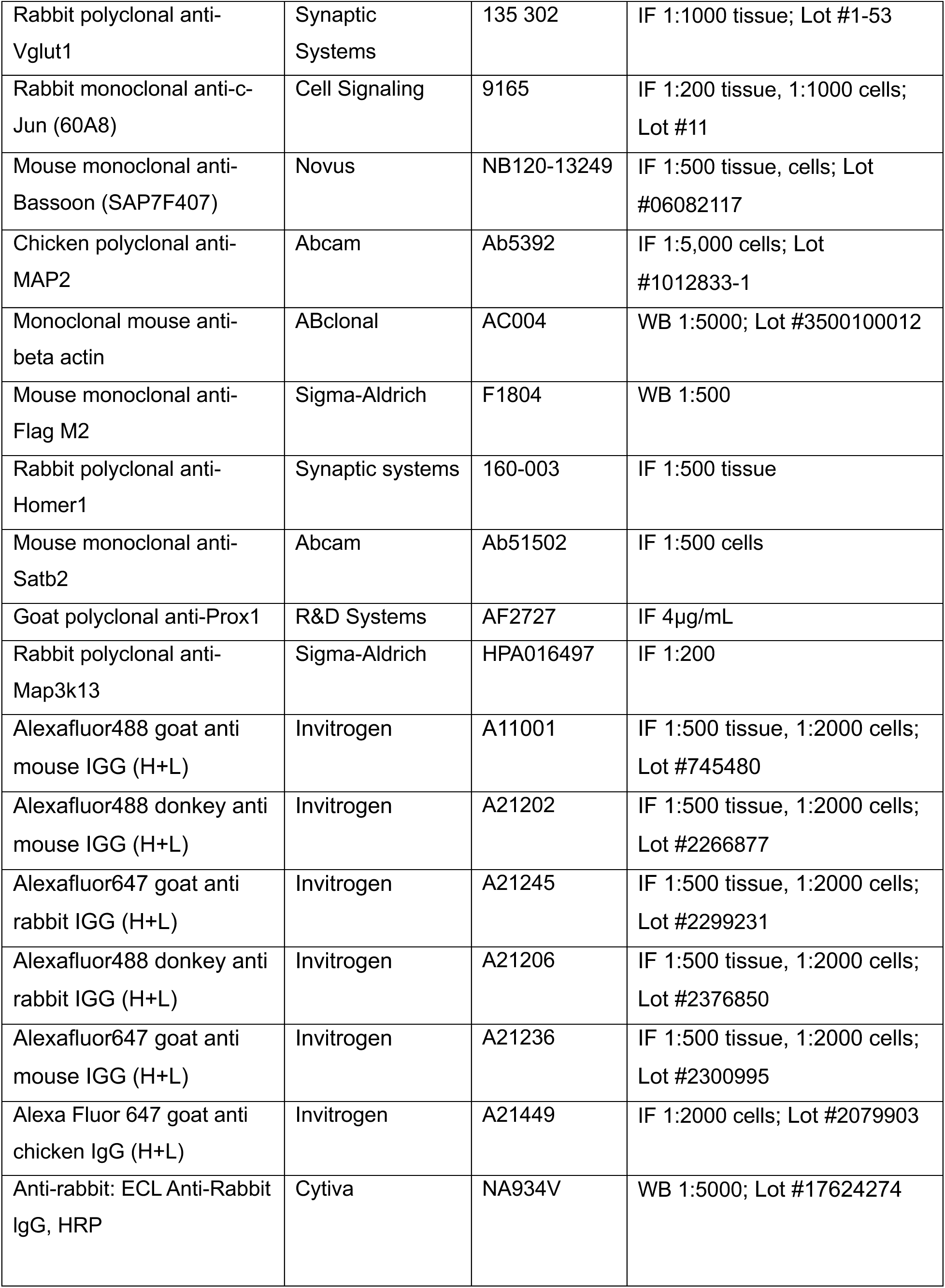

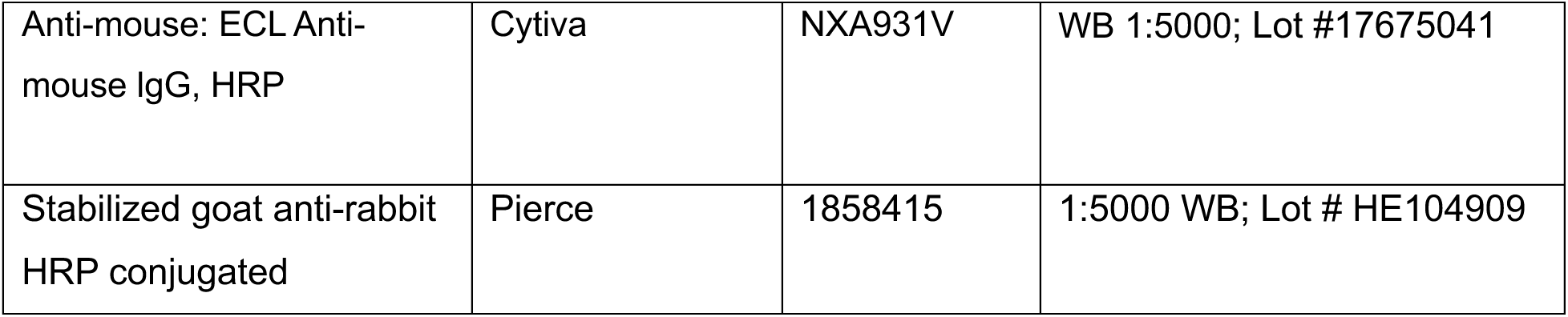

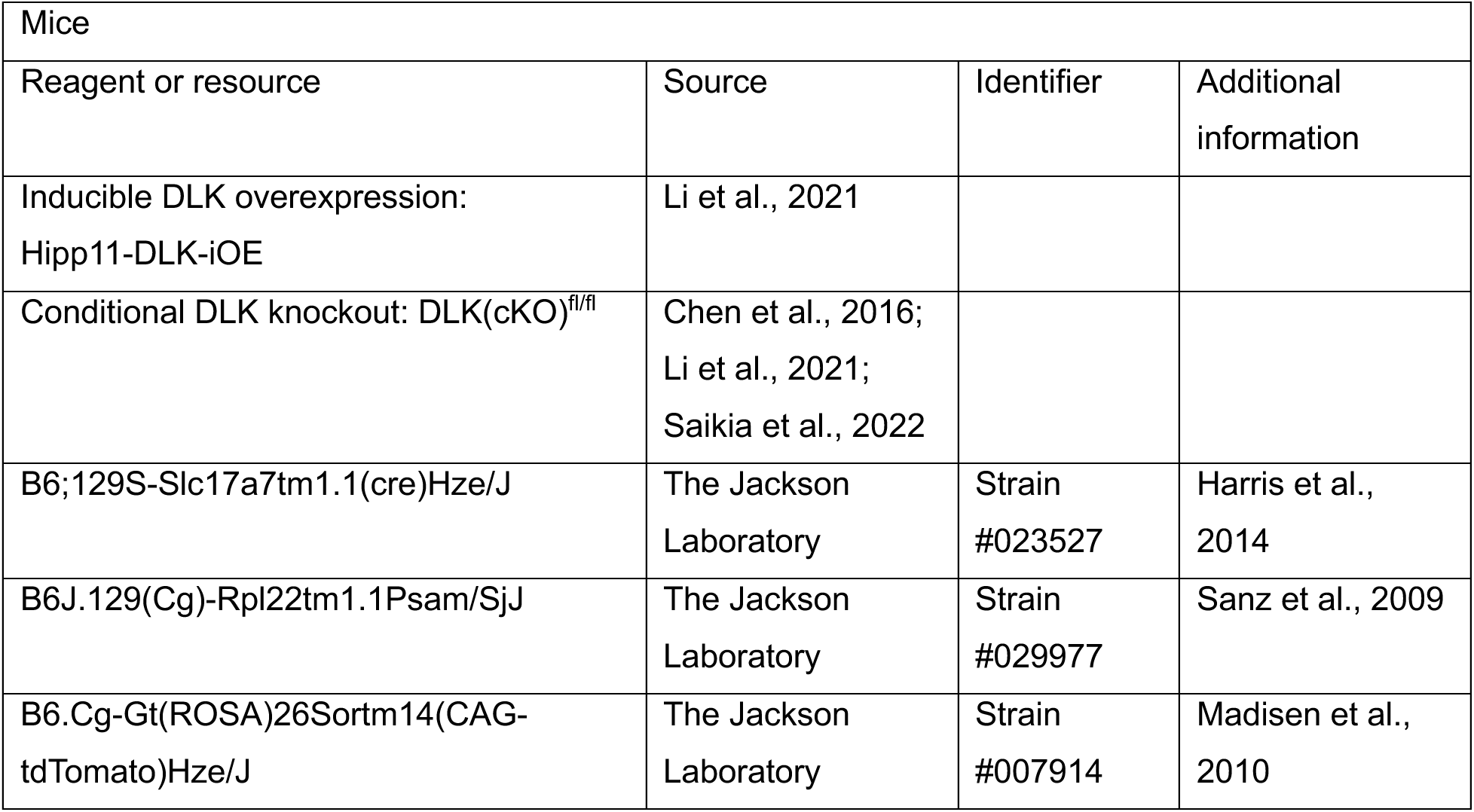

| Mice |  |  |  |
| --- | --- | --- | --- |
| Reagent or resource | Source | Identifier | Additional information |
| Inducible DLK overexpression: Hipp11-DLK-iOE | Li et al., 2021 |  |  |
| Conditional DLK knockout: DLK(cKO)<sup>fl/fl</sup> | Chen et al., 2016; Li et al., 2021; Saikia et al., 2022 |  |  |
| B6;129S-Slc17a7tm1.1(cre)Hze/J | The Jackson Laboratory | Strain #023527 | Harris et al., 2014 |
| B6J.129(Cg)-Rpl22tm1.1Psam/SjJ | The Jackson Laboratory | Strain #029977 | Sanz et al., 2009 |
| B6.Cg-Gt(ROSA)26Sortm14(CAG-tdTomato)Hze/J | The Jackson Laboratory | Strain #007914 | Madisen et al., 2010 |

| RNA probes |  |  |
| --- | --- | --- |
| Reagent or resource | Source | Identifier |
| RNAscope probe MAP3K12-C2 | ACD | 458151-C2 |
| RNAscope probe Slc17a7-C3 | ACD | 416631-C3 |
| RNAscope probe Gfap-C2 | ACD | 313211-C2 |
| RNAscope probe Stmn4 | ACD | 537541 |

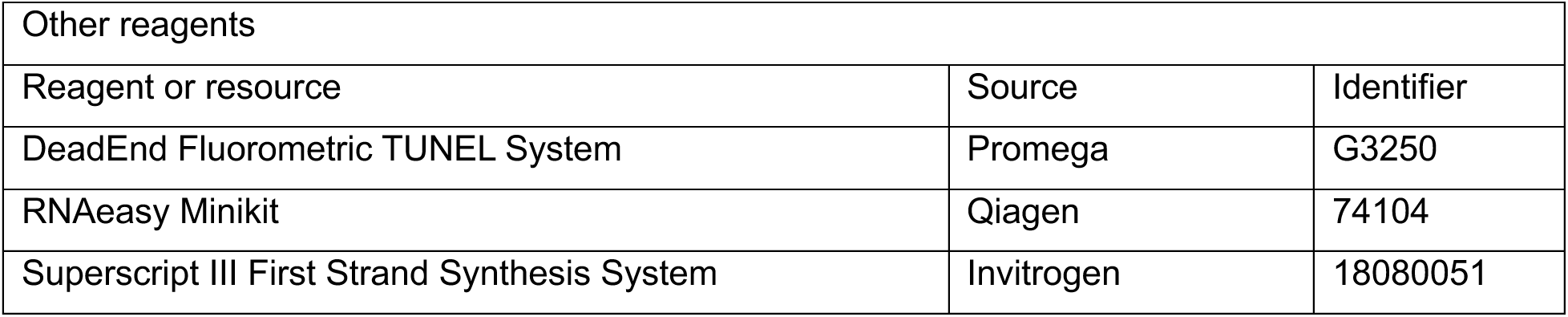

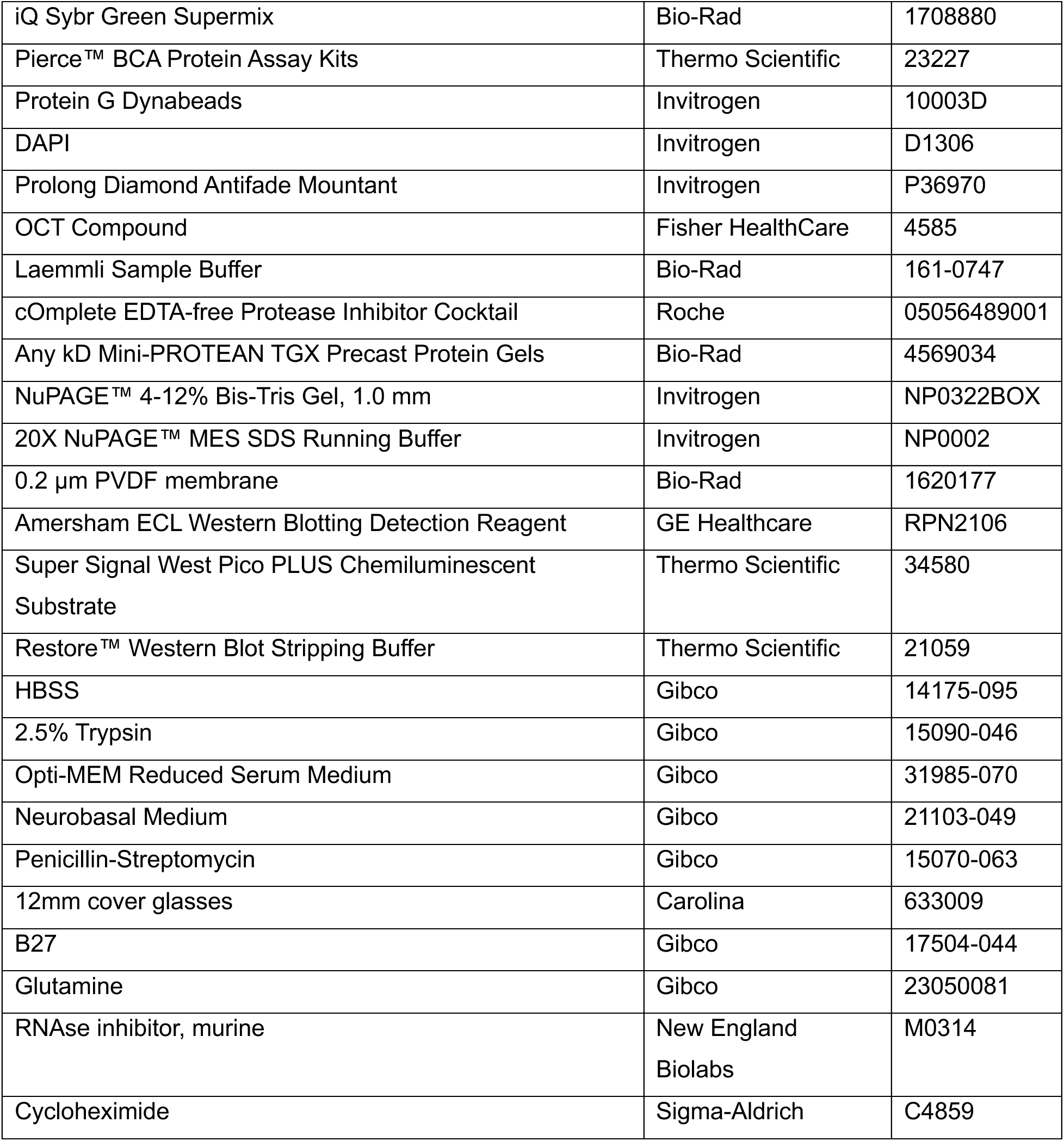

## Notes

### Competing Interest Statement

The authors have declared no competing interest.

### Summary of Updates

This version of the manuscript includes updated Fig 1, Fig 4, Fig 5, three new supplemental figures (Fig S2, S11, S12), and updated Fig S1, S3, S5, S6, S7, S8, S13, along with associated figure legends and text revision. Dilan Acar is added as an author, her contribution is described in the manuscript.

## References

Afgan E, Baker D, Batut B, van den Beek M, Bouvier D, Čech M, Chilton J, Clements D, Coraor N, Grüning BA, Guerler A, Hillman-Jackson J, Hiltemann S, Jalili V, Rasche H, Soranzo N, Goecks J, Taylor J, Nekrutenko A, Blankenberg D. 2018. The Galaxy platform for accessible, reproducible and collaborative biomedical analyses: 2018 update. Nucleic Acids Res 46:W537–W544.

Agrimi G, Russo A, Scarcia P, Palmieri F. 2012. The human gene SLC25A17 encodes a peroxisomal transporter of coenzyme A, FAD and NAD+. Biochem J 443:241–247. doi:10.1042/BJ20111420

An integrated encyclopedia of DNA elements in the human genome. 2012. . Nature 489:57–74. doi:10.1038/nature11247

Angel P, Hattori K, Smeal T, Karin M. 1988. The jun proto-oncogene is positively autoregulated by its product, Jun/AP-1. Cell 55:875–885. 10.1016/0092-8674(88)90143-2

Asghari Adib E, Shadrach JL, Reilly-Jankowiak L, Dwivedi MK, Rogers AE, Shahzad S, Passino R, Giger RJ, Pierchala BA, Collins CA. 2024. DLK signaling in axotomized neurons triggers complement activation and loss of upstream synapses. Cell Rep 43. doi:10.1016/j.celrep.2024.113801

Asghari Adib E, Smithson LJ, Collins CA. 2018. An axonal stress response pathway: degenerative and regenerative signaling by DLK. Curr Opin Neurobiol 53:110–119. doi:10.1016/j.conb.2018.07.002

Behrens A, Sibilia M, Wagner EF. 1999. Amino-terminal phosphorylation of c-Jun regulates stress-induced apoptosis and cellular proliferation. Nat Genet 21:326–329. doi:10.1038/6854

Blondeau A, Lucier JF, Matteau D, Dumont L, Rodrigue S, Jacques PÉ, Blouin R. 2016. Dual leucine zipper kinase regulates expression of axon guidance genes in mouse neuronal cells. Neural Dev 11:1–12. doi:10.1186/s13064-016-0068-8

Blouin R. 1996. Cell-specific expression of the ZPK gene in adult mouse tissues. DNA Cell Biol 15:631–642. doi:10.1089/dna.1996.15.631

Charbaut E, Curmi PA, Ozon S, Lachkar S, Redeker V, Sobel A. 2001. Stathmin Family Proteins Display Specific Molecular and Tubulin Binding Properties. J Biol Chem 276:16146–16154. doi:10.1074/jbc.M010637200

Chauvin S, Poulain FE, Ozon S, Sobel A. 2008. Palmitoylation of stathmin family proteins domain A controls Golgi versus mitochondrial subcellular targeting. Biol Cell 100:577–591. doi:10.1042/bc20070119

Chauvin S, Sobel A. 2015. Neuronal stathmins: A family of phosphoproteins cooperating for neuronal development, plasticity and regeneration. Prog Neurobiol 126:1–18. doi:10.1016/j.pneurobio.2014.09.002

Chen M, Geoffroy CG, Wong HN, Tress O, Nguyen MT, Holzman LB, Jin Y, Zheng B. 2016. Leucine Zipper-bearing Kinase promotes axon growth in mammalian central nervous system neurons. Sci Rep 6:1–16. doi:10.1038/srep31482

Collins CA, Wairkar YP, Johnson SL, DiAntonio A. 2006. Highwire Restrains Synaptic Growth by Attenuating a MAP Kinase Signal. Neuron 51:57–69. doi:10.1016/j.neuron.2006.05.026

DeVault L, Mateusiak C, Palucki J, Brent M, Milbrandt J, DiAntonio A. 2024. The response of Dual-leucine zipper kinase (DLK) to nocodazole: Evidence for a homeostatic cytoskeletal repair mechanism. PLoS One 19:e0300539.

Dobin A, Davis CA, Schlesinger F, Drenkow J, Zaleski C, Jha S, Batut P, Chaisson M, Gingeras TR. 2013. STAR: Ultrafast universal RNA-seq aligner. Bioinformatics 29:15–21. doi:10.1093/bioinformatics/bts635

Dodelet VC, Pazzagli C, Zisch AH, Hauser CA, Pasquale EB. 1999. A Novel Signaling Intermediate, SHEP1, Directly Couples Eph Receptors to R-Ras and Rap1A *. J Biol Chem 274:31941–31946. doi:10.1074/jbc.274.45.31941

Dong HW, Swanson LW, Chen L, Fanselow MS, Toga AW. 2009. Genomic-anatomic evidence for distinct functional domains in hippocampal field CA1. Proc Natl Acad Sci U S A 106:11794–11799. doi:10.1073/pnas.0812608106

Dotti CG, Sullivan CA, Banker GA. 1988. The establishment of polarity by hippocampal neurons in culture. J Neurosci 8:1454 LP – 1468. doi:10.1523/JNEUROSCI.08-04-01454.1988

Eilers A, Whitfield J, Babij C, Rubin LL, Ham J. 1998. Role of the Jun Kinase Pathway in the Regulation of c-Jun Expression and Apoptosis in Sympathetic Neurons. J Neurosci 18:1713 LP – 1724. doi:10.1523/JNEUROSCI.18-05-01713.1998

Furlanis E, Traunmüller L, Fucile G, Scheiffele P. 2019. Landscape of ribosome-engaged transcript isoforms reveals extensive neuronal-cell-class-specific alternative splicing programs. Nat Neurosci 22:1709–1717. doi:10.1038/s41593-019-0465-5

Gavet O, Messari S El, Ozon S, Sobel A. 2002. Regulation and subcellular localization of the microtubule-destabilizing stathmin family phosphoproteins in cortical neurons. J Neurosci Res 68:535–550. doi:10.1002/jnr.10234

Ghosh AS, Wang B, Pozniak CD, Chen M, Watts RJ, Lewcock JW. 2011. DLK induces developmental neuronal degeneration via selective regulation of proapoptotic JNK activity. J Cell Biol 194:751–764. doi:10.1083/jcb.201103153

Gómez de San José N, Massa F, Halbgebauer S, Oeckl P, Steinacker P, Otto M. 2022. Neuronal pentraxins as biomarkers of synaptic activity: from physiological functions to pathological changes in neurodegeneration. J Neural Transm 129:207–230. doi:10.1007/s00702-021-02411-2

Goodwani S, Fernandez C, Acton PJ, Buggia-Prevot V, McReynolds ML, Ma J, Hu CH, Hamby ME, Jiang Y, Le K, Soth MJ, Jones P, Ray WJ. 2020. Dual leucine zipper kinase is constitutively active in the adult mouse brain and has both stress-induced and homeostatic functions. Int J Mol Sci 21:1–24. doi:10.3390/ijms21144849

Habib N, Li Y, Heidenreich M, Swiech L, Avraham-Davidi I, Trombetta JJ, Hession C, Zhang F, Regev A. 2016. Div-Seq: Single-nucleus RNA-Seq reveals dynamics of rare adult newborn neurons. Science 353:925–928. doi:10.1126/science.aad7038

Harris JA, Hirokawa KE, Sorensen SA, Gu H, Mills M, Ng LL, Bohn P, Mortrud M, Ouellette B, Kidney J, Smith KA, Dang C, Sunkin S, Bernard A, Oh SW, Madisen L, Zeng H. 2014. Anatomical characterization of Cre driver mice for neural circuit mapping and manipulation. Front Neural Circuits 8:76. doi:10.3389/fncir.2014.00076

Hirai S, Banba Y, Satake T, Ohno S. 2011. Axon Formation in Neocortical Neurons Depends on Stage-Specific Regulation of Microtubule Stability by the Dual Leucine Zipper Kinase–c-Jun N-Terminal Kinase Pathway. J Neurosci 31:6468 LP – 6480. doi:10.1523/JNEUROSCI.5038-10.2011

Hirai S, Kawaguchi A, Hirasawa R, Baba M, Ohnishi T, Ohno S. 2002. MAPK-upstream protein kinase (MUK) regulates the radial migration of immature neurons in telencephalon of mouse embryo. Development 129:4483–4495. doi:10.1242/dev.129.19.4483

Hirai S, Kawaguchi A, Suenaga J, Ono M, Cui DF, Ohno S. 2005. Expression of MUK/DLK/ZPK, an activator of the JNK pathway, in the nervous systems of the developing mouse embryo. Gene Expr Patterns 5:517–523. 10.1016/j.modgep.2004.12.002

Hirai SI, De FC, Miyata T, Ogawa M, Kiyonari H, Suda Y, Aizawa S, Banba Y, Ohno S. 2006. The c-Jun N-terminal kinase activator dual leucine zipper kinase regulates axon growth and neuronal migration in the developing cerebral cortex. J Neurosci 26:11992–12002. doi:10.1523/JNEUROSCI.2272-06.2006

Holland SM, Collura KM, Ketschek A, Noma K, Ferguson TA, Jin Y, Gallo G, Thomas GM. 2016. Palmitoylation controls DLK localization, interactions and activity to ensure effective axonal injury signaling. Proc Natl Acad Sci U S A 113:763–768. doi:10.1073/pnas.1514123113

Hu Z, Deng N, Liu K, Zeng W. 2019. DLK mediates the neuronal intrinsic immune response and regulates glial reaction and neuropathic pain. Exp Neurol 322:113056. doi:10.1016/j.expneurol.2019.113056

Huang DW, Sherman BT, Lempicki RA. 2009. Systematic and integrative analysis of large gene lists using DAVID bioinformatics resources. Nat Protoc 4:44–57. doi:10.1038/nprot.2008.211

Huang Y-WA, Zhou B, Nabet AM, Wernig M, Südhof TC. 2019. Differential Signaling Mediated by ApoE2, ApoE3, and ApoE4 in Human Neurons Parallels Alzheimer’s Disease Risk. J Neurosci 39:7408–7427. doi:10.1523/JNEUROSCI.2994-18.2019

Huang Y-WA, Zhou B, Wernig M, Südhof TC. 2017. ApoE2, ApoE3, and ApoE4 Differentially Stimulate APP Transcription and Aβ Secretion. Cell 168:427–441.e21. doi:10.1016/j.cell.2016.12.044

Huntwork-Rodriguez S, Wang B, Watkins T, Ghosh AS, Pozniak CD, Bustos D, Newton K, Kirkpatrick DS, Lewcock JW. 2013. JNK-mediated phosphorylation of DLK suppresses its ubiquitination to promote neuronal apoptosis. J Cell Biol 202:747–763. doi:10.1083/jcb.201303066

Itoh A, Horiuchi M, Bannerman P, Pleasure D, Itoh T. 2009. Impaired regenerative response of primary sensory neurons in ZPK/DLK gene-trap mice. Biochem Biophys Res Commun 383:258–262. doi:10.1016/j.bbrc.2009.04.009

Itoh A, Horiuchi M, Wakayama K, Xu J, Bannerman P, Pleasure D, Itoh T. 2011. ZPK/DLK, a mitogen-activated protein kinase kinase kinase, is a critical mediator of programmed cell death of motoneurons. J Neurosci 31:7223–7228. doi:10.1523/JNEUROSCI.5947-10.2011

Iwano T, Masuda A, Kiyonari H, Enomoto H, Matsuzaki F. 2012. Prox1 postmitotically defines dentate gyrus cells by specifying granule cell identity over CA3 pyramidal cell fate in the hippocampus. Development 139:3051–3062. doi:10.1242/dev.080002

Janke C, Magiera MM. 2020. The tubulin code and its role in controlling microtubule properties and functions. Nat Rev Mol Cell Biol 21:307–326. doi:10.1038/s41580-020-0214-3

Jin Y, Zheng B. 2019. Multitasking: Dual Leucine Zipper-Bearing Kinases in Neuronal Development and Stress Management. Annu Rev Cell Dev Biol 35:501–521. doi:10.1146/annurev-cellbio-100617-062644

Joy MT, Ben Assayag E, Shabashov-Stone D, Liraz-Zaltsman S, Mazzitelli J, Arenas M, Abduljawad N, Kliper E, Korczyn AD, Thareja NS, Kesner EL, Zhou M, Huang S, Silva TK, Katz N, Bornstein NM, Silva AJ, Shohami E, Carmichael ST. 2019. CCR5 Is a Therapeutic Target for Recovery after Stroke and Traumatic Brain Injury. Cell 176:1143–1157.e13. doi:10.1016/j.cell.2019.01.044

Kaech S, Banker G. 2006. Culturing hippocampal neurons. Nat Protoc 1:2406–2415. doi:10.1038/nprot.2006.356

Koopmans F, van Nierop P, Andres-Alonso M, Byrnes A, Cijsouw T, Coba MP, Cornelisse LN, Farrell RJ, Goldschmidt HL, Howrigan DP, Hussain NK, Imig C, de Jong APH, Jung H, Kohansalnodehi M, Kramarz B, Lipstein N, Lovering RC, MacGillavry H, Mariano V, Mi H, Ninov M, Osumi-Sutherland D, Pielot R, Smalla K-H, Tang H, Tashman K, Toonen RFG, Verpelli C, Reig-Viader R, Watanabe K, van Weering J, Achsel T, Ashrafi G, Asi N, Brown TC, De Camilli P, Feuermann M, Foulger RE, Gaudet P, Joglekar A, Kanellopoulos A, Malenka R, Nicoll RA, Pulido C, de Juan-Sanz J, Sheng M, Südhof TC, Tilgner HU, Bagni C, Bayés À, Biederer T, Brose N, Chua JJE, Dieterich DC, Gundelfinger ED, Hoogenraad C, Huganir RL, Jahn R, Kaeser PS, Kim E, Kreutz MR, McPherson PS, Neale BM, O’Connor V, Posthuma D, Ryan TA, Sala C, Feng G, Hyman SE, Thomas PD, Smit AB, Verhage M. 2019. SynGO: An Evidence-Based, Expert-Curated Knowledge Base for the Synapse. Neuron 103:217–234.e4. doi:10.1016/j.neuron.2019.05.002

Larhammar M, Huntwork-Rodriguez S, Jiang Z, Solanoy H, Ghosh AS, Wang B, Kaminker JS, Huang K, Eastham-Anderson J, Siu M, Modrusan Z, Farley MM, Tessier-Lavigne M, Lewcock JW, Watkins TA. 2017. Dual leucine zipper kinase-dependent PERK activation contributes to neuronal degeneration following insult. Elife 6:1–27. doi:10.7554/eLife.20725

Le Pichon CE, Meilandt WJ, Dominguez S, Solanoy H, Lin H, Ngu H, Gogineni A, Ghosh AS, Jiang Z, Lee SH, Maloney J, Gandham VD, Pozniak CD, Wang B, Lee S, Siu M, Patel S, Modrusan Z, Liu X, Rudhard Y, Baca M, Gustafson A, Kaminker J, Carano RAD, Huang EJ, Foreman O, Weimer R, Scearce-Levie K, Lewcock JW. 2017. Loss of dual leucine zipper kinase signaling is protective in animal models of neurodegenerative disease. Sci Transl Med 9. doi:10.1126/scitranslmed.aag0394

Lein ES, Hawrylycz MJ, Ao N, Ayres M, Bensinger A, Bernard A, Boe AF, Boguski MS, Brockway KS, Byrnes EJ, Chen Lin, Chen Li, Chen T-M, Chi Chin M, Chong J, Crook BE, Czaplinska A, Dang CN, Datta S, Dee NR, Desaki AL, Desta T, Diep E, Dolbeare TA, Donelan MJ, Dong H-W, Dougherty JG, Duncan BJ, Ebbert AJ, Eichele G, Estin LK, Faber C, Facer BA, Fields R, Fischer SR, Fliss TP, Frensley C, Gates SN, Glattfelder KJ, Halverson KR, Hart MR, Hohmann JG, Howell MP, Jeung DP, Johnson RA, Karr PT, Kawal R, Kidney JM, Knapik RH, Kuan CL, Lake JH, Laramee AR, Larsen KD, Lau C, Lemon TA, Liang AJ, Liu Y, Luong LT, Michaels J, Morgan JJ, Morgan RJ, Mortrud MT, Mosqueda NF, Ng LL, Ng R, Orta GJ, Overly CC, Pak TH, Parry SE, Pathak SD, Pearson OC, Puchalski RB, Riley ZL, Rockett HR, Rowland SA, Royall JJ, Ruiz MJ, Sarno NR, Schaffnit K, Shapovalova N V, Sivisay T, Slaughterbeck CR, Smith SC, Smith KA, Smith BI, Sodt AJ, Stewart NN, Stumpf K-R, Sunkin SM, Sutram M, Tam A, Teemer CD, Thaller C, Thompson CL, Varnam LR, Visel A, Whitlock RM, Wohnoutka PE, Wolkey CK, Wong VY, Wood M, Yaylaoglu MB, Young RC, Youngstrom BL, Feng Yuan X, Zhang B, Zwingman TA, Jones AR. 2007. Genome-wide atlas of gene expression in the adult mouse brain. Nature 445:168–176. doi:10.1038/nature05453

Lein ES, Zhao X, Gage FH. 2004. Defining a Molecular Atlas of the Hippocampus Using DNA Microarrays and High-Throughput *In Situ* Hybridization. J Neurosci 24:3879 LP – 3889. doi:10.1523/JNEUROSCI.4710-03.2004

Lewcock JW, Genoud N, Lettieri K, Pfaff SL. 2007. The Ubiquitin Ligase Phr1 Regulates Axon Outgrowth through Modulation of Microtubule Dynamics. Neuron 56:604–620. doi:10.1016/j.neuron.2007.09.009

Li Y, Ritchie EM, Steinke CL, Qi C, Chen L, Zheng B, Jin Y. 2021. Activation of MAP3K DLK and LZK in Purkinje cells causes rapid and slow degeneration depending on signaling strength. Elife 10:e63509. doi:10.7554/eLife.63509

Liao Y, Smyth GK, Shi W. 2014. FeatureCounts: An efficient general purpose program for assigning sequence reads to genomic features. Bioinformatics 30:923–930. doi:10.1093/bioinformatics/btt656

Love MI, Huber W, Anders S. 2014. Moderated estimation of fold change and dispersion for RNA-seq data with DESeq2. Genome Biol 15:550. doi:10.1186/s13059-014-0550-8

Luo Y, Hitz BC, Gabdank I, Hilton JA, Kagda MS, Lam B, Myers Z, Sud P, Jou J, Lin K, Baymuradov UK, Graham K, Litton C, Miyasato SR, Strattan JS, Jolanki O, Lee J-W, Tanaka FY, Adenekan P, O’Neill E, Cherry JM. 2020. New developments on the Encyclopedia of DNA Elements (ENCODE) data portal. Nucleic Acids Res 48:D882–D889. doi:10.1093/nar/gkz1062

Lutjens R, Igarashi M, Pellier V, Blasey H, Di Paolo G, Ruchti E, Pfulg C, Staple JK, Catsicas S, Grenningloh G. 2000. Localization and targeting of SCG10 to the trans-Golgi apparatus and growth cone vesicles. Eur J Neurosci 12:2224–2234. 10.1046/j.1460-9568.2000.00112.x

Madisen L, Zwingman TA, Sunkin SM, Oh SW, Zariwala HA, Gu H, Ng LL, Palmiter RD, Hawrylycz MJ, Jones AR, Lein ES, Zeng H. 2010. A robust and high-throughput Cre reporting and characterization system for the whole mouse brain. Nat Neurosci 13:133–140. doi:10.1038/nn.2467

Mata M, Merritt SE, Fan G, Yu GG, Holzman LB. 1996. Characterization of dual leucine zipper-bearing kinase, a mixed lineage kinase present in synaptic terminals whose phosphorylation state is regulated by membrane depolarization via calcineurin. J Biol Chem 271:16888–16896. doi:10.1074/jbc.271.28.16888

Mielke K, Brecht S, Dorst A, Herdegen T. 1999. Activity and expression of JNK1, p38 and ERK kinases, c-Jun N-terminal phosphorylation, and c-jun promoter binding in the adult rat brain following kainate-induced seizures. Neuroscience 91:471–483. 10.1016/S0306-4522(98)00667-8

Mullen RJ, Buck CR, Smith AM. 1992. NeuN, a neuronal specific nuclear protein in vertebrates. Development 116:201–211. doi:10.1242/dev.116.1.201

Nakata K, Abrams B, Grill B, Goncharov A, Huang X, Chisholm AD, Jin Y. 2005. Regulation of a DLK-1 and p38 MAP kinase pathway by the ubiquitin ligase RPM-1 is required for presynaptic development. Cell 120:407–420. doi:10.1016/j.cell.2004.12.017

Nielsen J V, Blom JB, Noraberg J, Jensen NA. 2010. Zbtb20-Induced CA1 Pyramidal Neuron Development and Area Enlargement in the Cerebral Midline Cortex of Mice. Cereb Cortex 20:1904–1914. doi:10.1093/cercor/bhp261

Nihalani D, Merritt S, Holzman LB. 2000. Identification of structural and functional domains in mixed lineage kinase dual leucine zipper-bearing kinase required for complex formation and stress-activated protein kinase activation. J Biol Chem 275:7273–7279. doi:10.1074/jbc.275.10.7273

Paolicelli RC, Bolasco G, Pagani F, Maggi L, Scianni M, Panzanelli P, Giustetto M, Ferreira TA, Guiducci E, Dumas L, Ragozzino D, Gross CT. 2011. Synaptic Pruning by Microglia Is Necessary for Normal Brain Development. Science (80- ) 333:1456–1458. doi:10.1126/science.1202529

Plaisier SB, Taschereau R, Wong JA, Graeber TG. 2010. Rank–rank hypergeometric overlap: identification of statistically significant overlap between gene-expression signatures. Nucleic Acids Res 38:e169–e169. doi:10.1093/nar/gkq636

Pozniak CD, Sengupta Ghosh A, Gogineni A, Hanson JE, Lee S-H, Larson JL, Solanoy H, Bustos D, Li H, Ngu H, Jubb AM, Ayalon G, Wu J, Scearce-Levie K, Zhou Q, Weimer RM, Kirkpatrick DS, Lewcock JW. 2013. Dual leucine zipper kinase is required for excitotoxicity-induced neuronal degeneration. J Exp Med 210:2553–2567. doi:10.1084/jem.20122832

Risher WC, Ustunkaya T, Singh Alvarado J, Eroglu C. 2014. Rapid Golgi Analysis Method for Efficient and Unbiased Classification of Dendritic Spines. PLoS One 9:e107591.

Roussarie J-P, Yao V, Rodriguez-Rodriguez P, Oughtred R, Rust J, Plautz Z, Kasturia S, Albornoz C, Wang W, Schmidt EF, Dannenfelser R, Tadych A, Brichta L, Barnea-Cramer A, Heintz N, Hof PR, Heiman M, Dolinski K, Flajolet M, Troyanskaya OG, Greengard P. 2020. Selective Neuronal Vulnerability in Alzheimer’s Disease: A Network-Based Analysis. Neuron 107:821–835.e12. 10.1016/j.neuron.2020.06.010

Saikia JM, Chavez-Martinez CL, Kim ND, Allibhoy S, Kim HJ, Simonyan L, Smadi S, Tsai KM, Romaus-Sanjurjo D, Jin Y, Zheng B. 2022. A Critical Role for DLK and LZK in Axonal Repair in the Mammalian Spinal Cord. J Neurosci 42:3716–3732. doi:10.1523/JNEUROSCI.2495-21.2022

Sanz E, Bean JC, Carey DP, Quintana A, McKnight GS. 2019. RiboTag: Ribosomal Tagging Strategy to Analyze Cell-Type-Specific mRNA Expression In Vivo. Curr Protoc Neurosci 88:e77. 10.1002/cpns.77

Sanz E, Yang L, Su T, Morris DR, McKnight GS, Amieux PS. 2009. Cell-type-specific isolation of ribosome-associated mRNA from complex tissues. Proc Natl Acad Sci U S A 106:13939–13944. doi:10.1073/pnas.0907143106

Schindelin J, Arganda-Carreras I, Frise E, Kaynig V, Longair M, Pietzsch T, Preibisch S, Rueden C, Saalfeld S, Schmid B, Tinevez J-Y, White DJ, Hartenstein V, Eliceiri K, Tomancak P, Cardona A. 2012. Fiji: an open-source platform for biological-image analysis. Nat Methods 9:676–682. doi:10.1038/nmeth.2019

Schmidt-Kastner R. 2015. Genomic approach to selective vulnerability of the hippocampus in brain ischemia-hypoxia. Neuroscience 309:259–279. doi:10.1016/j.neuroscience.2015.08.034

Shin JE, Cho Y, Beirowski B, Milbrandt J, Cavalli V, DiAntonio A. 2012. Dual Leucine Zipper Kinase Is Required for Retrograde Injury Signaling and Axonal Regeneration. Neuron 74:1015–1022. doi:10.1016/j.neuron.2012.04.028

Shin JE, Ha H, Cho Y, Kim YK, DiAntonio A. 2019. DLK regulates a distinctive transcriptional regeneration program after peripheral nerve injury. Neurobiol Dis 127:178–192. doi:10.1016/j.nbd.2019.02.001

Smith M-L, Auer RN, Siesjö BK. 1984. The density and distribution of ischemic brain injury in the rat following 2–10 min of forebrain ischemia. Acta Neuropathol 64:319–332. doi:10.1007/BF00690397

Somasundaram P, Farley MM, Rudy MA, Stefanoff DG, Shah M, Goli P, Heo J, Wang S, Tran NM, Watkins TA. 2023. Coordinated stimulation of axon regenerative and neurodegenerative transcriptional programs by Atf4 following optic nerve injury. BioRxiv doi:10.1101/2023.03.29.534798

Subramanian A, Tamayo P, Mootha VK, Mukherjee S, Ebert BL, Gillette MA, Paulovich A, Pomeroy SL, Golub TR, Lander ES, Mesirov JP. 2005. Gene set enrichment analysis: A knowledge-based approach for interpreting genome-wide expression profiles. Proc Natl Acad Sci 102:15545–15550. doi:10.1073/pnas.0506580102

Summers DW, Frey E, Walker LJ, Milbrandt J, DiAntonio A. 2020. DLK Activation Synergizes with Mitochondrial Dysfunction to Downregulate Axon Survival Factors and Promote SARM1-Dependent Axon Degeneration. Mol Neurobiol 57:1146–1158. doi:10.1007/s12035-019-01796-2

Tapias A, Lázaro D, Yin BK, Rasa SMM, Krepelova A, Sacramento EK, Grigaravicius P, Koch P, Kirkpatrick J, Ori A, Neri F, Wang ZQ. 2021. Hat cofactor trrap modulates microtubule dynamics via sp1 signaling to prevent neurodegeneration. Elife 10:1–64. doi:10.7554/eLife.61531

Tedeschi A, Bradke F. 2013. The DLK signalling pathway - A double-edged sword in neural development and regeneration. EMBO Rep 14:605–614. doi:10.1038/embor.2013.64

Thornburg-Suresh EJC, Richardson JE, Summers DW. 2023. The Stathmin-2 membrane-targeting domain is required for axon protection and regulated degradation by DLK signaling. J Biol Chem 299:1–16. doi:10.1016/j.jbc.2023.104861

Tortosa E, Sengupta Ghosh A, Li Q, Wong WR, Hinkle T, Sandoval W, Rose CM, Hoogenraad CC. 2022. Stress-induced vesicular assemblies of dual leucine zipper kinase are signaling hubs involved in kinase activation and neurodegeneration. EMBO J 41:1–20. doi:10.15252/embj.2021110155

Vervoort VS, Roselli S, Oshima RG, Pasquale EB. 2007. Splice variants and expression patterns of SHEP1, BCAR3 and NSP1, a gene family involved in integrin and receptor tyrosine kinase signaling. Gene 391:161–170. 10.1016/j.gene.2006.12.016

Viana da Silva S, Haberl MG, Gaur K, Patel R, Narayan G, Ledakis M, Fu ML, de Castro Vieira M, Koo EH, Leutgeb JK, Leutgeb S. 2024. Localized APP expression results in progressive network dysfunction by disorganizing spike timing. Neuron 112:124–140.e6. 10.1016/j.neuron.2023.10.001

Vornov JJ, Tasker RC, Coyle JT. 1991. Direct observation of the agonist-specific regional vulnerability to glutamate, NMDA, and kainate neurotoxicity in organotypic hippocampal cultures. Exp Neurol 114:11–22. 10.1016/0014-4886(91)90079-R

Wang F, Flanagan J, Su N, Wang LC, Bui S, Nielson A, Wu X, Vo HT, Ma XJ, Luo Y. 2012. RNAscope: A novel in situ RNA analysis platform for formalin-fixed, paraffin-embedded tissues. J Mol Diagnostics 14:22–29. doi:10.1016/j.jmoldx.2011.08.002

Watkins TA, Wang B, Huntwork-Rodriguez S, Yang J, Jiang Z, Eastham-Anderson J, Modrusan Z, Kaminker JS, Tessier-Lavigne M, Lewcock JW. 2013. DLK initiates a transcriptional program that couples apoptotic and regenerative responses to axonal injury. Proc Natl Acad Sci U S A 110:4039–4044. doi:10.1073/pnas.1211074110

Welsbie DS, Mitchell KL, Jaskula-Ranga V, Sluch VM, Yang Z, Kim J, Buehler E, Patel A, Martin SE, Zhang PW, Ge Y, Duan Y, Fuller J, Kim BJ, Hamed E, Chamling X, Lei L, Fraser ID, Ronai ZA, Berlinicke CA, Zack DJ. 2017. Enhanced Functional Genomic Screening Identifies Novel Mediators of Dual Leucine Zipper Kinase-Dependent Injury Signaling in Neurons. Neuron 94:1142–1154.e6. doi:10.1016/j.neuron.2017.06.008

West MJ, Coleman PD, Flood DG, Troncoso JC. 1994. Differences in the pattern of hippocampal neuronal loss in normal ageing and Alzheimer’s disease. Lancet 344:769– 772. 10.1016/S0140-6736(94)92338-8

Wickham H. 2016. Ggplot2: Elegant graphics for data analysis, 2nd ed. Cham, Switzerland: Springer International Publishing.

Wilde GJC, Pringle AK, Wright P, Iannotti F. 1997. Differential Vulnerability of the CA1 and CA3 Subfields of the Hippocampus to Superoxide and Hydroxyl Radicals In Vitro. J Neurochem 69:883–886. 10.1046/j.1471-4159.1997.69020883.x

Yoshihara Y, De Roo M, Muller D. 2009. Dendritic spine formation and stabilization. Curr Opin Neurobiol 19:146–153. 10.1016/j.conb.2009.05.013

Zeisel A, Hochgerner H, Lönnerberg P, Johnsson A, Memic F, van der Zwan J, Häring M, Braun E, Borm LE, La Manno G, Codeluppi S, Furlan A, Lee K, Skene N, Harris KD, Hjerling-Leffler J, Arenas E, Ernfors P, Marklund U, Linnarsson S. 2018. Molecular Architecture of the Mouse Nervous System. Cell 174:999–1014.e22. 10.1016/j.cell.2018.06.021

