## Supplemental figures and Table S1 for "Translatome analysis reveals cellular network in DLK-dependent hippocampal glutamatergic neuron degeneration"

**Supplemental methods**

HIPPOCAMPAL SPATIAL EXPRESSION ANALYSIS

False color expression images from the Allen Mouse Brain Atlas were used for evaluating expression pattern, and numbers were assigned based on color in dorsal hippocampus (Red=3, Yellow=2, Blue/Green=1, No=0). When intensity varied across sections or intensity was in-between two categories, preference was given to depicting general patterns of relative expression over absolute signal. When *in situ* data was not available, or expression patterns were unclear, we used additional transcriptomic data to assess spatial expression (Habib et al., 2016; Zeisel et al., 2018), and values were chosen to reflect relative expression. Generally, the following scale was used for Habib et al., 2016 data through the Single Cell Portal: 0 if next to no signal, 1 if expression in some cells, but average was still zero, 2 if quartile 3 value in violin plot is >0, 3 if higher average signal, again values were chosen to reflect relative expression. Genes were categorized as enriched in a region/s if one or two regions show higher values than another region. If two regions show different expression levels but are two levels above third region, the gene is considered as enriched in both (i.e., CA1=2, CA3=3, DG=0, considered as CA1, CA3 enriched). If only one level above other regions, the gene is enriched only in the region with strongest expression (i.e., CA1=3, CA3=2, DG=1, considered as CA1 enriched). Most expressed elsewhere in hippocampus used when the strongest expression is found in another region/cell type, and other descriptions don't explain where most of the signal is.

STMN2/STMN4 ANTIBODY SPECIFICITY

Given their highly similar protein size and sequences, we wanted to evaluate STMN2 and STMN4 antibody specificity. We used two antibodies for each. STMN2 antibodies were a mouse monoclonal anti-STMN2 (R&D Systems, MAB6930) and a rabbit polyclonal anti-STMN2 (Proteintech, 10586-1-AP). STMN4 antibodies were a mouse monoclonal anti-STMN4 (Santa Cruz, Sc-376936) and a rabbit polyclonal anti-STMN4 (Proteintech, 12027-1-AP). We tested the specificity of STMN2 and STMN4 antibodies by co-staining for STMN2 and STMN4, or with two separate STMN2 or STMN4 antibodies. In each case, antibody signal overlapped in the cell soma, presumably at the Golgi, as well as larger puncta elsewhere, with some overlapping small puncta, and some non-overlapping small puncta (Fig.S14A). Overlapping and non-overlapping signal can also be visualized by plotting intensity of signal along the neurite. By western blot STMN2 and STMN4 are highly similar at the protein level, with similar size proteins, though the STMN4 antibodies tested display a larger MW band specific to STMN4, suggesting some specificity. The STMN2 antibodies also occasionally recognized a smaller MW band only recognized with one of the STMN4 antibodies (Fig.S14B). Furthermore, while STMN4 protein levels increased relative to β-actin in mice with increased DLK, STMN2 protein levels did not show significant increases. These different expression patterns further validate some degree of specificity with these antibodies. Based on our analysis, the STMN4 Santa Cruz antibody (Sc-376936) may be more specific to STMN4 than the STMN4 Proteintech antibody (12027-1-AP) though it appears less sensitive. The STMN2 antibodies show strongest overlap of puncta and similar MW proteins, thus we were unable to detect differences in specificity. Whether the antibodies may also detect some of the same isoforms is not clear without further analysis.

**Supplemental Figure 1. Additional evidence for expression levels of DLK and effects on hippocampal morphology at 1 year of age**

(A) Confocal images of *Dlk* and *Vglut1* RNAscope analysis in hippocampal glutamatergic neurons at P15. Scale bar 1000μm, 10μm zoomed ROI. (B) Western blot of DLK and β-actin from protein extracts of hippocampal tissue of genotype indicated (age P60, each lane representing individual mice, N=3 mice/genotype). Arrow points to faint band of lower molecular weight in DLK(cKO) that is visible only under longer exposure, which may represent N-terminal truncated DLK produced using an alternative start codon (see C1). (C) IGV visual representation of RiboTag reads of *Dlk* in *Vglut1*$^{Cre/+}$ and *Vglut1*$^{Cre/+}$*;DLK(cKO)*$^{fl/fl}$, showing that mRNA for the floxed exon is reduced to ~1/3 level of control, while mRNA for other exons remained at a similar level as control. Dark blue illustration shows *Dlk* exon-intron structure, with pink triangles denoting loxP sites. Height of reads (y-axis) in gray or blue represents number of reads for the respective sequence. (C1) Enlarged view of the dashed box in C, corresponding to the floxed exon and exons encoding the kinase domain shown in orange with red marking ATP binding site and green marking DLK palmitoylation site. Start ATG shown in bold, and candidate downstream alternative start ATGs labeled with arrows. (D) Illustration of DLK overexpression transgene. (E,F) Confocal z-stack images of NeuN immunostaining and DAPI of coronal sections of dorsal hippocampus in mice of genotype indicated, with enlarged view of CA1, CA3, and DG (dashed boxes, respectively). ~1 year old control and *Vglut1*$^{Cre/+}$*;DLK(cKO)*$^{fl/fl}$ mice for (E); 44-46 weeks old, control and *Vglut1*$^{Cre/+}$*;H11-DLK*$^{iOE/+}$ mice for (F). Scale bar hippocampus 1000μm, enlarged view 50μm. (G,H,I) Quantification of cross-sectional area from CA1, CA3, or DG in dorsal hippocampus, respectively (as outlined in S2A,C). Data points represent individual mice, averaged across 3 sections per mouse, N≥3 mice/genotype. Statistics: One way ANOVA with Dunnett's multiple comparison test. ns, not significant; **** P<0.0001.

**Supplemental Figure 2. Hemibrain images across timepoints.**

Shown are lower magnification confocal images of NeuN staining of coronal sections of mice of genotype and age indicated. (A) ~1 year old control and *Vglut1*$^{Cre/+}$*;DLK(cKO)*$^{fl/fl}$ animals. (B) P10, P15, and P60 control and *Vglut1*$^{Cre/+}$*;H11-DLK*$^{iOE/+}$ animals. P60 tissue sections were stained at different time from those at P10, P15. (C) ~44-46 week old control and *Vglut1*$^{Cre/+}$*;H11-DLK*$^{iOE/+}$ animals. Arrows point to cortical thinning and ventricle expansion observed in some animals. Scale bar: 1000μm.

**Supplemental Figure 3. Evidence for induced DLK expression visualized at RNA and protein levels.**

(A) Confocal z-stack images of *Vglut1* and *Dlk* mRNAs at P15 in control and $Vglut1^{Cre/};H11\text{-}DLK^{iOE/+}$ mice. Scale bar: 1000μm. Inset shows single slice image of CA1 neurons, with dashed line showing individual nuclei as used for quantification. Scale bar: 10μm. (B) Quantification of RNAscope puncta shown as ratio of *Dlk* to *Vglut1* within individual CA1 neuron nuclei. Data points represent individual mice, averaged across 3 sections per mouse, ≥50 cells per genotype, N=5, 3 mice. Statistics: Mann-Whitney U test. **** P<0.0001. (C) Confocal z-stack images of control and $Vglut1^{Cre/+};H11\text{-}DLK^{iOE/+}$ mice immunostained for DLK protein (P15, P60, ~1 year) or no primary or LZK antibody control (P15) Dashed boxes are enlarged to the right. Individual strata are labeled in CA1, CA3; stratum pyramidale (SP), stratum radiatum (SR), stratum oriens (SO), stratum lacunosum-moleculare (SLM), stratum lucidum (SL). Scale bar 500μm in hippocampi, 100μm in CA1 and CA3. (D,E) Quantification of DLK mean fluorescence intensity from all layers in CA1, CA3, and DG at P10 (D) and P15 (E). N=4 mice/genotype for each timepoint. Statistics: Unpaired t-test. ns, not significant; * P<0.05.

**Supplemental Figure 4. Regional vulnerability observed with increased DLK expression.**

(A) Confocal z-stack images of NeuN immunostaining in ventral hippocampus in control and $Vglut1^{Cre/+};H11\text{-}DLK^{iOE/+}$ mice (P60). Dashed line regions are enlarged to the right, showing dorsal (posterior) CA1 neurons (top) with some thinning of pyramidal layer in mice with increased DLK, and ventral CA1 neurons (bottom) appearing similar between genotypes. Scale bar: 1,000μm in hippocampi; 100μm in CA1 layer. (B) Quantification of pyramidal layer thickness across CA1, with CA1 dorsal (anterior) data from Figure 1B. Data points represent individual mice, averaged across 2-3 sections per mouse. Statistics: Unpaired t-test. ns, not significant; **** P<0.0001. (C) Confocal z-stack images of p-c-Jun immunostaining in dorsal and ventral hippocampus of mice of indicated genotype at P60. Dashed line regions are enlarged to the right. Scale bar: 1,000μm in hippocampi; 100μm in CA1 layer.

**Supplemental Figure 5. Additional evidence for DLK(iOE) induced hippocampal neuron death.**

(A) Confocal z-stack imaging of GFAP immunostaining of hippocampus in mice of genotype indicated at P15 and P60, with enlarged view of the dashed boxes for CA1, CA3 shown below. Scale bar: 500μm in hippocampi, 100μm in CA1 and CA3. (B-D) Mean fluorescent intensity of GFAP in CA1, CA3, and DG regions (boxed area in A), respectively at P15. N= 5, 4 mice, quantified from 3 sections per mouse. Statistics: Unpaired t-test. ns, not significant; * P<0.05. P15, P60 GFAP animals imaged under separate conditions due to staining at different times. (E) Confocal z-stack images of IBA1 immunostaining of hippocampus in mice of genotype indicated at P15 and P60. Boxed microglia for CA1 and CA3 are enlarged below. Scale bar 100μm in CA1 and CA3 top, 10μm in individual microglia bottom. (F) Confocal single-slice images showing TUNEL positive CA1 pyramidal neuron in P15 *Vglut1*$^{Cre/+}$*;H11-DLK*$^{iOE/+}$ overlapping with tdTomato expressed from DLK(iOE) transgene and pyknotic nuclei (DAPI). Scale bar, 25μm. (G) Quantification of TUNEL positive neurons in CA1 pyramidal layer in genotype indicated. Data points represent individual mice, averaged across 3 sections per mouse, N=4 mice. Statistics: Unpaired t-test. ** P<0.01.

**Supplemental Figure 6. Evidence for RiboTag immunoprecipitated samples and additional analysis of genes showing differential dependence on DLK expression levels.**

(A) Western blot of HA-tagged Rpl22 immunoprecipitates. (B) qRT-PCR analysis of *Vglut1* (glutamatergic neurons), *Wfs1* (CA1), *Vgat* (inhibitory neurons), *Gfap* (astrocytes) shows expression of transcripts for glutamatergic neurons and depletion of non-glutamatergic neuron transcripts. N=3 biological replicates, data shown as expression fold change for marker genes in immunoprecipitated glutamatergic neuron RNAs relative to whole hippocampal RNAs, normalized to *Gapdh*. (C) Venn diagram showing overlap of statistically significant differentially expressed genes by RiboTag analysis of glutamatergic neurons in DLK(cKO) and DLK(iOE). (D,E) Heatmaps of the differentially expressed genes in glutamatergic neurons in DLK(cKO) and DLK(iOE). Columns represent expression levels in individual mice; rows represent individual genes, and the right-hand labels the RiboTag dataset where a given gene shows statistical significance. Data were normalized by row, with color keys shown above the heatmap. (F,G,H,I) Pie charts show distribution of differentially expressed genes detected in DLK(iOE) or DLK(cKO) in hippocampal neurons, based on *in situ* data in the Allen Mouse Brain Atlas (P56 mice). (J) SynGO sunburst plot shows enrichment of 10 differentially expressed genes from hippocampal glutamatergic neurons of DLK(cKO) mice, with color corresponding to significance. (K,L) GSEA enrichment plots show distribution of CA1 or CA3 genes from higher expression in WT to higher expression in DLK(cKO). Entire list of translated genes is represented by red to blue spectrum, genes expressed higher in WT in red, genes expressed higher in DLK(cKO) in blue (Supplemental excel file S1. WT vs DLKcKO DEGs). Vertical black lines (middle) represent genes in the respective gene set and where each lies along the spectrum from genes higher in WT to genes higher in DLK(cKO). Gene set for (K) CA1 genes (enriched in CAMK2 vs GRIK4 neurons); (L) CA3 genes (enriched in GRIK4 vs CAMK2 neurons) (Supplementary excel S3 CamK2 Grik4 enriched genes). Green line reflects running enrichment score (negative value representing CA1 genes tending to be more expressed in DLK(cKO)). Dashed box highlights region contributing to enrichment score. (K) Normalized enrichment score -1.89. False discovery rate q-value: 0.000; (L) Normalized enrichment score 1.42. False discovery rate q-value: 0.030.

**Supplemental Figure 7. Levels of c-Jun and p-c-Jun show dependency on expression levels of DLK.**

(A-B) Confocal z-stack images of c-Jun (A) and p-c-Jun (B) immunostaining in CA1, CA3, and DG in control and *Vglut1*$^{Cre/+}$*;H11-DLK*$^{iOE/+}$ mice (P10). Dashed lines in corresponding DAPI staining outline region used for quantification. Scale bar 50μm. Graphs below image panels show quantification of MFI of c-Jun (A) or p-c-Jun nuclei above intensity threshold per 100 μm of pyramidal or granule cell layer (B). Data points represent individual mice. 3 sections per mouse, N=4,5 mice for (A); N=4,4 mice for (B). Statistics: Unpaired t-test. ns, not significant, * P<0.05, ** P<0.01. (C-D) Confocal z-stack images of c-Jun (C) and p-c-Jun (D) immunostaining in CA1, CA3, and DG in control and *Vglut1*$^{Cre/+}$*;H11-DLK*$^{iOE/+}$ mice (P15). Dashed lines in corresponding DAPI staining outline region used for quantification. Scale bar 50μm. Graphs below image panels show quantification of MFI of c-Jun (C) or p-c-Jun nuclei above intensity threshold per 100 μm of pyramidal or granule cell layer (D). Data points represent individual mice. 3 sections per mouse, N=7,8 mice for (C); N=3,5 mice for (D). Statistics: Unpaired t-test. ns, not significant; * P<0.05; ** P<0.01; **** P<0.0001. (E) Confocal z-stack images of c-Jun (E) and p-c-Jun (F) immunostaining in CA1, CA3, and DG in control and *Vglut1*$^{Cre/+}$*;DLK(cKO)*$^{fl/fl}$ mice (P60). Dashed lines in corresponding DAPI staining outline region used for quantification. Scale bar 50μm. Graphs below image panels show quantification of mean fluorescence intensity (MFI) of c-Jun and p-c-Jun in pyramidal or granule cell layer of each region. Data points represent MFI of individual mice. 3 sections per mouse, N=4,4 mice for (E); N=6, 7 mice for (F). Statistics: Unpaired t-test. ns, not significant; * P<0.05.

**Supplemental Figure 8. Stathmin transcript abundance and western blot analysis of DLK, STMN4 in mice aged P10 to 1 yr.**

(A) RiboTag analysis of Stathmin family members, shown as transcripts per million (TPM). Differential expression analysis significance shown ($p_{adj}$). ns, not significant; **** P<0.0001. Stmn2 and Stmn4 appear to have comparable reads, however, this includes reads in a retained intron in Stmn2, which increases the reference gene length used for the TPM calculation for Stmn2. The significance of this intron retention may need further study. (B,C) Western blots of protein extracts from hippocampal tissue from P1, P8, P15, P60, and ~1yr old mice, blotted for DLK, Flag, STMN4, STMN2, and actin in *Vglut1*$^{Cre/+}$*;DLK(cKO)*$^{fl/fl}$ (B) and *Vglut1*$^{Cre/+;}$*H11-DLK*$^{iOE/+}$ (C), respectively. Larger molecular weight band of DLK in Vglut1Cre/+;H11-DLKiOE/+ would match the predicted molecular weight of DLK-T2A-tdTomato if T2A-peptide induced 'self-cleavage' due to ribosomal skipping is ineffective (Fig.S1D). (D,E) Relative DLK protein level normalized to actin and P1 control. N=3 mice/genotype. (F,G) Relative STMN4 protein level normalized to actin and P1 control. N=3 mice/genotype. Statistics: Two-way ANOVA with Sidak multiple comparisons test. ns, not significant; * P<0.05.

**Supplemental Figure 9. Additional evidence for *Stmn4* and microtubule expression in DLK(cKO) and DLK(iOE) in hippocampus.**

(A-D) Confocal single-slice images of *Stmn4* and *Vglut1* RNAscope stained sections from (A,B) CA3 and (C,D) DG in DLK(iOE) (A,C) and DLK(cKO) (B,D) mice at P15. Scale bar 10μm. (E) Confocal z-stack images of Tuj1 immunostaining in CA1, CA3, and DG of mice genotype indicated at P15. Scale bar 50μm. (F) Confocal z-stack images of MAP2 immunostaining of CA1 pyramidal layer from control and *Vglut1*$^{Cre/+}$*;H11-DLK*$^{iOE/+}$ mice at P60. Arrows point to apical dendrites with elevated signal. Arrowheads point to thin neurites with elevated signal. Scale bar 10μm. (G) Normalized MAP2 MFI after thresholding signal in SR (dashed outlines on images in F). N=9, 9 mice, 3 sections averaged per mouse. Statistics: Unpaired t-test. ns, not significant.

**Supplemental Figure 10. VGLUT1 pattern in dorsal CA1 in DLK(cKO) and DLK(iOE).**

(A) Confocal single-slice image of VGLUT1 immunostaining of CA1 SR in control and *Vglut1*$^{Cre/+}$*;DLK(cKO)*$^{fl/fl}$ mice at P60. Scale bar 5μm, inset scale bar 1μm. (B-C) Quantification of VGLUT1 puncta density and size. Data points represent averages from individual mice across 3 sections per mouse. N=5 control, and 8 *Vglut1*$^{Cre/+}$*;DLK(cKO)*$^{fl/fl}$ mice. (D) Confocal single-slice image of VGLUT1 immunostaining of CA1 SR in control and *Vglut1*$^{Cre/+}$*;H11-DLK*$^{iOE/+}$ mice at P15. Scale bar 5μm, inset scale bar 1μm. (E-F) Quantification of VGLUT1 puncta density (E) and size (F). Data points represent averages from individual mice across 3 sections per mouse, N=9 control, and 8 *Vglut1*$^{Cre/+}$*;H11-DLK*$^{iOE/+}$ mice. Statistics: unpaired t-test. ns, not significant; * P<0.05.

**Supplemental Figure 11. Analysis of Bassoon and Homer1 immunostaining in dorsal CA1 in P10 of DLK(iOE)**

(A) Confocal single-slice images of Bassoon and Homer1 immunostaining in CA1 SR of control and *Vglut1*$^{Cre/+}$*;H11-DLK*$^{iOE/+}$ mice at P10. Scale bars, 5μm in panel images, and 1μm in enlarged images. (B,C) Quantification of Bassoon and Homer1 puncta density, respectively. (D) Quantification of synapses displaying co-localization of Bassoon and Homer1. (E-F) Quantification from control and *Vglut1*$^{Cre/+}$*;H11-DLK*$^{iOE/+}$ mice for (E) Bassoon puncta size, (F) Homer1 puncta size. Data points represent average values per mouse from 3 sections. N=4 control, and 7 *Vglut1*$^{Cre/+}$*;H11-DLK*$^{iOE/+}$ mice. Statistics: unpaired t-test. ns, not significant; * P<0.05.

**Supplemental Figure 12. Analysis of Bassoon and Homer1 immunostaining in CA3 synapses of DLK(iOE) at P10 and P15.**

(A) Confocal single-slice images of Bassoon and Homer1 immunostaining in CA3 SR of control and *Vglut1*$^{Cre/+}$*;H11-DLK*$^{iOE/+}$ mice at P10. Scale bars, 5μm in panel images, and 1μm in enlarged images. (B,C) Quantification of Bassoon and Homer1 puncta density, respectively. (D) Quantification of synapses displaying co-localization of Bassoon and Homer1. (E,F) Quantification from control and *Vglut1*$^{Cre/+}$*;H11-DLK*$^{iOE/+}$ mice for (E) Bassoon puncta size, (F) Homer1 puncta size. Data points represent average values per mouse from 3 sections. N=4 control, and 7 *Vglut1*$^{Cre/+}$*;H11-DLK*$^{iOE/+}$ mice. Statistics: unpaired t-test. ns, not significant. (G)

Confocal single-slice images of Bassoon and Homer1 immunostaining in CA3 SL of control and *Vglut1*$^{Cre/+}$*;H11-DLK*$^{iOE/+}$ mice at P10. Scale bars, 5μm in panel images, and 1μm in enlarged images. (H,I) Quantification of Bassoon and Homer1 puncta density, respectively. (J) Quantification of synapses displaying co-localization of Bassoon and Homer1. (K,L) Quantification from control and *Vglut1*$^{Cre/+}$*;H11-DLK*$^{iOE/+}$ mice for (K) Bassoon puncta size, (L) Homer1 puncta size. Data points represent average values per mouse from 3 sections. N=4 control, and 7 *Vglut1*$^{Cre/+}$*;H11-DLK*$^{iOE/+}$ mice. Statistics: unpaired t-test. ns, not significant. (M) Confocal single-slice images of Bassoon and Homer1 immunostaining in CA3 SR of control and *Vglut1*$^{Cre/+}$*;H11-DLK*$^{iOE/+}$ mice at P15. Scale bars, 5μm in panel images, and 1μm in enlarged images. (N,O) Quantification of Bassoon and Homer1 puncta density, respectively. (P) Quantification of synapses displaying co-localization of Bassoon and Homer1. (Q,R) Quantification from control and *Vglut1*$^{Cre/+}$*;H11-DLK*$^{iOE/+}$ mice for (Q) Bassoon puncta size, (R) Homer1 puncta size. Data points represent average values per mouse from 3 sections. N=9 control, and 6 *Vglut1*$^{Cre/+}$*;H11-DLK*$^{iOE/+}$ mice. Statistics: unpaired t-test. ns, not significant. (S) Confocal single-slice images of Bassoon and Homer1 immunostaining in CA3 SL of control and *Vglut1*$^{Cre/+}$*;H11-DLK*$^{iOE/+}$ mice at P15. Scale bars, 5μm in panel images, and 1μm in enlarged images. (T,U) Quantification of Bassoon and Homer1 puncta density, respectively. (V) Quantification of synapses displaying co-localization of Bassoon and Homer1. (W,X) Quantification from control and *Vglut1*$^{Cre/+}$*;H11-DLK*$^{iOE/+}$ mice for (W) Bassoon puncta size, (X) Homer1 puncta size. Data points represent average values per mouse from 3 sections. N=9 control, and 6 *Vglut1*$^{Cre/+}$*;H11-DLK*$^{iOE/+}$ mice. Statistics: unpaired t-test. ns, not significant.

**Supplemental Figure 13. Analysis of Stathmins in primary cultured hippocampal neurons from DLK(cKO) and DLK(iOE).**

(A) Confocal z-stack images of neuron morphology at DIV2 from mice of genotype indicated, visualized by tdTomato from Rosa26-tdTomato. Neurons with indicated genotypes are labeled by tdTomato from Cre-dependent Rosa26-tdTomato generated from the following crosses: for control: *Vglut1*$^{Cre/+}$ X *Rosa26*$^{tdT/+}$; for DLKcKO: *Vglut1*$^{Cre/+}$*;DLK(cKO)*$^{fl/fl}$ X *Vglut1*$^{Cre/+}$*;DLK(cKO)*$^{fl/fl}$*;Rosa26*$^{tdT/+}$*;* for DLKiOE: *H11-DLK*$^{iOE/iOE}$ X *Vglut1* $^{Cre/+}$*;Rosa26*$^{tdT/+}$*.* Red arrowheads point to long processes considered as axons. Scale bar 100μm. (B) Co-immunostaining of DLK and STMN4 in DIV2 control cultured neuron growth cone shows non-overlapping puncta. (C) Confocal z-stack images of DLK and STMN2 co-immunostaining of

DIV2 primary hippocampal neurons from genotypes indicated; neurons are labeled with tdTomato from Rosa26-tdTomato. Scale bar 10μm. (D) Quantification of association between DLK level and STMN2 in cell soma. N=3 cultures/genotype, ≥45 cells/genotype. Spearman correlation r=0.4693. (E) Analysis of cell types in culture at DIV14 using Prox1 and Satb2 markers. Quantification from N=3 cultures/genotype, ≥200 cells/genotype. Statistics: Two-way ANOVA with Dunnett's multiple comparison test. ns, not significant.

**Supplemental Figure 14. Comparison of STMN2 and STMN4 antibodies**

(A) Confocal single-slice images of STMN2 and STMN4 immunostaining using two independent antibodies on control primary hippocampal neurons DIV3. Dashed boxes are enlarged below. Scale bar,10μm full cell, 1μm enlarged region. Line scan below reflects normalized intensity, with asterisks reflecting overlapping peaks. (B) Western blots of protein extracts from P15 hippocampal tissue of genotype indicated for STMN2 and STMN4 antibodies. Lanes 1-3 are littermate controls (+) for lanes 4-6 *Vglut1*$^{Cre/+;}$*H11-DLK*$^{iOE/+}$*,* referred to as "Tg"; and lanes 7-9 are littermate controls (+) for lanes 10-12 *Vglut1*$^{Cre/+}$*;DLK(cKO)*$^{fl/fl}$ , referred to as "-". (C) Quantification of STMN2 or STMN4 protein level normalized to actin. N=3 mice/genotype. Statistics: One way ANOVA with Sidak's multiple comparison test. ns, not significant; * P<0.05.

### Supplementary Figure 1

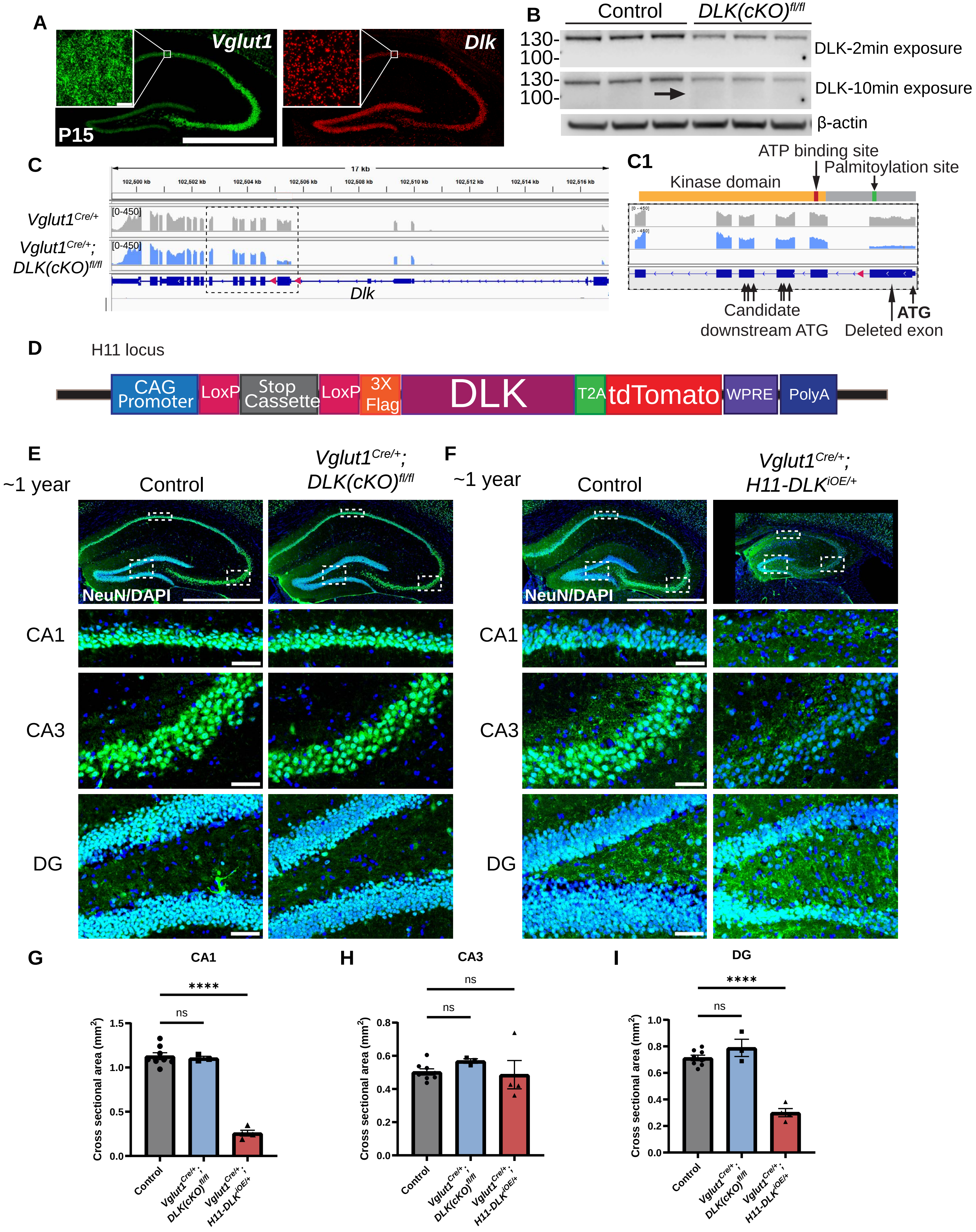

### Supplementary Figure 2

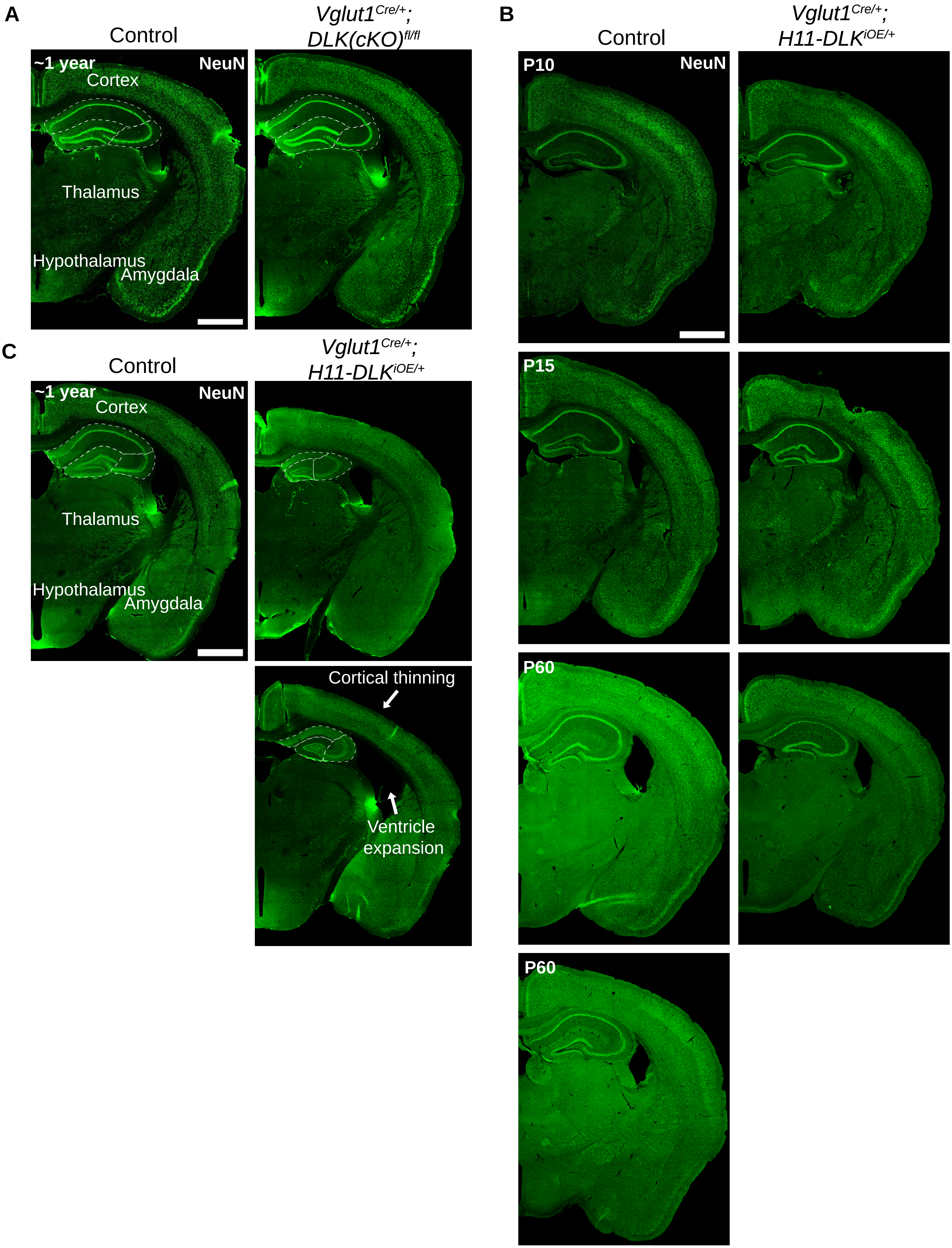

### Supplementary Figure 3

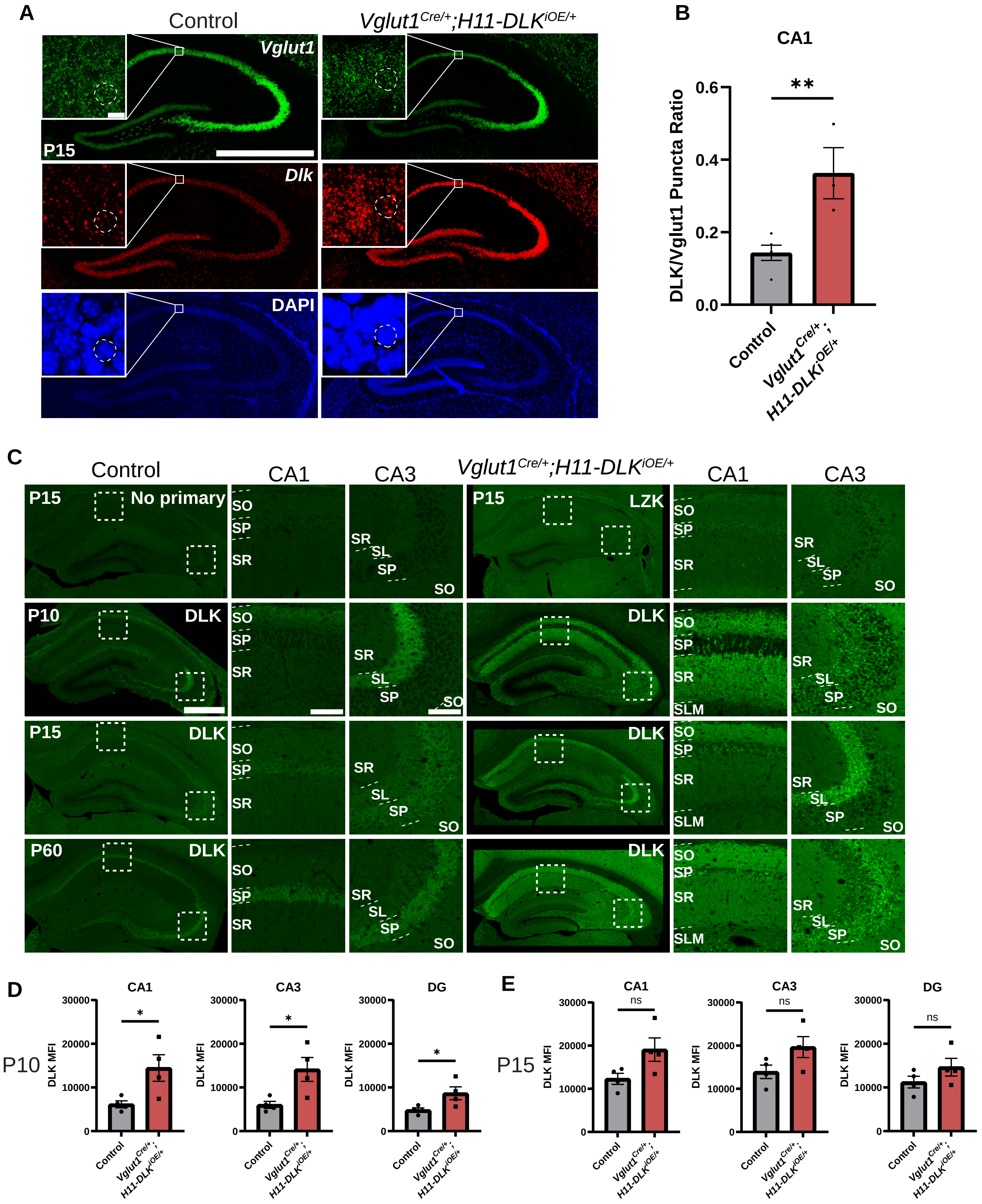

### Supplementary Figure 4

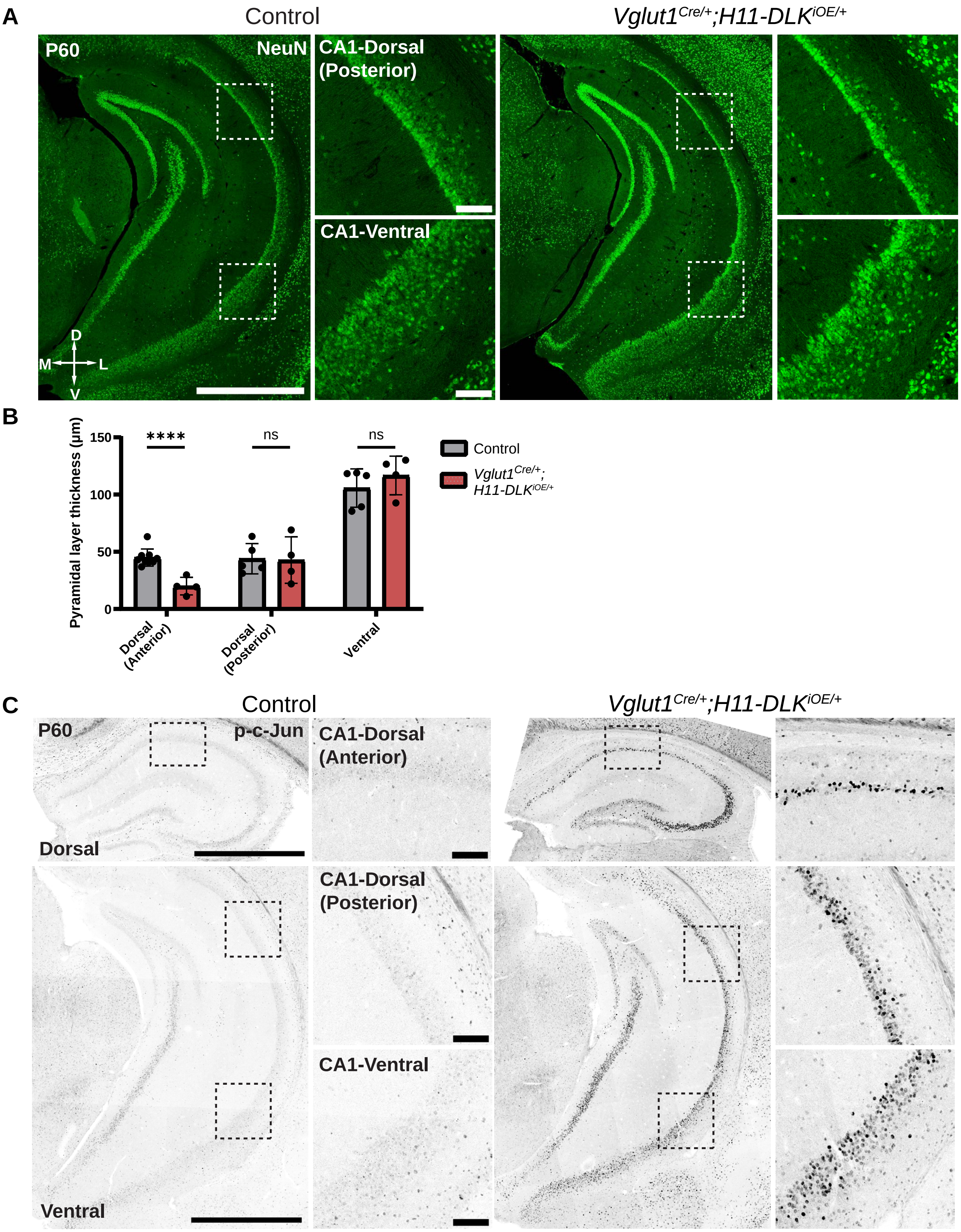

### Supplementary Figure 5

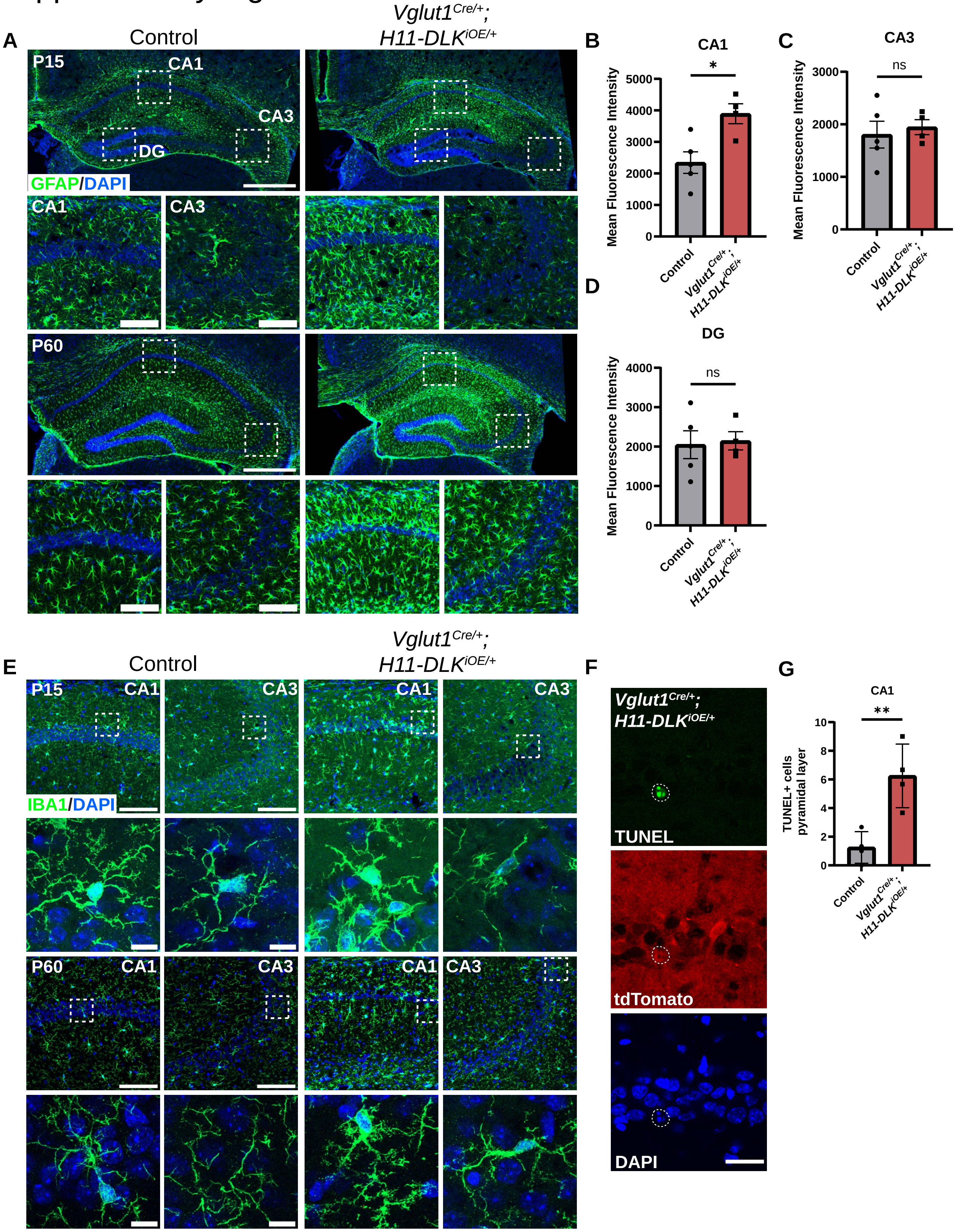

### Supplemental Figure 6

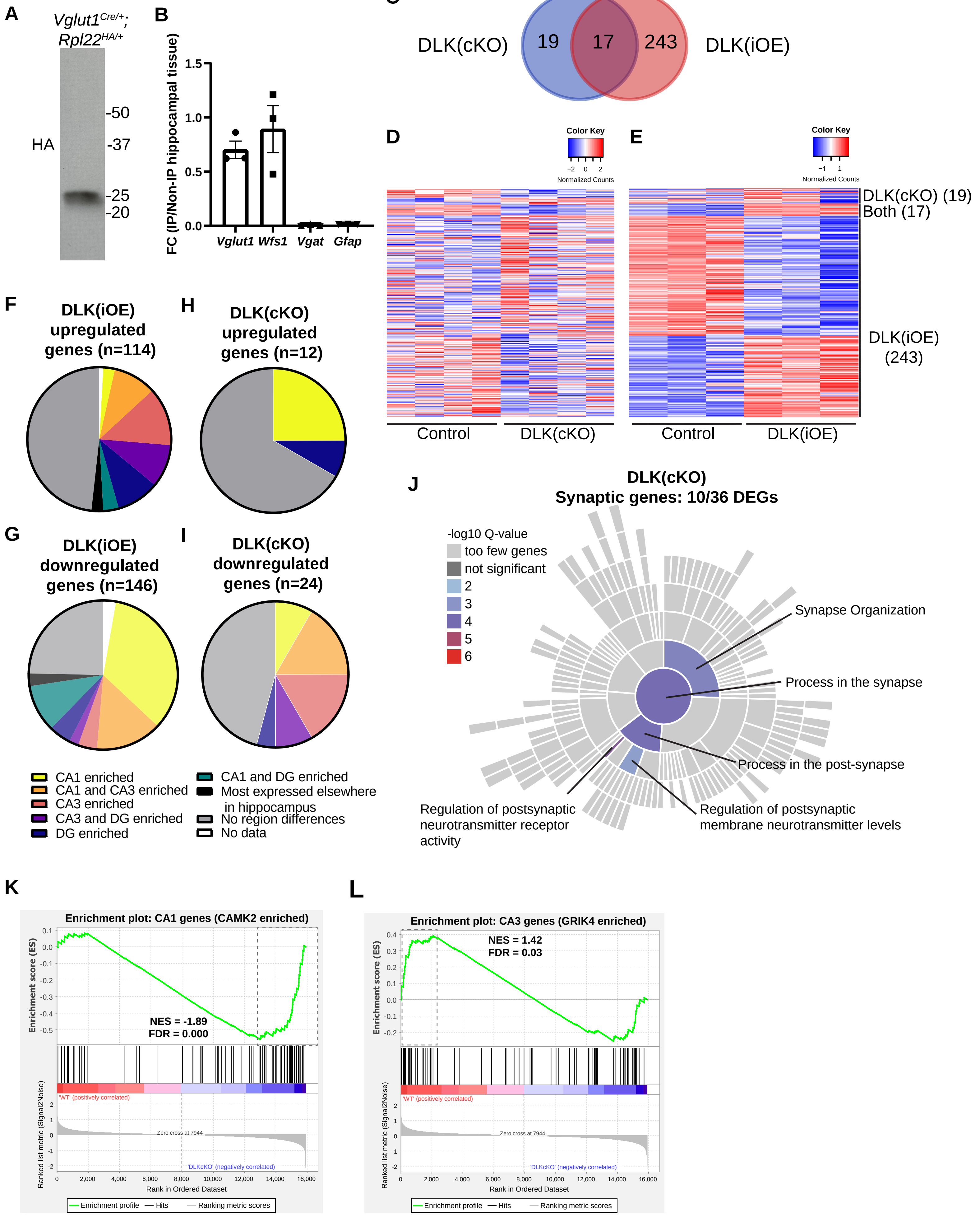

### Supplemental Figure 7

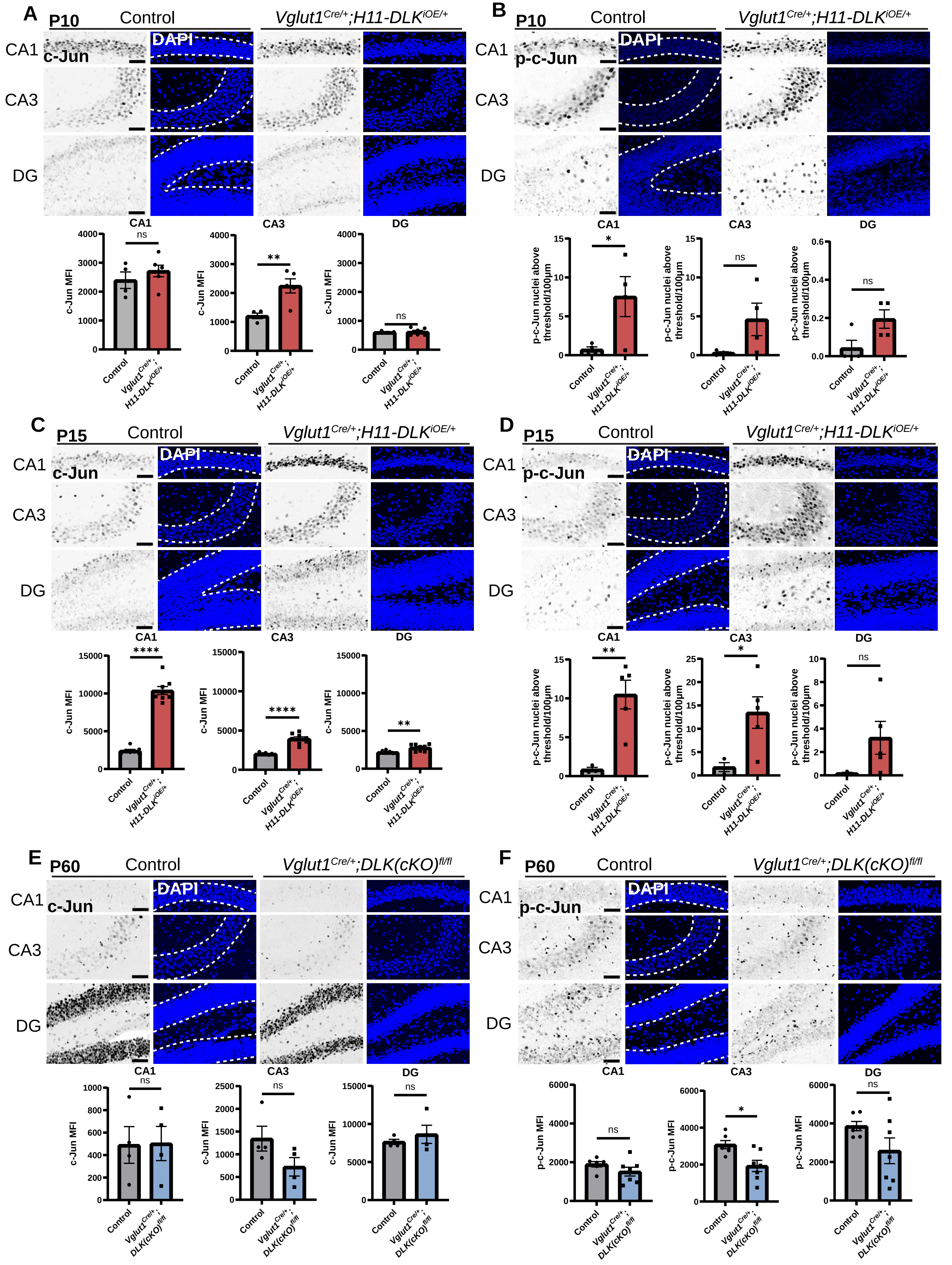

Supplemental Figure 8

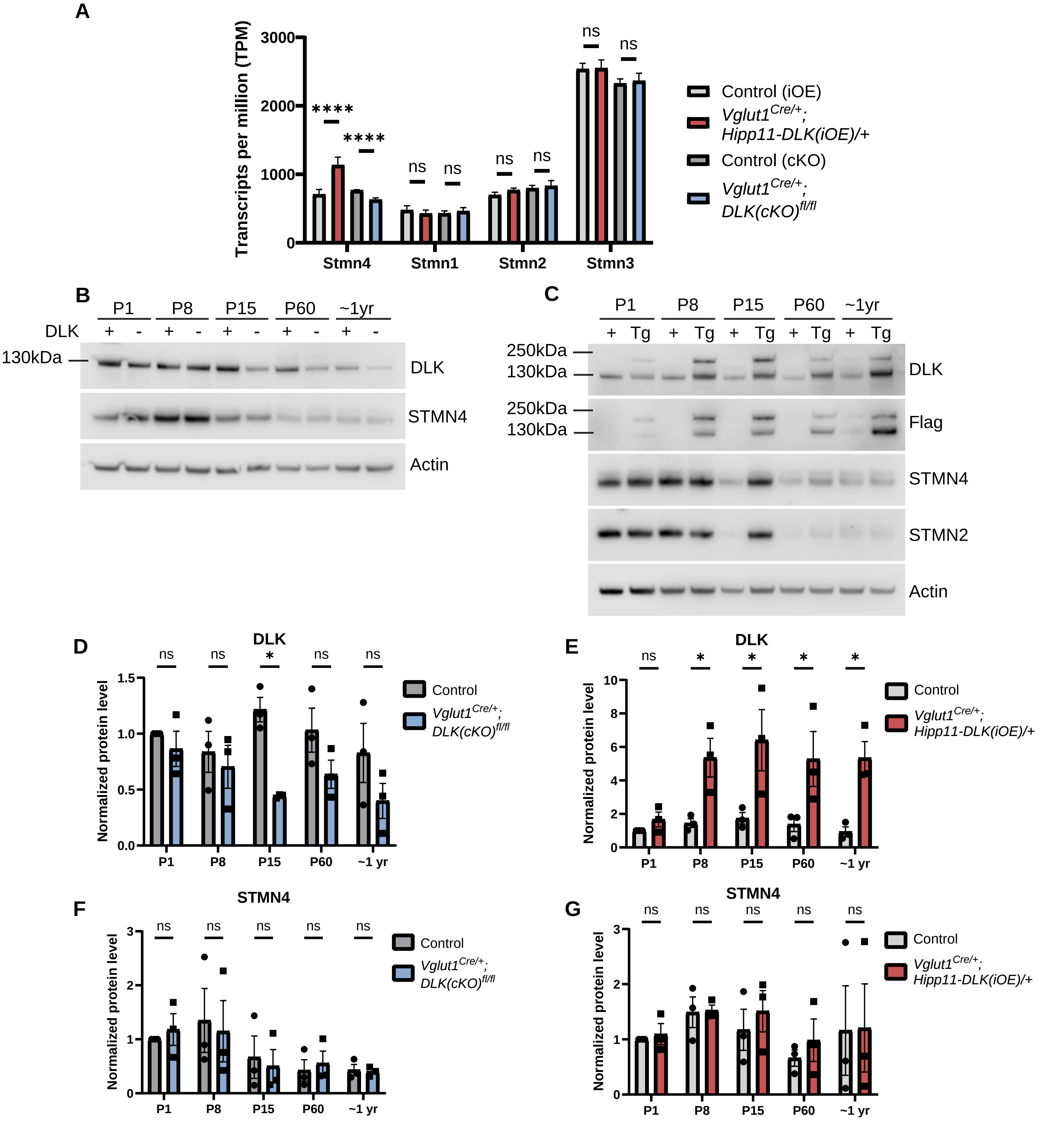

### Supplemental Figure 9

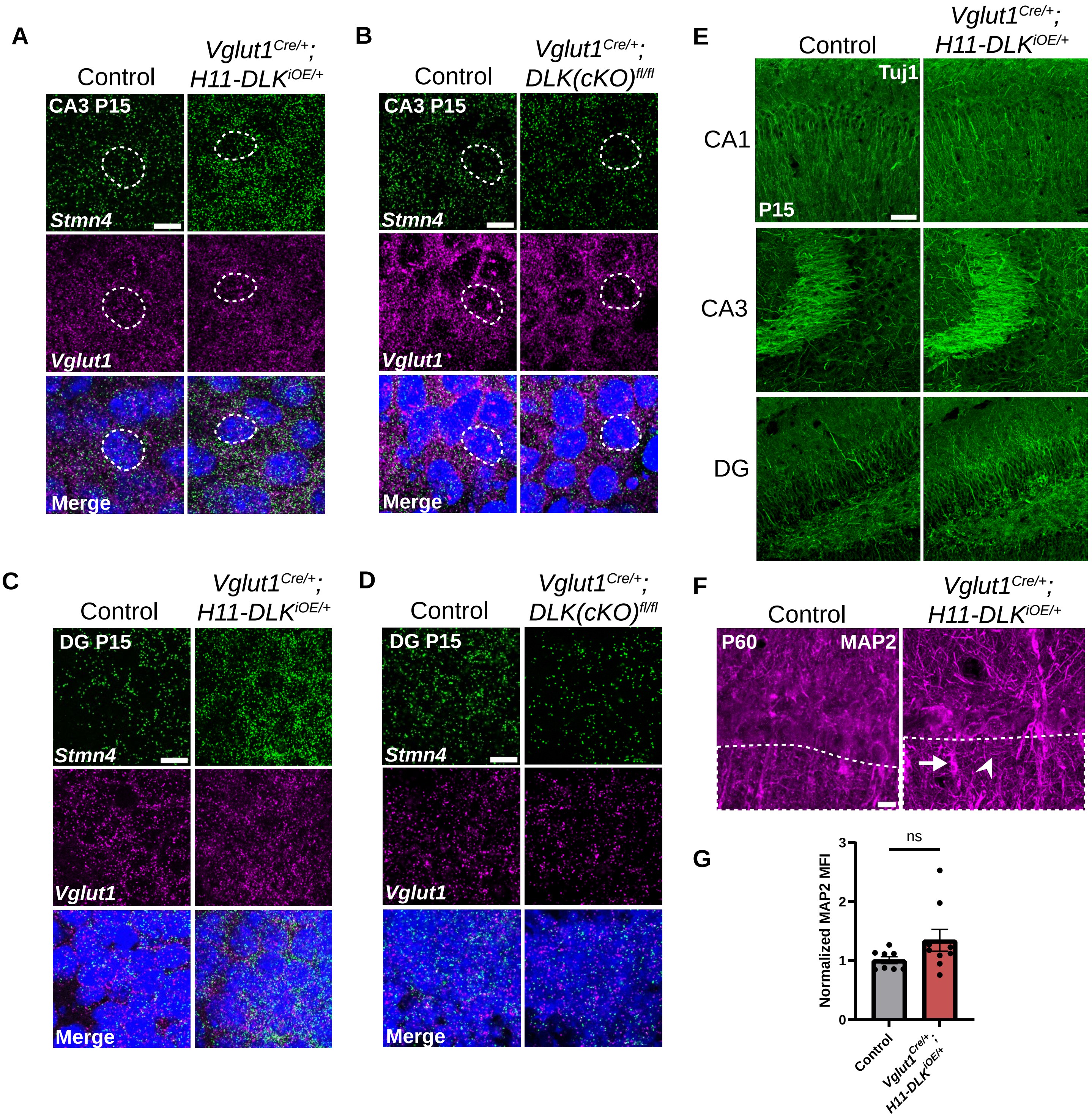

### Supplemental Figure 10

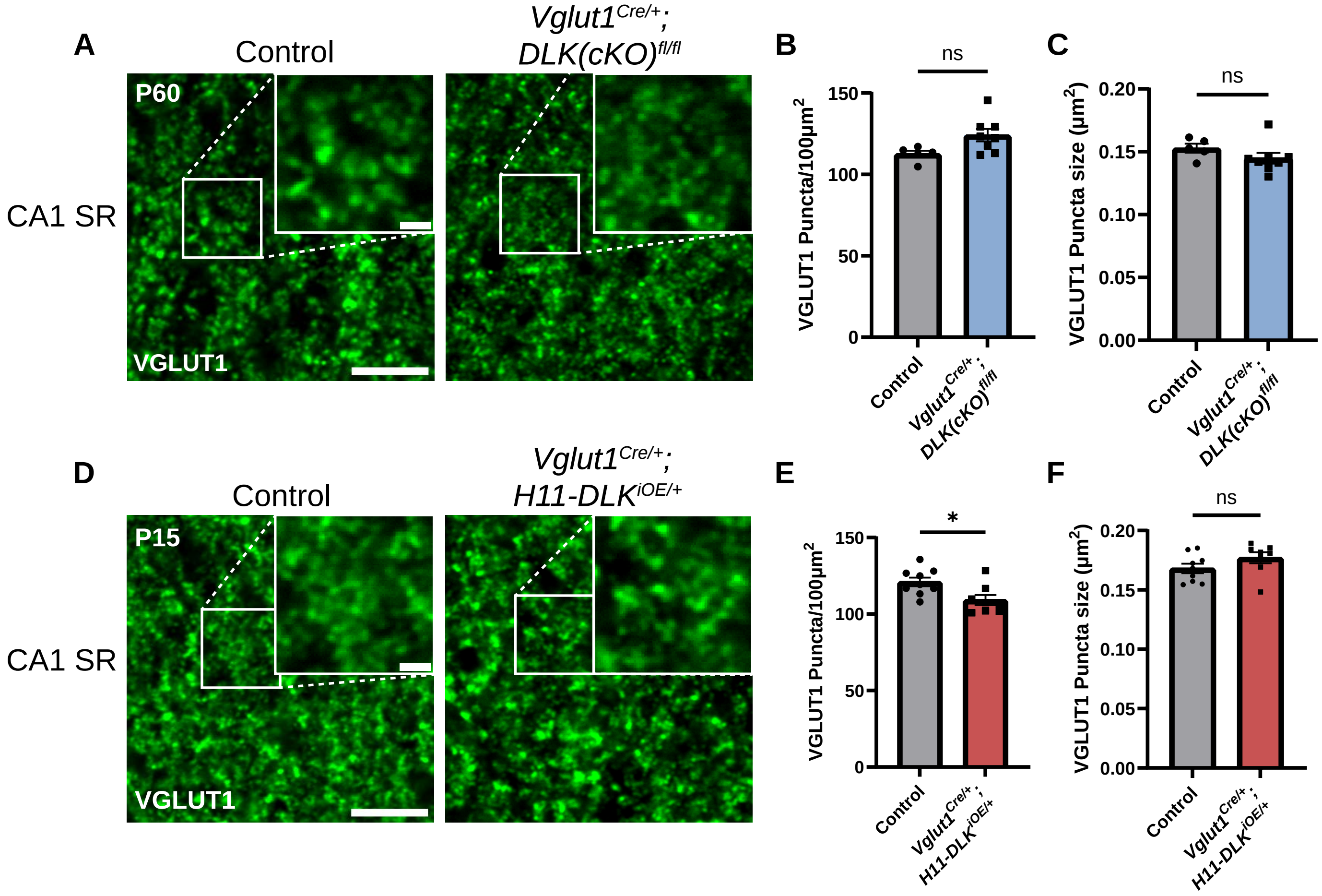

### Supplemental Figure 11

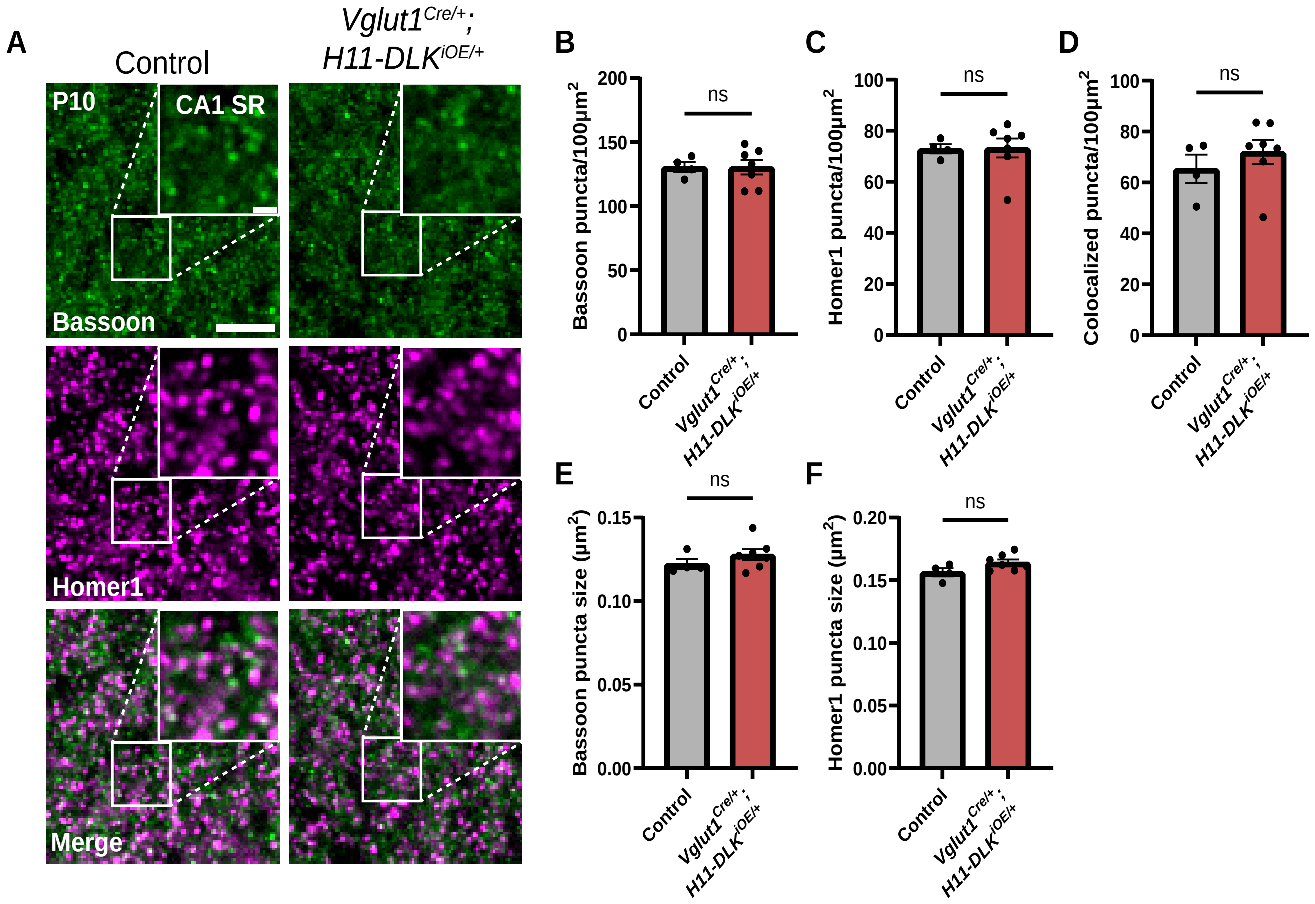

### Supplemental Figure 12

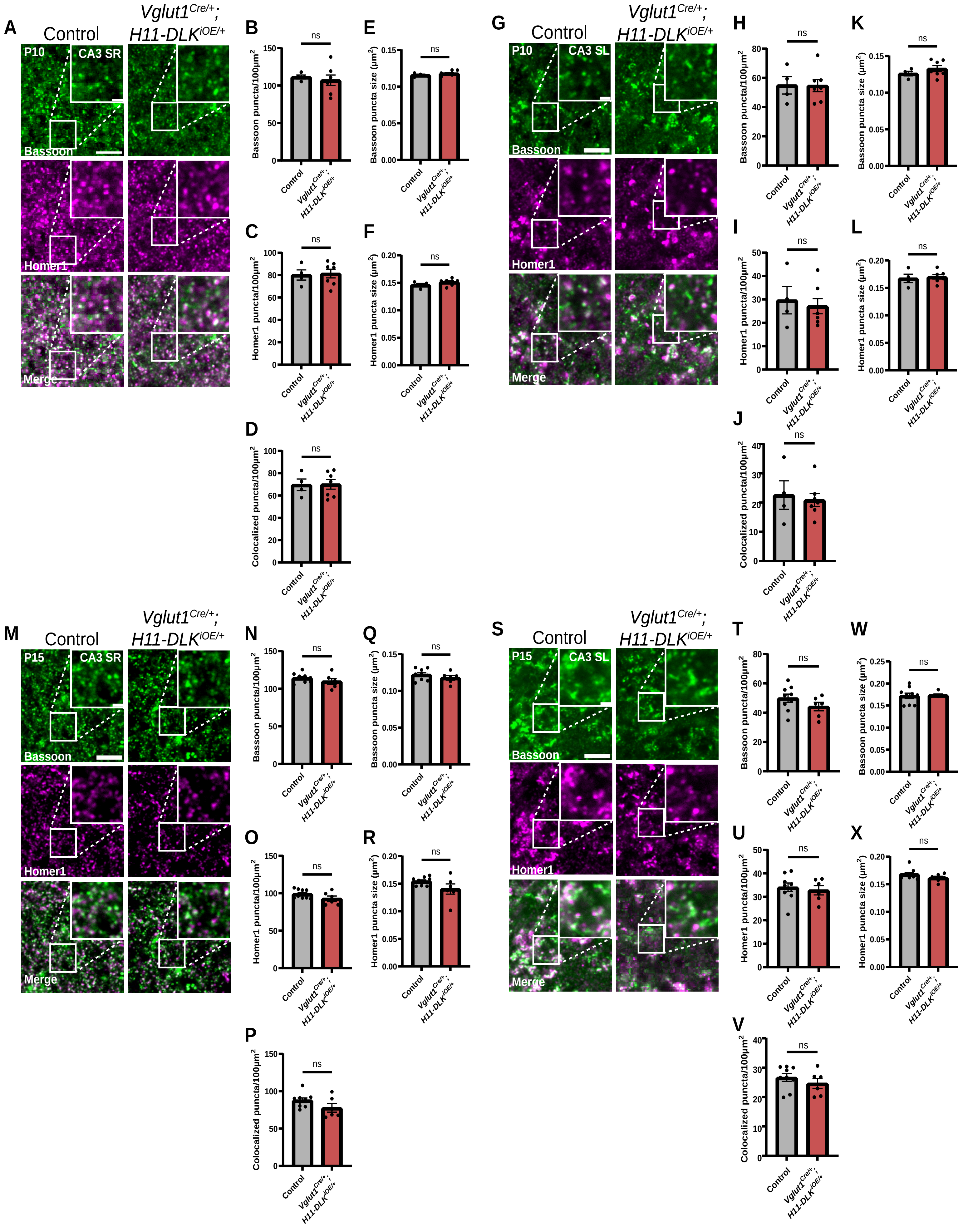

### Supplemental Figure 13

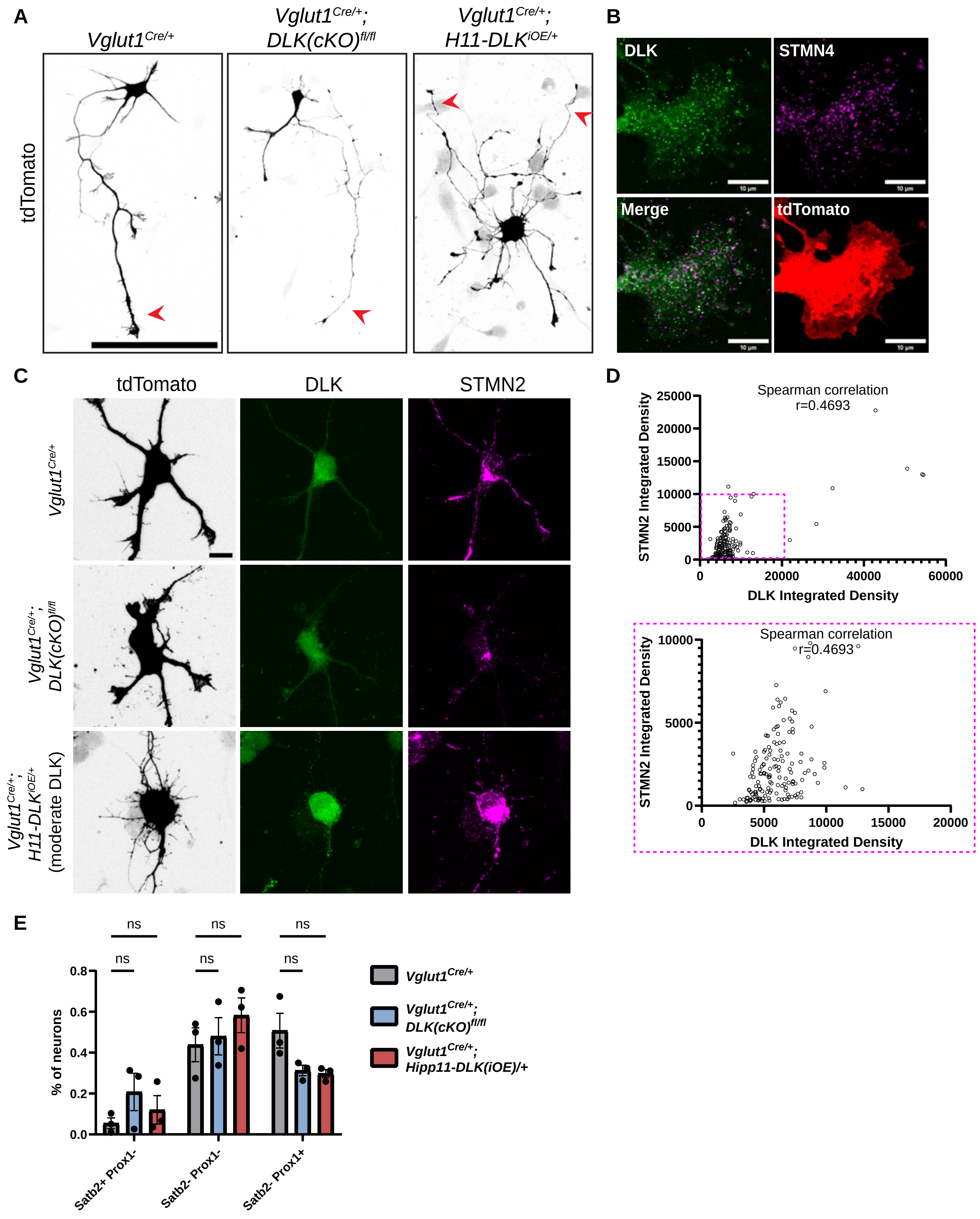

### Supplemental Figure 14

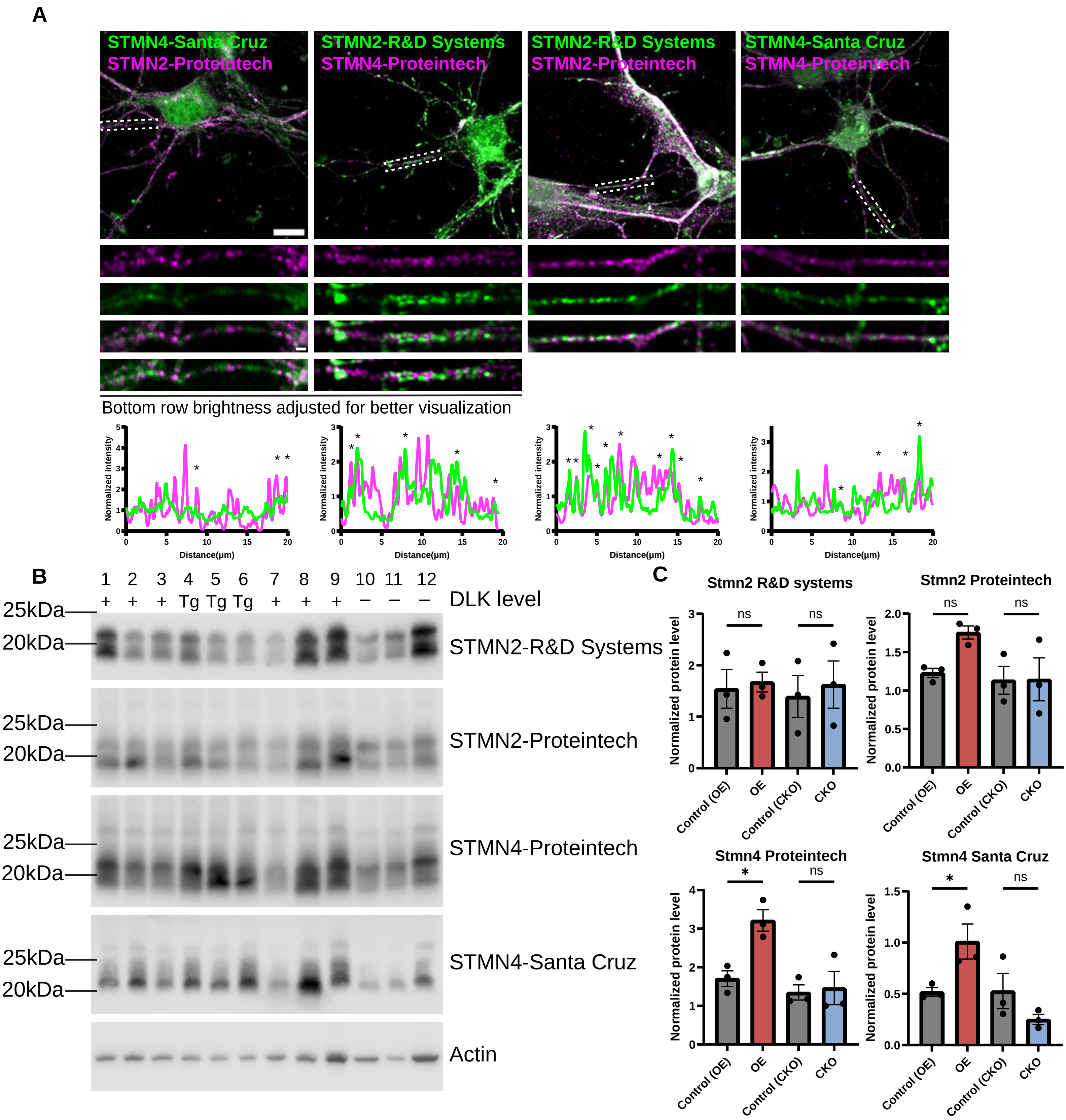

**Table S1. Primers**

| Primers for genotyping | | | |
|---|---|---|---|
| Reagent or resource | Source | Identifier | Additional information |
| Hipp11 WT-Fw: TGGAGGAGGACAAACTGGTCA | Li et al., 2021 | YJ12520 | 323bp (*Dlk* Wild-type) ~500bp ($Dlk^{OE}$ mutant) |
| Hipp11 WT-Re: TTCCCTTTCTGCTTCATCTTGC | Li et al., 2021 | YJ12521 | |
| CAG-Re: CATATATGGGCTATGAACTAATGACCCCGT | Li et al., 2021 | YJ12522 | |
| $DLK^{fx/fx}$-Fw: GATGATTGCTAGTCATGGAGTAGTAGG | Li et al., 2021 | YJ12523 | 350bp (*Dlk* Wild-type) 500bp ($Dlk^{fl/fl}$ mutant) |
| $DLK^{fx/fx}$-Re: GGTGGTGTTATCATAGTTCCATCATG | Li et al., 2021 | YJ12524 | |
| RiboTag Fw: GGGAGGCTTGCTGGATATG | Sanz et al., 2009 | | 260bp (WT) 290bp (floxed allele) |
| RiboTag Re: TTTCCAGACACAGGCTAAGTA | Sanz et al., 2009 | | |
| Vglut1-cre Re: CCCTAGGAATGCTCGTCA AG | The Jackson Laboratory | 12231 | 218bp (WT) 344bp (mutant) |
| Vglut1-cre Fw: ATGAGCGAGGAGAAGTGTGG | The Jackson Laboratory | 17904 | |
| Vglut1-cre Re: GTGGAAGTCCTGGAAACTGC | The Jackson Laboratory | 17905 | |
| Cre Fw: AGAACCTGAAGATGTTCGCG | | | ~330bp (mutant) No band WT |
| Cre Re: GGCTATACGTAACAGGGTGT | | | |
| Rosa tdTomato Re: GGCATTAAAGCAGCGTATCC | The Jackson Laboratory | oIMR9103 | 196bp (mutant) 297bp (WT) |
| Rosa tdTomato Fw: CTGTTCCTGTACGGCATGG | The Jackson Laboratory | oIMR9105 | |
| WT tdTomato Fw: AAGGGAGCTGCAGRGGAGTA | The Jackson Laboratory | oIMR9020 | |
| WT tdTomato Re: CCGAAAATGTGTGGGAAGTC | The Jackson Laboratory | oIMR9021 | |

| Primers for qRT-PCR | |
|---|---|
| Reagent or resource | Source |
| Gapdh-Fw: GCTTGTCATCAACGGGAAG | Furlanis et al., 2019 |
| Gapdh-Re: TTGTCATATTTCTCGTGGTTCA | Furlanis et al., 2019 |
| Vgat-Fw: CGTGACAAATGCCATTCAG | Furlanis et al., 2019 |
| Vgat-Re: AAGATGATGAGGAACAACCC | Furlanis et al., 2019 |
| Vglut1-Fw: ACCCTGTTACGAAGTTTAACAC | Furlanis et al., 2019 |
| Vglut1-Re: CAGGTAGAAGGTCCAGCTG | Furlanis et al., 2019 |
| Wsf1-Fw: CATCATTCCCACCAACCTG | Furlanis et al., 2019 |
| Wsf1-Re: TACTTCACCACCTTCTGGC | Furlanis et al., 2019 |
| Gfap-Fw: CTCGTGTGGATTTGGAGAG | Furlanis et al., 2019 |
| Gfap-Re: AGTTCTCGAACTTCCTCCT | Furlanis et al., 2019 |
